# Potent and Selective IL-4 Inhibitors with Anti-Tumor Activity

**DOI:** 10.64898/2026.02.01.702098

**Authors:** Raavi, Imron Chaudhry, Daniel F. Sheehy, Sean P. Quinnell, Chandler Ruping, Jacob Lee, Sha Hu, Haoyi Hou, Pinghua Liu, Arturo J. Vegas

## Abstract

Interleukin-4 (IL-4) is an important immunoregulatory cytokine involved in T-cell maturation, B-cell activation, and macrophage polarization. Dysregulated IL-4 signaling contributes to several immune-mediated diseases such as cancer, allergic inflammation, and autoimmunity. The clinical use and indication expansion of the anti-IL-4Rα antibody dupilumab has made IL-4 signaling an attractive target for therapeutic modulation. We previously discovered a first-in-class small molecule inhibitor to the soluble cytokine IL-4, which we named Nico-52, that inhibits the soluble IL-4 cytokine with single-digit micromolar potency. Here, we determined structure-activity relationships around the Nico-52 scaffold that impact potency and selectivity and evaluated the *in vivo* anti-tumor potential of small molecule IL-4 inhibition. Improved analogs featured structural changes to the p-fluorophenyl group ranging from submicromolar to double-digit nanomolar potency. Our two most potent analogs showed selective binding to IL-4 over other related cytokines in thermal shift assays and more potent inhibition of IL-4 over IL-13 in a HEK Blue IL-4/IL-13 reporter assay. We further established that our lead analogs inhibit both type I and type II IL-4 receptor signaling. Nico-52 and an optimized lead analog exhibited favorable *in vitro* ADME/T properties, such as high stability and low cytotoxicity. Furthermore, Nico-52 and a lead analog were investigated for their tumor suppressive effects in syngeneic murine tumor models, where small-molecule IL-4 inhibition yielded significant tumor inhibition, shifted macrophage polarization, and our optimized lead analog improved animal survival. These studies show the promise of small-molecule cytokine inhibitors for IL-4 mediated processes of disease.

## INTRODUCTION

Interleukin-4 (IL-4) is an important immunoregulator of inflammation and plays a role in the progression of auto-immunity, allergic reactions, arthritis, and cancer^1^. In asthma, IL-4 induces airway inflammation, obstruction, and hyperresponsiveness^2,3^ while in cancer IL-4 activity is linked to promoting tumor progression, immunosuppression, and increasing tumor resistance to apoptosis. IL-4 engages with two types of heterodimeric cell membrane receptors, type I (IL-4Rα & γc) and type II (IL-4Rα and the IL-13Rα1), for downstream JAK1/2/3-TyK2 mediated phosphorylation by ligand-induced receptor dimerization.^4–6^ The expression of these IL-4 receptors is determined primarily by tissue, with type I receptor complexes being restricted mainly to hematopoietic lineage cells whereas type II expression is more widespread. The dysregulation of cellular signaling mediated by these receptors is affiliated with numerous inflammatory conditions.^7,8^

Due to the wide range of conditions associated with IL-4-mediated inflammation, there has been considerable clinical and commercial interest in inhibiting IL-4 and related signaling proteins. Inhibiting the protein-protein interactions between cytokines and their receptors has typically been the domain of therapeutic antibodies. For IL-4 the current clinical standard is the anti-IL-4Rα antibody dupilumab, first approved for treating atopic dermatitis, which inhibits IL-4 activity by binding to the IL-4 membrane receptor (IL-4Rα). The antibody is now approved for treating a wide range of inflammatory conditions including moderate-to-severe asthma, eosinophilic esophagitis, prurigo nodularis, and chronic rhinosinusitis with nasal polyposis.^9–11^ Dupilumab is also currently being clinically investigated for treating metastatic non-small cell lung cancer (NSCLC) in conjugation with PD-L1 after the observation that anti-IL-4 treatment reduced tumor burden in murine models (NCT05013450).^12,13^ Results from a Phase 3 clinical trial assessing dupilumab for Chronic Obstructive Pulmonary Disease (COPD) are expected soon, highlighting yet another indication (Clinical Trial Identifier NCT03930732). In addition to IL-4, blocking cytokine signaling has proven to be a promising therapeutic strategy for multiple conditions, but advances have centered around antibody therapeutics.^14–31^ Monoclonal antibody (mAbs) therapeutics face limitations in administration routes, primarily due to their large size^32^ and hydrophilic nature as glycoproteins.^33^ These characteristics contribute to their gastrointestinal instability and non-specific elimination by the reticuloendothelial system, restricting administration largely to parenteral routes such as predominantly intravenous, growing preference for subcutaneous delivery (∼30%) by strong patient acceptability, while intramuscular and intralesional routes remain rare and indication-specific delivery.^34–41^ These mAbs can pose toxicity issues arising from both target-related effects and immunogenicity (target-independent).^42–45^ The risk of immunogenicity differs among mAbs, being highest in murine and chimeric, lower in humanized, and minimal in fully human antibodies, and can be exacerbated by unnatural linkers or fusion constructs, potentially reducing therapeutic efficacy.^46–51^ Maintaining mAbs therapy presents challenges due to characteristics such as a long half-life, limited tissue residence time, and slower progression to peak concentration.^52–55^ In contrast, small molecules are characterized by low molecular weights, rapid absorption, good bioavailability, lack of immunogenicity, and rapid metabolism that offer advantages such as oral delivery, consistent dosing, and precise control over the timing of therapy.^32,46,52–57^

Recent advances in screening technologies and assays for interrogating protein-protein interactions have renewed interest in directly disrupting cytokine binding with small molecules.^58^ DiCE Molecules and LEO Pharma are pursuing small-molecule or macrocyclic IL-17A inhibitors to rival the anti-IL-17A antibody cosentyx, with both companies in early Phase I clinical trials.^59–61^ Similarly, SPD304, initially an anti-TNFα inhibitor with low potency and poor physiochemical properties, paved the way for the discovery of a benpyrine compound as an anti-TNFα inhibitor.^62–64^ SAR441566 is a TNFα inhibitor that is currently in a Phase 2 Clinical Trial for Rheumatoid Arthritis (NCT06073093). Moreover, a combination of high throughput screening, virtual screening, and biophysical & biochemical assays have been utilized to discover small molecule inhibitors for the soluble cytokines IL-1β, IL-2, IL-33(sST2), IL-36γ, IL-15, and dimeric IL-23.^58,65–70^

We previously reported the discovery of the first small-molecule IL-4 inhibitor which we have named **Nico-52**.^71^ Here, we determined structure-activity relationships around the core scaffold of **Nico-52** to characterize and improve the potency and selectivity of the compound. We evaluated 55 analogs in a HEK-Blue IL-4/IL-13 reporter assay at a single dose of 10 μM. Analogs that inhibited IL-4 signaling >90% were then tested for dose-dependent responses to better characterize their potency, with 2 analogs exhibiting potencies in the nanomolar range. Potent analogs featured structural changes to the p-fluorophenyl substituent whereas the amino nicotinonitrile core and the ortho-hydroquinone appeared to be less amenable to changes. Differential scanning fluorimetry (DSF) screening against related cytokines suggested that **Nico-52** and improved analogs are selective for binding to IL-4 over IL-13, a related protein. We further established that **Nico-52** and improved analogs inhibit both type I and II receptor complex signaling, inhibition of murine IL-4 signaling by **Nico-52** and determined *in vitro* ADME/T properties of our lead analogs. In this study, we determined the salient structural features for small-molecule IL-4 inhibition, identified IL-4 inhibitors with nanomolar potency, characterized their suitability for *in vivo* studies, and performed an *in vivo* assessment of this scaffold in syngeneic murine tumor models (**Figure 1**). We discovered that the ADME/T properties of **Nico-52** and its lead analogs suggest *in vivo* stability and low cytotoxicity, and that the inhibitors suppress B16-F10 (melanoma) and 4T1 (breast) tumor growth *in vivo*. Moreover, it was found that the tumor-associated macrophages (TAMs) from excised B16-F10 tumors were shifted towards the M1 tumor-killing phenotype.

**Figure 1.**
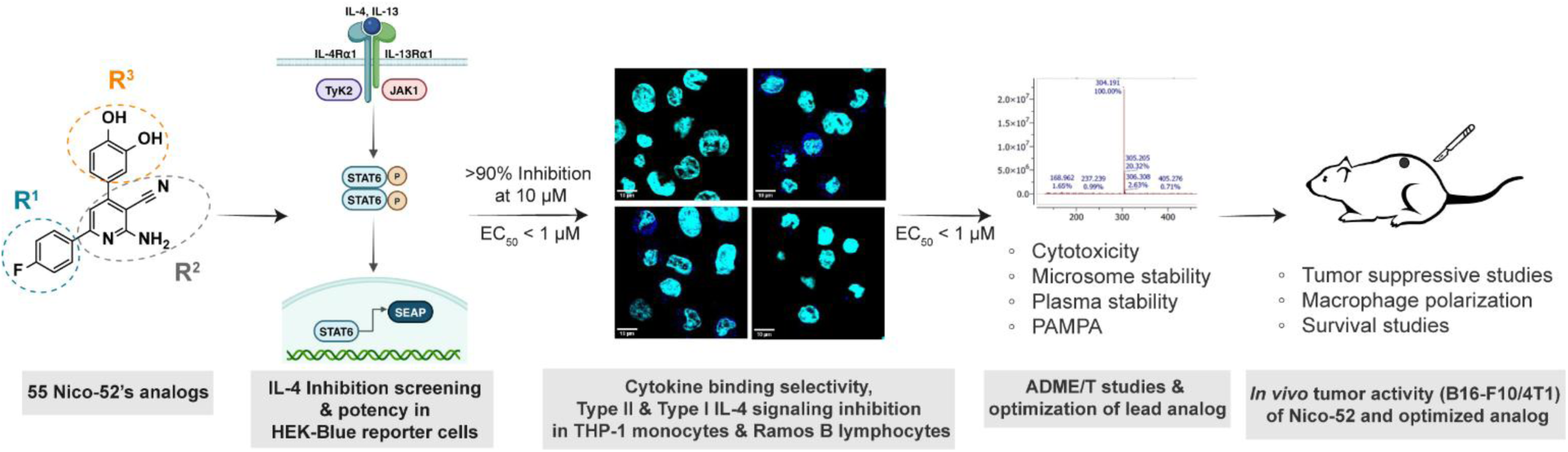
Workflow for identification and functional characterization of Nico-52’s potent analogs. 55 analogs were screened for preliminary inhibition in IL-4/IL-13 HEK blue reporter assay, with analogs that showed >90% inhibition at 10 µM being prioritized for EC_50_ determination by dose-response analysis. Analogs with EC_50_’s less than 1 µM were further characterized for their cytokine binding selectivity and inhibition of Type II and Type I IL-4 signaling in non-engineered THP-1 monocytes and Ramos B lymphocytes, respectively. The *in vitro* ADME/T properties of **Nico-52** and lead analogs were also determined and lead to the development of a pharmacokinetically stable and optimized lead analog **53**. The anti-tumor activity of both **Nico-52** and lead analog **53** were evaluated *in vivo* in B16-F10 and 4T1 murine tumor models.

## RESULTS

### Nico-52 Analog Design and Synthesis

A total of 55 analogs were tested for preliminary inhibition, of which 44 analogs were synthesized and 11 analogs were commercially sourced. We pursued a three-part model for structural elaboration to determine structure-activity relationships (SAR) **(Figure 2)** based on the 3-ring system that comprises the parent compound: the p-fluorophenyl (R^1^), the ortho-hydroquinone (R^2^), and the core amino nicotinonitrile (R^3^). The synthesis of Nico-52 and its analogs was adapted from Serry et al^72^ which used a one-pot, three-component reaction to combine an acetophenone, an aromatic aldehyde, and malononitrile using ammonium acetate and refluxed in ethanol for 10-14 hours **(Figure S1A)**. We were able to access the parent compound Nico-52 and apply this method widely to synthesize the analogs described here with yields varying based on the components involved. Hydroxy nicotinonitrile R^3^ analogs were synthesized by replacing malononitrile with ethyl cyanoacetate **(Figure S1B)**.

**Figure 2.**
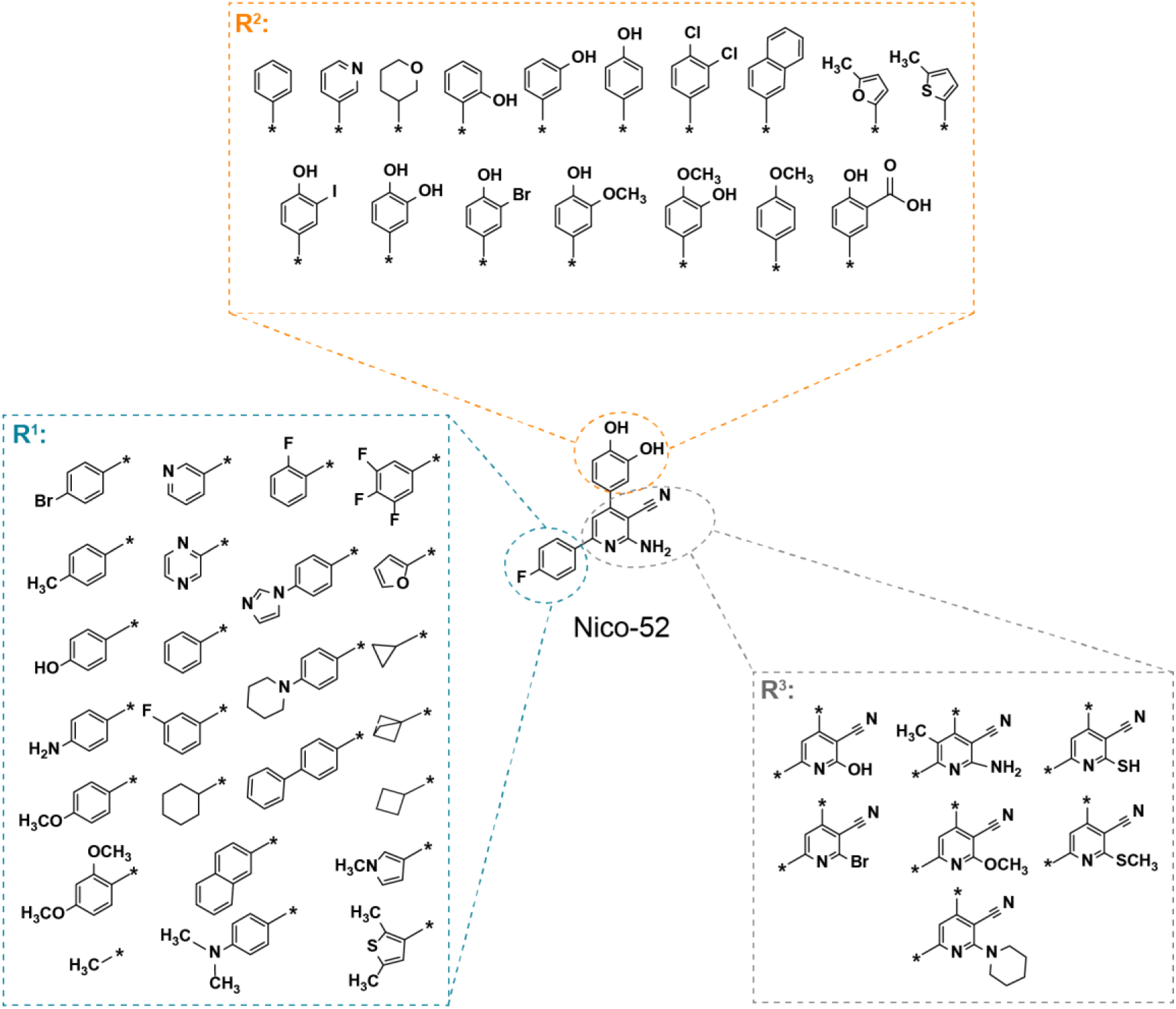
Nico-52 Analogs and Synthesis Scheme for Investigating Structure-Activity Relationships. Structural changes at the three rings comprising **Nico-52** to establish which features contribute to **Nico-52’s** binding to IL-4.

### Cellular IL-4 Inhibition of Nico-52 Analogs

All **Nico-52** analogs were initially evaluated for their IL-4 inhibition using a well-established HEK Blue IL-4/IL-13 reporter assay at a 10 μM concentration **(Table S1,2,3)**.^71,73,74^ Since **Nico-52** shows near complete inhibition (>97%) at this concentration, analogs could be efficiently grouped as either being at least as potent or less potent than the parent compound. Analogs that showed similar inhibition at 10 μM (>90% inhibition) were then prioritized for a complete dose-response EC_50_ determination in this reporter system to further determine which analogs possess increased potency over **Nico-52**. Analogs were then prioritized based on their potency for further testing in non-engineered cell lines for IL-4 inhibition of the type II IL-4Rα/IL-13Rα1 receptor complex (THP-1 cells)^71,75,76^ and type I IL-4R/γc receptor complex (Ramos cells).^77,78^ The binding selectivity of our top analogs was also evaluated against other related cytokines by differential scanning fluorimetry (DSF, thermal shift) as well as functional inhibition of IL-13, the closest functional homolog to IL-4 which shares the type II receptor. We then determined the *in vivo* potential of **Nico-52** and our lead analogs through *in vitro* ADME/T studies. The parent compound and our best analog were then evaluated in anti-cancer studies using syngeneic murine tumor models, where anti-tumor activity can be measured against a tumor microenvironment with an immunocompetent background.

#### R^1^ analogs

We selected the R^1^ position for systematic substitution, focusing on the para-fluoro group and the adjacent aromatic core, to determine how positional changes of the fluoro, different para substituents (halogen, hydroxy, methoxy, methyl, aniline), heteroaromatic replacements, ring extensions, and loss of aromaticity affect IL-4 inhibitory activity and to probe the roles of hydrogen bonding, steric bulk, and pKa/protonation on activity.

Altering the F-group showed that fluoro positioning critically impacted activity: moving the fluoro to the ortho position in analog **1** abolished activity, placing it at the meta position in analog **2** reduced inhibition to 40% **(Table S1)**, whereas para substitutions such as **3** (p-bromine), **4** (p-hydroxy), **5** (p-methyl), and **14** (aniline) exhibited >80% IL-4 inhibition. A p-methoxy group in analog **6** gave 81.7% inhibition and was tested specifically to distinguish the hydrogen-bonding effects of the phenoxy group in analog **4**. Notably, p-aniline in analog **14** and the unsubstituted phenyl in analog **15** produced near-complete inhibition similar to Nico-52 (94.5% and 95.5%, respectively).

Altering the aromatic core further revealed that a pyridinyl ring in analog **7** retained potency (94.3%), whereas a furanyl ring in analog **8** abolished activity and a pyrazine ring in analog **9** lowered inhibition to 64.2%. Extension of the aromatic system with naphthyl or biphenyl motifs in analogs **10** and **11** was poorly tolerated. Removal of aromaticity with a cyclohexane derivative in analog **12** reduced inhibition to 49.4%, while replacing the phenyl ring with a methyl group in analog **13** retained substantial potency (70.2%). Following the comparable activity of aniline **14**, we evaluated commercially sourced analogs **16**, **17**, and **18** to probe pKa effects on the aniline. These either abolished activity **(16)**, or substantially reduced it **(17)**, or retained most activity **(18)**, suggesting that steric effects at the aniline nitrogen are more important than modulation of its protonation state and that aromatic substitution is not strictly required for activity, as evidenced by near-total inhibition from the unsubstituted phenyl analog **15**.

In summary, the R^1^ position appears tolerant to different substitutions, as compound activity was comparable with halogen, amino, methoxy, and methyl substitutions. Given these determinations, we prioritized **3**, **7**, **14**, and **15** for dose-response evaluations.

#### R^2^ analogs

We selected the R^2^ position for systematic modification to assess the role of the ortho-hydroquinone in IL-4 binding and to determine how variations in hydroxyl positioning, replacement with methoxy groups, or heteroaromatic substitutions influence activity **(Table S2)**. Removal of the diol in analog **19** nearly abolished inhibition, showing only 6.13% at 10 μM, and similar reductions were observed with heterocycle replacements in analogs **20** (33.9%) and **21** (37.5%), highlighting the critical contribution of the hydroxyl groups to polar and hydrogen-bonding interactions with the protein. Altering the hydroxyl pattern or substituting with methoxy groups revealed that activity depends strongly on hydroxyl positioning: a single hydroxyl at the 4-position in analog **22** decreased inhibition to 45.2%, whereas the ortho-hydroxy analog **23** retained near-complete activity at 98.8%, and the meta-methoxy, para-hydroxy analog **24** preserved >99% inhibition, comparable to the parent compound. Bulky or poorly positioned modifications, including the pyridinyl replacement in analog **26** (0% inhibition) or halogen/carboxyl substitutions at the meta position in analogs **27** (12.4%), 28 (15.7%), and 29 (9.8%), were not tolerated, indicating that although hydrogen bonding at the ortho and para positions is essential, the binding site cannot accommodate larger H-bonding groups or substitutions at the meta position.

Based on the IL-4 inhibition profile at 10 μM, it is clear that the ortho and para positions of the diol are critical for maintaining activity, whereas modifications at the meta position or introduction of sterically demanding groups substantially reduce inhibition. Analogs **23** and **24** maintained near-complete inhibition, demonstrating that carefully positioned hydroxyl or methoxy substitutions can preserve binding interactions, and these analogs were therefore selected for subsequent dose-response studies to further quantify their potency and validate their structure–activity relationships.

#### R^3^ and commercial analogs

We selected the amino group and para substituents of the nicotinonitrile core for systematic modification to assess how these changes influence IL-4 inhibition **(Table S3)**. Substitution of the amino group in the parent compound with hydroxy nicotinonitrile **(30)** resulted in considerable but reduced inhibition at 10 μM (71.2%), whereas 5-methyl amino nicotinonitrile **(31)** retained near-complete inhibition (97.7%), suggesting that the core structure can tolerate slight modifications at the 2 and 5 positions **(Table S3)**. Commercially sourced analogs **(32–41)** provided additional insights, even though some contain changes at multiple positions and cannot be directly compared to the parent compound. Most of these derivatives showed reduced IL-4 inhibition, with the exception of analog **35**, which inhibited IL-4 by 96.8% and features both a para substitution at R^1^ and a methyl at the R^3^ 5-position, suggesting that the R^3^ methyl can offset the inhibitory impact of the R^1^ methoxy substitution, as reflected by comparison with analog **6** (81.7%). Analogs **36** and **37**, containing p-methoxy at R^1^ and a phenyl group at R^2^, displayed inhibition comparable to analog **6**, consistent with the near-zero activity of analog **19** (6.13%) and highlighting the contribution of thiol/sulfide additions to the nicotinonitrile core. To further probe the role of the amino group, hydroxy nicotinonitrile analogs of compound **30** were prepared, which also allowed access to several R^1^ and R^2^ analogs that were difficult to isolate in the amino series. Among all R^3^ hydroxy nicotinonitrile analogs, only the ortho-phenoxy derivative **49** achieved inhibition greater than **31** and comparable to the parent compound (>99%), emphasizing the importance of ortho-hydroxyl substitution at R^1^ for hydrogen bonding, similar to observations for analog **23**. In contrast, analog **43**, which combines an aniline at R^1^ with a carboxyl at R^2^ (similar to analogs **14** and **27**), exhibited poor inhibition (6.63%), indicating that this combination of modifications is unfavorable for binding. Based on these trends and the IL-4 inhibition profiles at 10 μM, analogs **31**, **35**, and **49** were prioritized for subsequent dose-response evaluation. Together, these findings indicate that modest substitutions at the amino group or R3 can be tolerated, whereas bulky or mismatched groups at R1 or R2, and particularly carboxylates, disrupt hydrogen bonding and sharply reduce IL-4 inhibition.

Taken together, the trends across the three-substitution series reveal clear structure–activity relationships. For R^1^, IL4 inhibition was largely maintained with halogen, hydroxy, methoxy, methyl, or aniline groups at the para position, whereas moving the fluoro to the ortho position eliminated activity and placing it at the meta position weakened it. Replacing the aromatic ring with larger or less electron-rich systems also caused a marked loss of potency. For R^2^, the data highlighted the critical role of the ortho and para hydroxyl groups of the hydroquinone, which supported hydrogen bonding and polarity required for binding, while modifications at the meta position or introduction of bulky or charged groups sharply reduced activity. In R^3^ and the commercial analogs, minor changes at the amino group or at the 5 position of the core were tolerated, but extensive modification at R^1^ or R^2^ within this set, especially the addition of carboxyl or large groups, sharply reduced inhibition. Overall, the optimal profile for IL4 inhibition emerged as a nicotinonitrile scaffold with an unsubstituted or simply substituted aromatic ring at R^1^, an intact ortho hydroquinone at R^2^, and modestly tolerated substitutions at R^3^, with bulky or mismatched groups being detrimental.

#### Summary of Optimal Structural Features

Across all series, we prioritized nine analogs that showed similar IL-4 inhibition in our HEK-Blue IL-4/IL-13 reporter assay to the parent compound at 10 μM (> 90%) and determined their EC_50_ of inhibition **(Table 1)** to further define their efficacy in the reporter cell line. Analogs **15** and **14** with modified R^1^ positions demonstrated nanomolar potency of 551.6 nM and 627.2 nM, respectively **(Table 1**, **Figure 3A)**. These results supported the observation that the R^1^ position was tolerant to derivatization and changes can improve potency of the parent compound. Furthermore, that unsubstituted phenyl moiety of **15** was the most potent analog suggesting that while substitution of the R^1^ aromatic ring can be tolerated, optimal inhibition is achieved without it. For R^2^ analogs the 3,4-diol of the ortho-hydroquinone appears to play a critical role in maintaining potency, likely through hydrogen bonding, with none of the other prioritized R^2^ analogs able to produce lower EC_50_’s than the parent compound. Modification of the hydroxyl groups with methoxy groups did not improve potency as seen in analog **24** (EC_50_ = 6.4 μM). Interestingly, a single hydroxyl could retain considerable potency in the R^2^ position as the phenol in analog **23** demonstrated a potency of 3.4 μM, suggesting that conformational restriction of the axial biphenyl bond may also be important along with hydrogen bonding at this position. For R^3^ and the commercial analogs, none of the changes produced compounds with improved potency. Analog **31** inhibited IL-4 in the cell reporter assay with less potency than **Nico-52**. Analog **35**, with a substituted methoxy group at the R^1^ para position and methyl at the R^3^ 5-position, had a potency of 4.22 μM. Analogs **31** and **35** showcased that a methyl group at the R^3^ 5-position does not improve IL-4 inhibition. Hydroxy nicotinonitrile analog **49** with an R^2^ ortho alcohol had a potency of 1.75 μM similar to the parent compound. Taken together, the optimal structural features we identified for IL-4 inhibition is a nicotinamide scaffold with an unsubstituted aromatic ring at R^1^ and an ortho-hydroquinone at R^3^. Our most potent derivatives (EC_50_ < 1 μM) **14** and **15** were advanced for further testing as lead analogs.

**Table 1.**
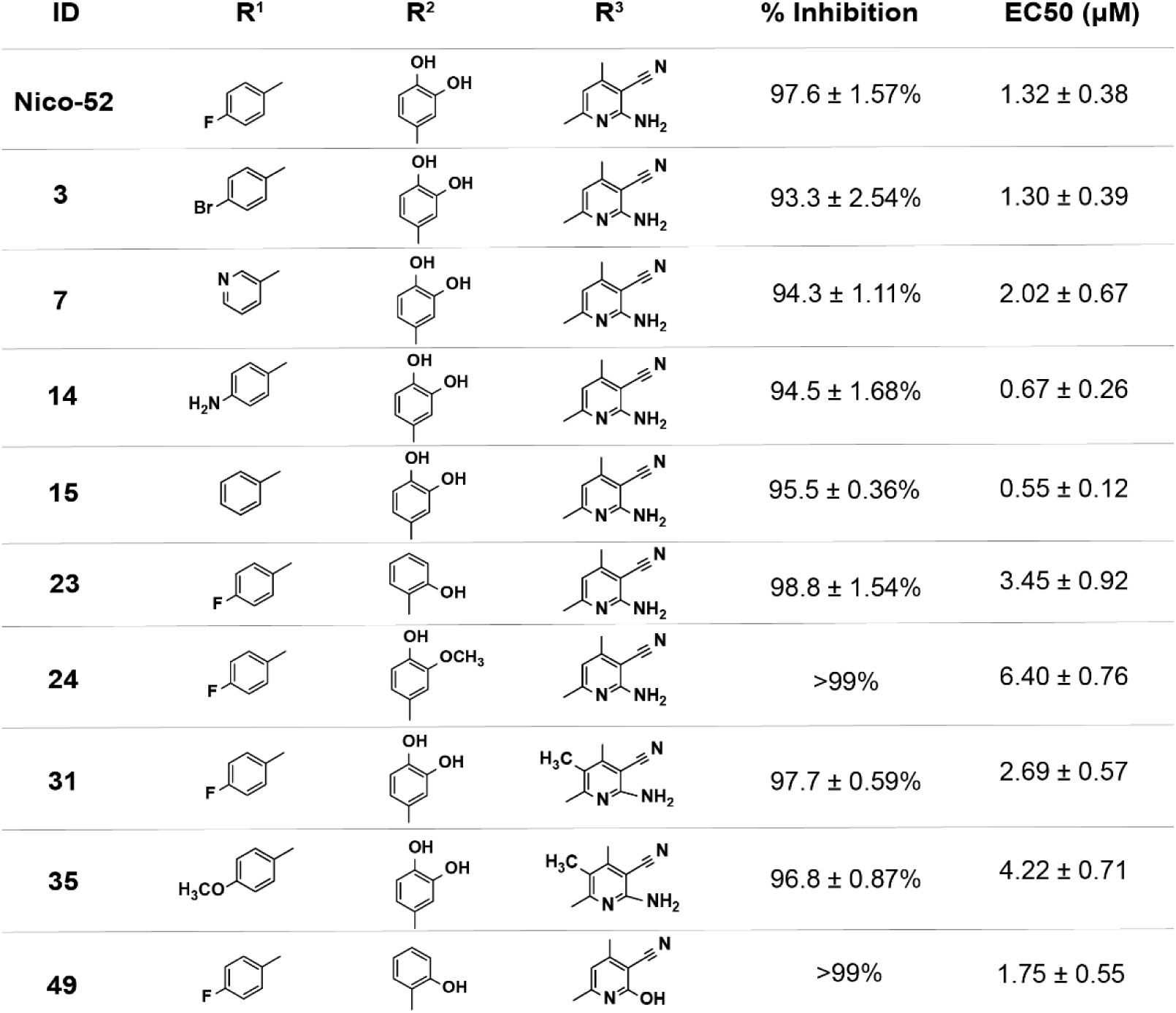
Prioritized analogs for dose-response evaluation in HEK Blue IL-4/IL-13 reporter assay. For each analog, the structural change made at each position (R^1^, R^2^, or R^3^) is indicated. Blank spaces indicate that the structural position matches that of the parent compound Nico-52. For analogs that were comparably potent to Nico-52 at 10 μM, the EC_50_ was determined, and the coefficient of determination (r2) is reported in **Table S4**.

| ID | R<sup>1</sup> | R<sup>2</sup> | R<sup>3</sup> | % Inhibition | EC<sub>50</sub> (μM) |
| --- | --- | --- | --- | --- | --- |
| **Nico-52** |  |  |  | 97.6 ± 1.57% | 1.32 ± 0.38 |
| **3** |  |  |  | 93.3 ± 2.54% | 1.30 ± 0.39 |
| **7** |  |  |  | 94.3 ± 1.11% | 2.02 ± 0.67 |
| **14** |  |  |  | 94.5 ± 1.68% | 0.67 ± 0.26 |
| **15** |  |  |  | 95.5 ± 0.36% | 0.55 ± 0.12 |
| **23** |  |  |  | 98.8 ± 1.54% | 3.45 ± 0.92 |
| **24** |  |  |  | >99% | 6.40 ± 0.76 |
| **31** |  |  |  | 97.7 ± 0.59% | 2.69 ± 0.57 |
| **35** |  |  |  | 96.8 ± 0.87% | 4.22 ± 0.71 |
| **49** |  |  |  | >99% | 1.75 ± 0.55 |

**Figure 3.**
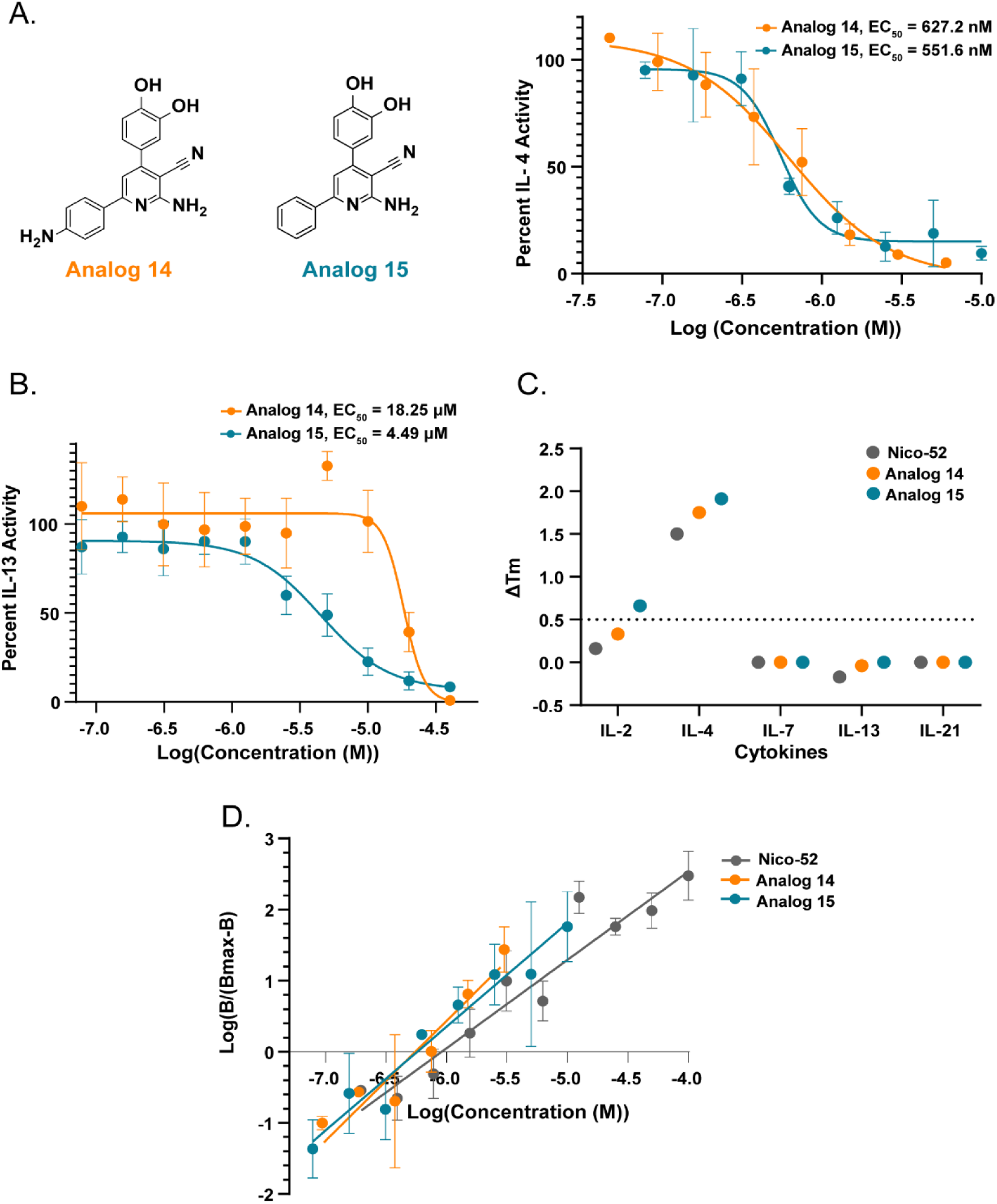
Nico-52 potent analogs dose response in HEK Blue IL-4/IL-13 reporter assay and selectivity across structurally and functionally related cytokines. **(A)** Dose dependence and EC_50_ of **14** and **15** in the HEK Blue IL-4/IL-13 reporter assay. **(B)** IL-13 inhibition by analogs **14** and **15** in the HEK Blue IL-4/IL-13 reporter assay. **(C)** Thermal shift analysis of **Nico-52**, **14,** and **15** against IL-2, IL-4, IL-7, IL-13, and IL-21 at 2 μM (0.1% DMSO) final concentration, a cut-off = 0.5℃ was used to determine the threshold for the determination of binding events. **(D)** Hill plots of **Nico-52**, **14,** and **15** dose responses in the HEK Blue IL-4/IL-13 reporter assay. The Hill coefficients were determined for **Nico-52** = 1.24 ± 0.08, **14** = 1.69 ± 0.25, and for analog **15** = 1.49 ± 0.15. *Error bars represent mean ± SD*

### Selectivity of Lead Analogs

IL-4 and IL-13 share the same type II receptor for their proinflammatory signaling.^77^ This receptor consists of the heterodimerization of IL-4Rα and the IL-13Rα1 subunit and can also be activated by IL-13 binding.^78^ Previously, we reported that **Nico-52** had 10-fold higher inhibition for IL-4 activity than IL-13 activity.^71^ To determine if our more potent analogs **14** and **15** maintained this level of selectivity, they were tested for dose-dependent inhibition of IL-13 activity in the HEK Blue IL-4/IL-13 reporter assay. Analogs **14** and **15** inhibited soluble IL-13 mediated signaling with significantly less potency, with an EC_50_ = 18.25 µM (29-fold less than for IL-4) and an EC_50_ = 4.49 µM (8-fold less than for IL-4), respectively (**Figure 3B)**.

IL-4 also mediates signal transduction via a type I receptor complex, which contains IL-4Rα and a common cytokine receptor γ_c_ chain shared by IL-2, IL-7, and IL-21 cytokines.^77^ We therefore tested **Nico-52** and analogs **14** and **15** for target specificity against the soluble cytokines IL-2, IL-7, IL-21, and IL-13 at a 2 µM final concentration in a thermal shift assay using DSF. Analog **15** specifically binds to soluble IL-4 with a T_m_ shift of 1.75 ℃ and analog **14** binds to soluble IL-4 with a T_m_ shift of 1.66 ℃ **(Figure 3C)**. While analog **15** also bound weakly to soluble IL-2 with a T_m_ shift of 0.66 ℃, this just barely crossed our 0.5 ℃ shift threshold to qualify as a binder and the thermal shift for IL-4 was much stronger than IL-2 by over 1℃. To further assess selectivity, we evaluated analog **15** in the HEK-Blue IL-2 reporter assay at 1 and 10 µM, where it did not alter IL-2–induced SEAP production, confirming its selectivity for IL-4 **(Figure S2)**. Taken together, these results further supported that **Nico-52** and analogs **15** and **14** are indeed strongly selective towards soluble IL-4.

For all three compounds, **Nico-52**, **14**, and **15** dose-response data was fitted using simple linear regression where each data point was normalized based on the maximal response to determine the Hill coefficient, providing insights into the binding dynamics of these compounds. All three compounds showed positive cooperativity with a Hill coefficient >1, suggesting that there may be multiple binding sites on the IL-4 surface. The Hill coefficient for **Nico-52** was 1.24 ± 0.08, for analog **14**: 1.69 ± 0.25, for analog **15**: 1.49 ± 0.15 **(Figure 3D)**.

### Inhibition in Cells that Natively Express the IL-4 Receptor

While the HEK Blue IL-4/IL13 reporter assay is a useful tool to identify compounds with inhibitory activity, we sought to further characterize the inhibitory activity and potency of the analogs against cell lines that are not engineered and natively express the IL-4 complexes. IL-4 binding its receptor in both type I and type II signaling leads to JAK1-mediated STAT-6 phosphorylation.^73^ THP-1 monocytes express exclusively the type II IL-4 complex ^74^ **(Figure 4A)**. To confirm the inhibitory effects of analogs **14** and **15** in this cell line and compare them to our previously reported results for **Nico-52**^71^ (EC_50_ = 3.1 µM), THP-1 monocytes were treated with either soluble IL-4 or IL-4 pre-incubated with these potent analogs in a dose-dependent manner. Levels of STAT-6 phosphorylation were then quantified by western blot to measure pathway activation. Analog **14** inhibited STAT-6 phosphorylation with an EC_50_ = 686.5 nM which was consistent with our HEK Blue IL-4/IL-13 reporter assay data **(Figure 4B&C S3, and S4)**. STAT-6 levels were not impacted by analog **14**. Surprisingly, analog **15** inhibited IL-4 induced STAT-6 phosphorylation with an EC_50_ = 49.6 nM which was ∼9 fold more active than what was observed in the HEK Blue IL-4/IL-13 reporter assay **(Figure 4B&D, S5, and S6)**. STAT-6 levels were not impacted by analog **15**. The difference in potency between the engineered reporter system and the THP-1 cells may be attributed to a difference in expression levels of the type II receptor complex for the two cell lines. To further corroborate our results, we examined analogs **14** and **15** in a dose-dependent inhibition of IL-4 induced pSTAT-6 levels using immunofluorescence. Treatment of THP-1 cells with IL-4 and analog **14** again demonstrated a significant reduction in STAT-6 phosphorylation at 500 nM and almost no visible STAT-6 phosphorylation at a 5 µM concentration in comparison to IL-4 treated with vehicle (RPMI-1640, 2% DMSO) **(Figure 4E-I)** Additionally, THP-1 cells treated with analog **15** were analyzed using immunofluorescence. Treatment of THP-1 cells with analog **15** exhibited reduced STAT-6 phosphorylation at 50 nM and complete inhibition of phosphorylation at 500 nM **(Figure 4J-O)**.

**Figure 4.**
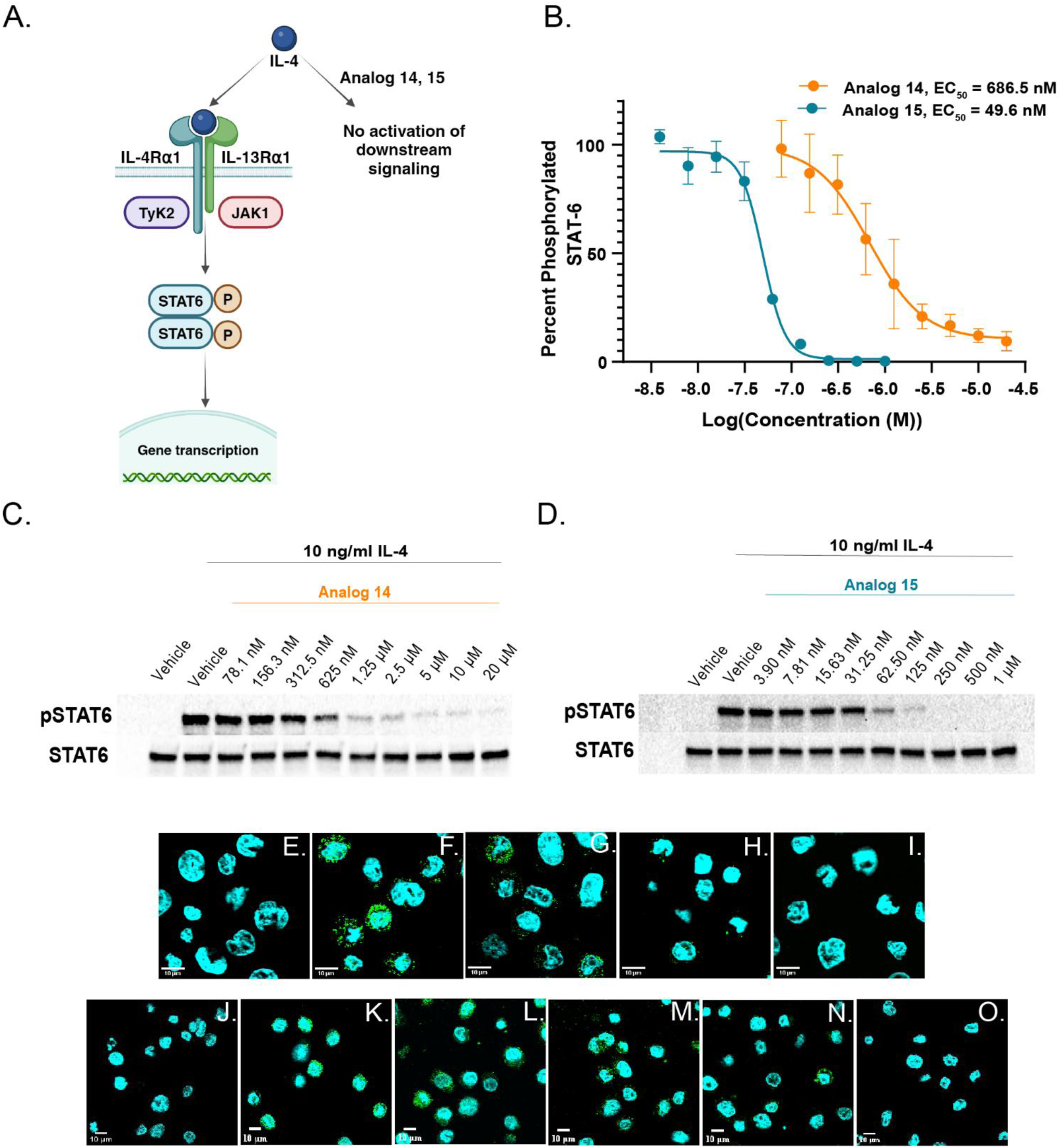
Dose-dependent inhibition of IL-4 induced STAT-6 phosphorylation in THP-1 monocytes with analogs 14 and 15 (type II IL-4 receptor). **(A)** IL-4 type II signaling schematic. **(B)** Dose-dependent IL-4 inhibition with **14** and **15** in THP-1 monocytes, normalized to STAT-6 levels. **(C)** Western blots of pSTAT-6 to STAT-6 upon treatment with IL-4 and vehicle or analog **14**. *Error bars represent mean ± SD.* (D) Western blots of pSTAT-6 to STAT-6 upon treatment with IL-4 and vehicle or analog **15**. Immunofluorescence of THP-1 monocytes upon treatment with **(E)** vehicle alone, **(F)** IL-4 + vehicle, **(G)** IL-4 + 50 nM **14**, **(H)** IL-4 +500 nM **14**, **(I)** IL-4 + 5 μM **14**. Immunofluorescence of THP-1 monocytes upon treatment with **(J)** vehicle alone, **(K)** IL-4 + vehicle **(L)** IL-4 + 5 nM **15**, **(M)** IL-4 + 50 nM **15**, **(N)** IL-4 + 500 nM **15**, **(O)** IL-4 + 5 μM **15** (Green: pSTAT-6, Cyan: nucleus) *(Vehicle: RPMI-1640, 2% DMSO)*

Inhibitory effects of the parent compound (**Nico-52**) and our lead analogs **14** and **15** were also evaluated in Ramos B lymphocytes which exclusively express the IL-4 type I receptor complex^75,76^ **(Figure 5A)**. Ramos cells were exposed to IL-4 with vehicle (RPMI-1640, 2% DMSO) or **Nico-52** in a dose-dependent manner, and again pSTAT-6 relative to STAT-6 levels were quantified by western blot analysis. **Nico-52** inhibited IL-4 induced STAT-6 phosphorylation with an EC_50_ = 1.03 µM (**Figure 5B, S7, S8)**. A comparative evaluation of potency between **Nico-52**, **14**, and **15** was conducted in Ramos B lymphocytes employing a similar method of quantification of pSTAT-6 in relation to STAT-6 as discussed above. Expectedly at 50 nM, minimum inhibition of phosphorylation of STAT-6 by **Nico-52**, **14,** and **15** was observed. At 500 nM, analog **15** demonstrated a percent inhibition of 61.9%, and analog **14** and **Nico-52** exhibited a percent inhibition of 31.4% and 14.6% respectively **(Figure 5B, S9, S10, S11, and S12)**.

**Figure 5.**
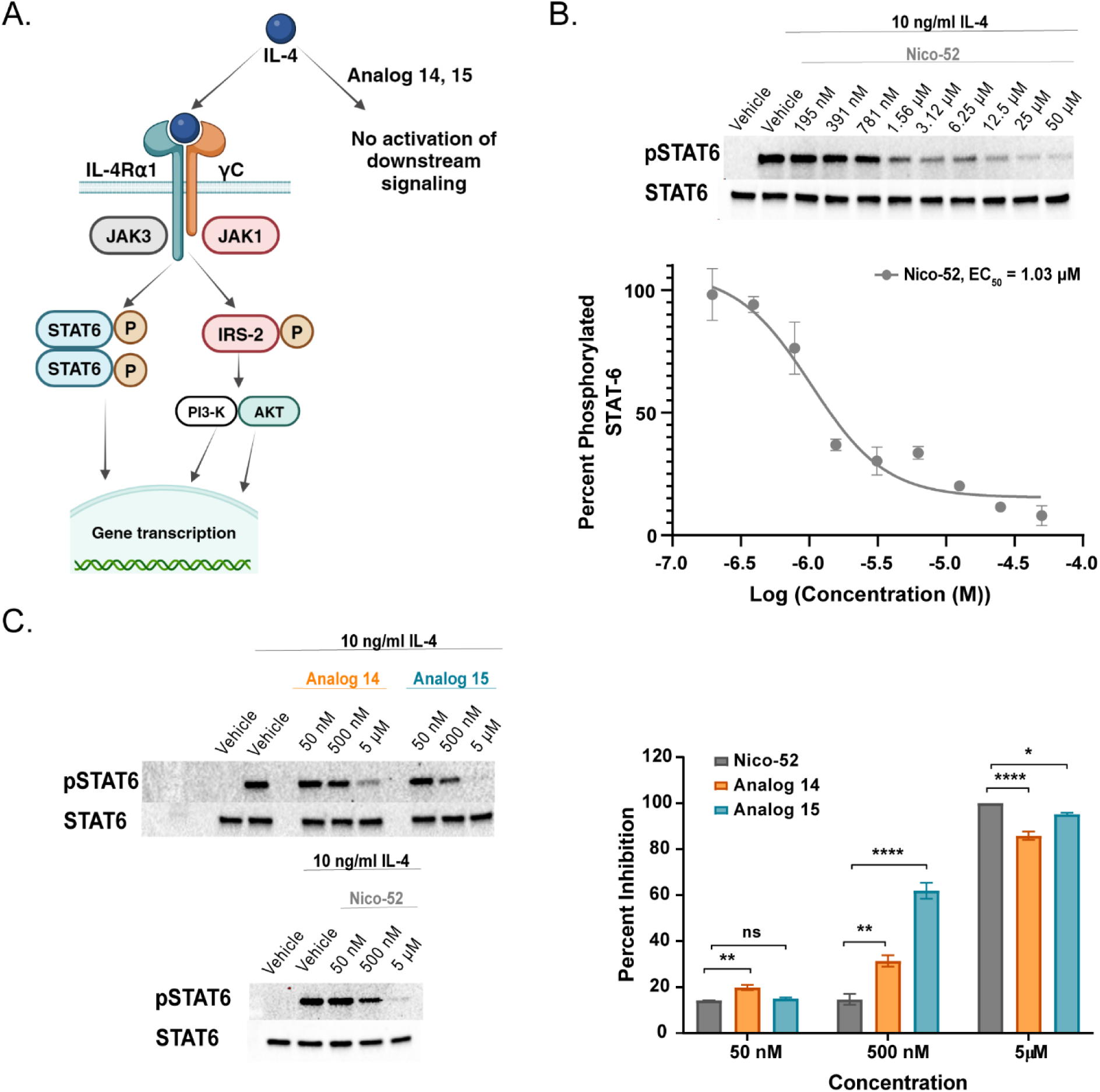
Nico-52 and lead analogs dose-dependent inhibition of IL-4 induced STAT6 phosphorylation in Ramos B lymphocytes (type I IL-4 receptor). **(A)** IL-4 type I signaling schematic. **(B)** Western blots of pSTAT-6 and STAT-6 upon treatment with IL-4 and vehicle/**Nico-52**. **Nico-52** quantified dose-dependent IL-4 inhibition in Ramos B lymphocytes normalized to STAT-6. **(C)** Western blots of pSTAT-6 to STAT-6 upon treatment with IL-4 with vehicle**/Nico-52**/**14**/**15**. Quantified **Nico-52**, **14** and **15** percent IL-4 inhibition at 50 nM, 500 nM, and 5 μM, normalized to STAT-6. Significance determined by Dunnett’s multiple comparison test * = p < 0.05, ** = p < 0.01, **** = p < 0.0001, ns = not significant. *Error bars represent mean ± SD*

At a 5 µM concentration, near complete inhibition was observed for all three compounds. Percent Inhibition at 500 nM was more informative as it revealed that analog **15** demonstrated significantly higher levels of inhibition at 500 nM followed by analog **14** and then Nico-52. This result supported that analogs **14** and **15** also have higher potency in Ramos B lymphocytes for type I signaling as well in comparison to **Nico-52**.

Finally, to evaluate if the **Nico-52,** analog **14** and **15** are also suitable for the inhibition of murine IL-4 signaling and animal studies in mice, RAW 264.7 macrophages were incubated with murine IL-4 and **Nico-52/**Analog **14/**Analog **15** or vehicle (DMEM, 2% DMSO). Dose-dependent inhibition of STAT-6 phosphorylation was measured and quantified via western blotting **(S13-S18)**. **Nico-52** and Analog **15** exhibit a dose-dependent reduction in pSTAT-6 levels with an EC_50_ of 1.28 µM and ∼21 nM respectively whereas Analog **14** showed no inhibition of IL-4 induced STAT-6 phosphorylation. STAT-6 levels were not affected by **Nico-52/**Analog **14/**Analog **15**. Crystal structure analysis of IL-4 bound to its receptor ⍺ chain (IL-4R⍺) in an intermediate complex revealed three discrete interaction clusters (I, II, and III) which are formed by distinct polar and charge-complementary interactions at the binding interface^79^. Prior structural studies have implicated Cluster II residues as a critical interaction surface for receptor engagement, while Clusters I and III provide additional stabilizing contacts.^80,81^ To evaluate small-molecule interactions in this context, we applied Boltz-2^82^ using the IL-4 FASTA sequence with **Nico-52** and Lead Analog **53**. The predictions localized both Nico-52 **(Figure 6A, blue)** and Lead Analog 53 **(Figure 6B, magenta)** to residues within Cluster II, consistent with its role as a central binding hotspot. Although Clusters I (yellow) and III (cyan) lie in close spatial proximity, no direct ligand contacts were observed. Boltz-2 assigned high confidence scores (∼0.96) for Cluster II binding in both cases.

**Figure 6.**
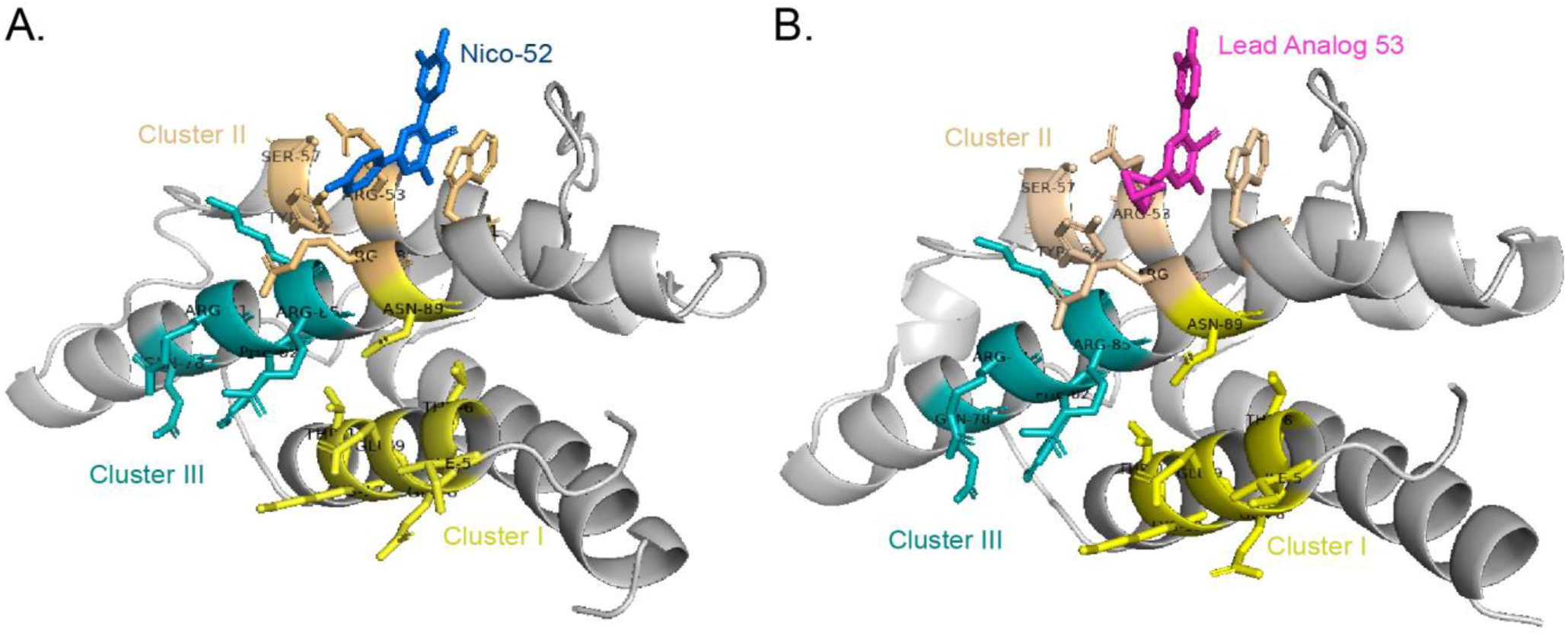
Predicted binding of small molecules to IL-4 using Boltz-2. (A) Nico-52. (blue) predicted to bind Cluster II residues of IL-4. **(B)** Lead Analog **53** (magenta) similarly predicted to localize within Cluster II.

### *In vitro* ADME/T of Lead Analogs

**Nico-52**, analogs **14** and **15** were evaluated for cytotoxicity in B16-F10 cells at 25 µM, revealing no toxicity after a 24-hour incubation period. Metabolic stability assessment utilizing mouse hepatic microsomes indicated that after one hour **Nico-52** and **14** demonstrated retention of 86.5% and 75.4% of their initial concentrations, respectively. In contrast, analog **15** exhibited rapid degradation with only 30.3% remaining after the same duration. All three compounds displayed stability in both mouse and human plasma. After three hours, 97% of **Nico-52** was retained in mouse plasma and > 99% in human plasma. Analog **14** displayed > 99% retention in both mouse and human plasma. Similarly, analog **15** also exhibited significant retention with 94.3% in mouse plasma and 97.4% in human plasma. PAMPA assays unveiled distinct permeability coefficients comparable to other membrane-permeable, orally bioavailable compounds, further highlighting the potential of this scaffold **(Table 2)**.

**Table 2.** *In vitro* ADME/T of Nico-52 and lead analogs. **Nico-52, 14,** and **15** cytotoxicity evaluation at 25 µM in B16-F10 cells for 24 hours. All three compounds were also tested for mouse hepatic microsome stability, mouse & human plasma stability, and PAMPA permeability.

| ADME/T | Nico-52 | Analog 14 | Analog 15 |
| --- | --- | --- | --- |
| Cytotoxicity in B16-F10 cells (24 hours) | n.s. < 25 μM | n.s. < 25 μM | n.s. < 25 μM |
| Mouse hepatic microsome stability<br>(% remaining after 1 hour) | 86.5 % | 75.4 % | 30.3 % |
| Mouse/Human plasma stability<br>(% remaining after 3 hours) | 97 % / >99 % | >99 % / >99 % | 94.3 % / 97.4 % |
| PAMPA Permeability<br>($\times 10^{-6}$ cm/s) | 16.4 | 6.2 | 12.7 |

Since **15** was also our most potent analog, we sought to mitigate its metabolic liability. A closer examination of the LC/MS profile from the microsomal stability assay indeed supported oxidation (hydroxylation) of the analog (data not shown). Only the R^1^ position varied between **15** and the parent compound, so we hypothesized that implementing a bioisostere for the phenyl group at this position could improve metabolic stability. The phenyl ring at R^1^ was substituted with three bioisosteres: bicyclopentanyl (**53**), cyclopropyl (**54**), and cyclobutyl (**55**) groups. We then first verified if these new analogs retained some of the potency of **15.** Analog **53** with the bicyclopentane moiety exhibited inhibition of 86.1% at 10 µM in the HEK Blue IL-4/IL-13 reporter assay **(Table 3)**, while the remaining two analogs lost considerable potency. Additionally, analog **53** exhibits an EC_50_ of 690 nM in a dose-dependent evaluation in HEK Blue IL-4/IL-13 reporter assay **(Figure S19)**. Satisfyingly, **53** showed improved metabolic stability compared to **15** with 68.3% remaining after an hour upon treatment with mouse hepatic microsomes **(Table 3)**. Given these results, we then determined the *in vivo* anti-cancer potential of **Nico-52** and optimized analog **53.**

**Table 3.**
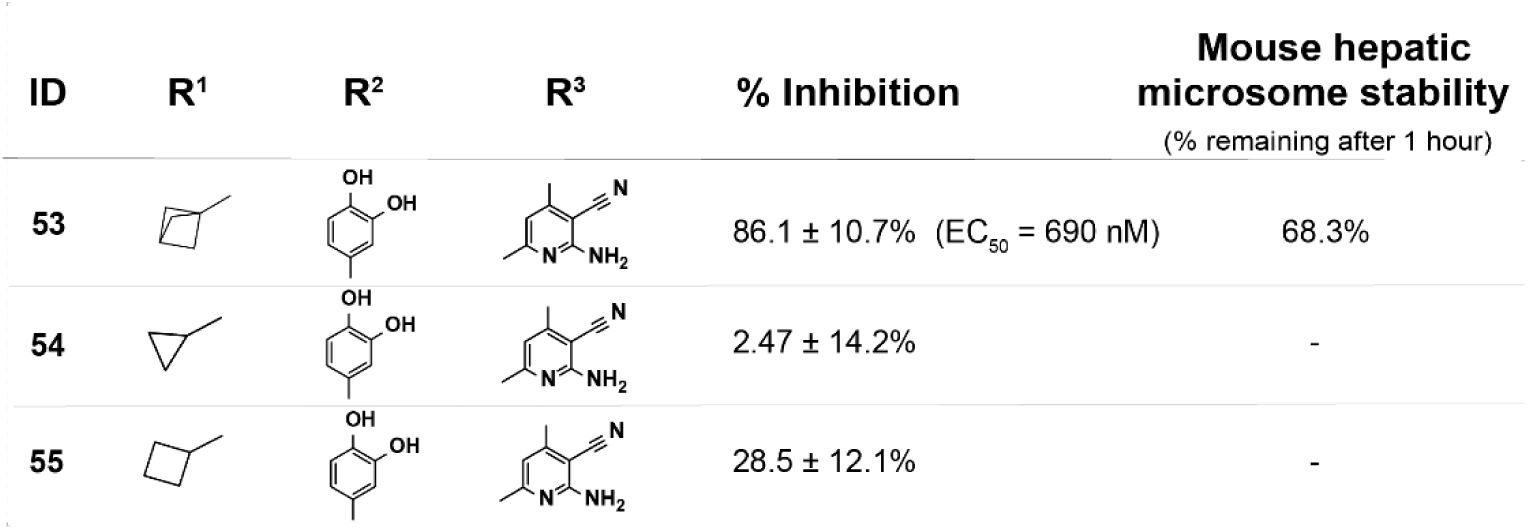
Preliminary optimization of analog 15 for metabolic stability. Analogs of **15** with phenyl group bioisosteres at the R^1^ position were first evaluated for IL-4 inhibition in the HEK Blue IL-4/IL-13 reporter assay at 10 µM. Having retained activity, analog **53** was further evaluated for metabolic stability in the mouse hepatic microsomal stability assay, where 68.3% of it remained after an hour treatment with mouse hepatic microsomes.

| ID | R<sup>1</sup> | R<sup>2</sup> | R<sup>3</sup> | % Inhibition | Mouse hepatic microsome stability<br>(% remaining after 1 hour) |
| --- | --- | --- | --- | --- | --- |
| 53 | | | | 86.1 ± 10.7% (EC<sub>50</sub> = 690 nM) | 68.3% |
| 54 | | | | 2.47 ± 14.2% | - |
| 55 | | | | 28.5 ± 12.1% | - |

### Nico-52 suppresses solid tumor growth in the *in vivo* B16-F10 model

The anti-tumor effects of **Nico-52** were first determined through preliminary intratumoral injections in B16-F10 tumors. Intratumoral injections of 10 mg/kg **Nico-52** every 48 hours (**Figure 7A**) lead to the reduction of B16-F10 tumors to 6.21% of the vehicle (50% DMSO, 50% 1X PBS) treated tumor volume (**Figure 7B**). Tumor masses measured after excision on the final day were similarly smaller in treated tumors, on average 0.853% of control tumor mass (**Figure 7E**), demonstrating a significant anti-tumor activity when the compound is introduced to the tumor microenvironment.^83^

**Figure 7.**
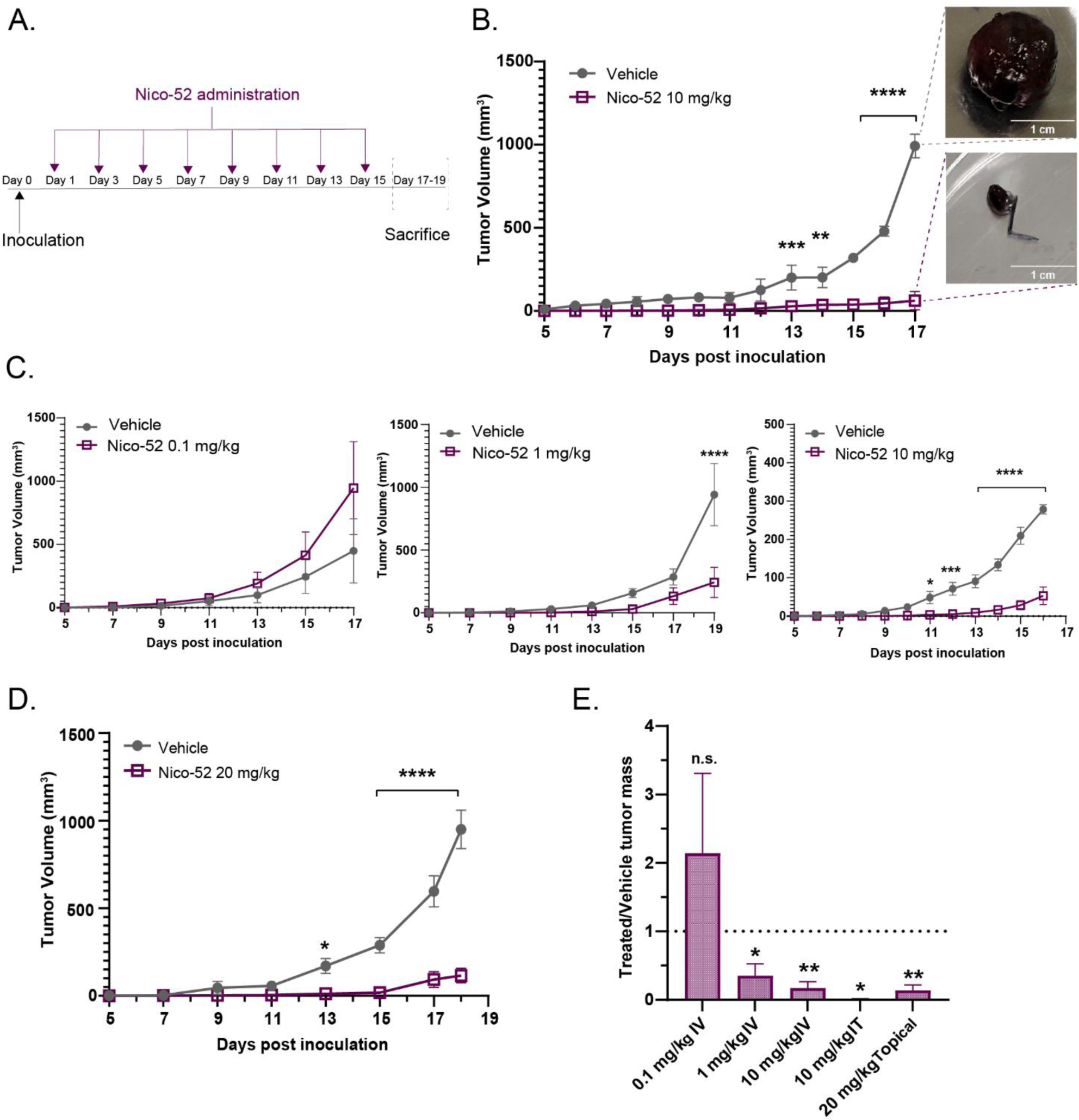
Endpoint study of Nico-52 in B16-F10 tumor-bearing C57BL/6 mice. **(A)** Tumor-bearing C57BL/6 mice were treated with **Nico-52** every 48 hours post-inoculation with B16-F10 cells and sacrificed on day 17-19. **(B)** Intratumoral (IT) injections of 10 mg/kg **Nico-52** every 48 hours starting one day after inoculation suppressed solid tumor growth of B16-F10 tumor-bearing mice (vehicle, n =2, **Nico-52**, n =3) with images of representative tumors at day 17 from each treatment group shown. **(C) Nico-52** intravenous (IV) dose response in B16-F10 tumor-bearing mice via tail vein injection every 48 hours with 0.1 mg/kg (n = 5), 1 mg/kg (n = 5), and 10 mg/kg (n = 3) starting 1 day post inoculation. **(D)** Topical application of 20 mg/kg **Nico-52** every 48 hours inhibited solid tumor growth of B16-F10 tumor-bearing mice (n = 3-4). Estimated tumor volume in **B-D** from digital caliper measurements (0.5 * length * width^2^) and significance determined by two-way ANOVA with Šídák’s multiple comparisons test. **(E)** Ratio of treated: vehicle tumor mass after euthanasia of mice at endpoint and excision of tumors (IT: day 17, 0.1 mg/kg IV: day 17, 1 mg/kg IV: day 19, 10 mg/kg IV: day 16, topical: day 18). Significance determined by student’s unpaired t-test. * = p < 0.05, ** = p < 0.01, *** = p < 0.001, **** = p < 0.0001. *(Intratumoral & intravenous treatment vehicle : 50% DMSO, 50% 1X PBS, Topical treatment vehicle: 100% DMSO). Error bars represent mean ± SEM*.

Intravenous administration of the compound corroborated the intratumoral results. The estimated tumor volumes of B16-F10 tumor-bearing mice were reduced in a dose-dependent manner when treated with intravenous **Nico-52** (**Figure 7C**). Significant reduction in estimated tumor volume was found with 10 mg/kg (19.1% of vehicle (50% DMSO, 50% 1X PBS) treated tumor volume at day 16) and 1 mg/kg doses (25.8% of control volume at day 19), indicating that therapeutic doses of the compound can reach the tumor at those concentrations. However, 0.1 mg/kg doses were insufficient in showing any tumor suppression. Similar results were reflected in the masses of the tumors excised at the end of each corresponding experiment (10 mg/kg: 17.2% of control mass at day 16, 1 mg/kg: 35.1% of control mass at day 19, **Figure 7C**). During the experiments, systemic **Nico-52** injections did not alter the behavior or body weight of mice **(Figure S20)**, suggesting that the compound is tolerated *in vivo*. Rapid body weight loss was sometimes observed as a result of declining health due to tumor burden. Since topical treatment at tumor sites is clinically relevant for melanoma,^84^ we also evaluated if **Nico-52** could be administered dermally. Formulation of **Nico-52** in 100% DMSO at a concentration of 25 mM was applied topically at a dose of 20 mg/kg at the tumor site of B16-F10 tumor-bearing mice (**Figure 7D**). A significant reduction in tumor volume was observed with topical treatment. On day 18, the tumors of treated mice were on average 12.2% of the estimated volume of control tumors (**Figure 7D**). Similarly, the mass of treated tumors on day 18 was also 13.9% of control tumor masses on average. These findings suggest that **Nico-52** can be absorbed through the skin in therapeutic amounts.

### Nico-52 treatment impacts macrophage polarization in the tumor microenvironment

Previous studies utilizing anti-IL-4 antibodies to inhibit tumor growth reported remodeling of the tumor microenvironment by impacting macrophage polarization.^85–87^ To determine if small-molecule IL-4 inhibition impacts the tumor microenvironment in a similar manner, tumors from mice treated with **Nico-52** intratumorally at 10 mg/kg every 48-72 hours were excised at day 17 and analyzed through flow cytometry (**Figure 8A&B, S21**). Macrophages were identified as double positive for both F4/80 and CD11b cellular markers. Macrophages positive for CD206 indicated an M2 phenotype, while those positive for CD80 indicated an M1 phenotype.^88^ The percentage of M1 macrophages was higher (6.17% in control vs. 35.7% in treated) while M2 macrophages were decreased (24.6% in control vs. 2.51% in treated) with intratumoral 10 mg/kg **Nico-52** treatment three times a week for 17 days (**Figure 8A**). This result is consistent with the hypothesis that small-molecule inhibition of IL-4 would prevent M2 macrophage polarization, leading to greater amounts of M1 macrophages and a more immunostimulatory microenvironment.

**Figure 8.**
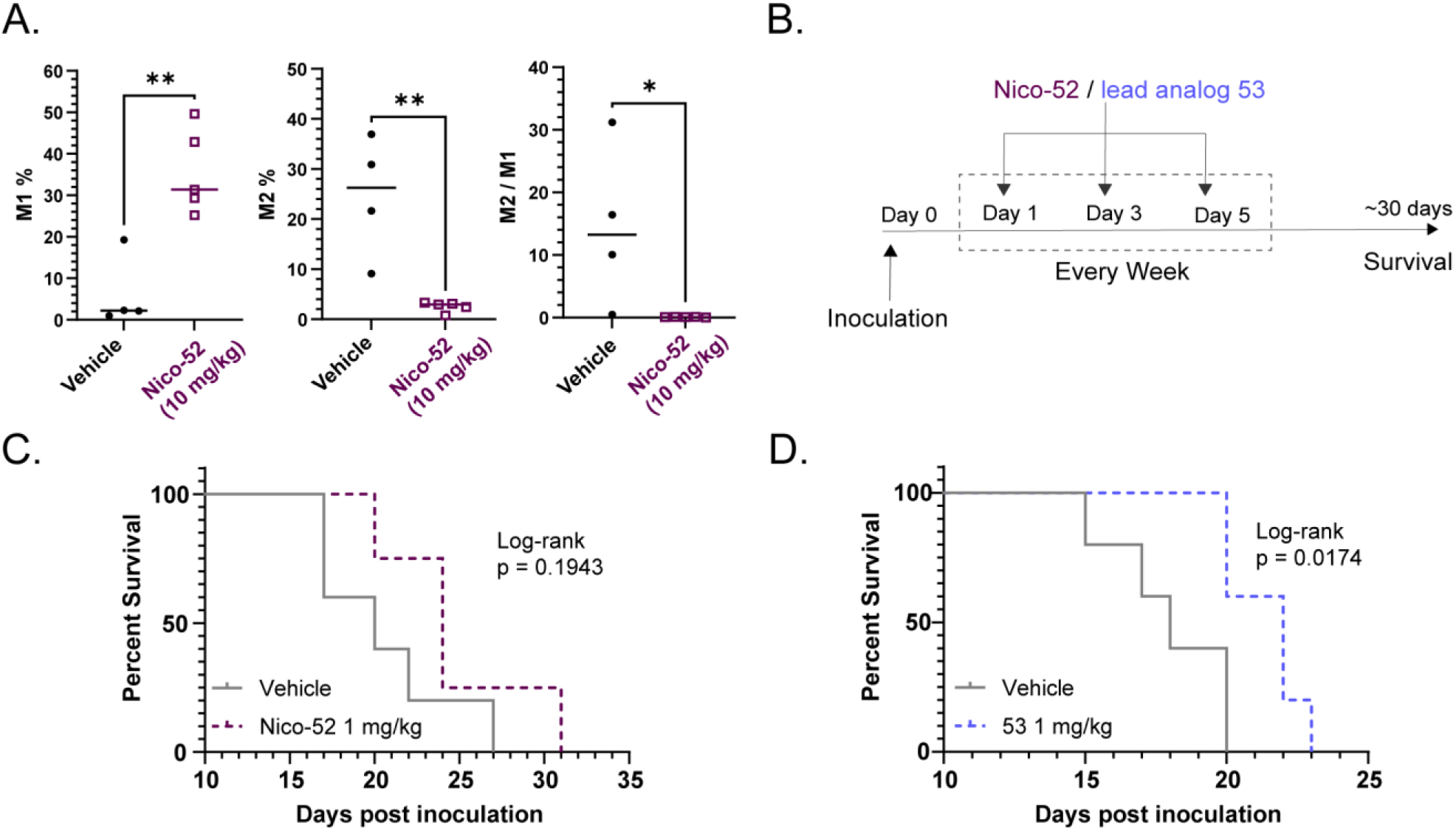
Flow cytometry of dissociated B16-F10 tumors 17 days post-inoculation and survival studies in B16-F10 tumor-bearing C57BL/6 mice using Nico-52 and lead analog 53. **(A)** M1% (p = 0.0025), M2% (p = 0.0044), and M2/M1 ratio (p = 0.0379) from tumors treated with 10 mg/kg **Nico-52** intratumorally three times a week, or vehicle control, collected at 17 days post inoculation (student’s unpaired t-test). *Error bars represent mean ± SEM.* (B) Tumor-bearing C57BL/6 mice were treated thrice a week post-inoculation until week 8 for survival studies. **(C) Nico-52** dosed intravenously (IV) at 1 mg/kg three times a week failed to exhibit increased survival (n=4-5). **(D)** Intraperitoneal (IP) treatment with more potent analog **53** at the same 1 mg/kg doses did lead to significantly increased survival (n=5, p=0.0174, Mantel-Cox test). *(Intravenous treatment vehicle: 50% DMSO, 50% 1X PBS. Intraperitoneal treatment vehicle: 50% PEG-400, 20% DMSO in 1X PBS)*

Together with the lack of *in vitro* cytotoxic anti-cancer activity with these cells (**Table 2**), the flow cytometry results support the expectation that small-molecule IL-4 inhibition acts upon immune-mediated mechanisms for tumor growth/protection in a macrophage-dependent manner.

### Optimized IL-4 inhibitor analog 53 improves survival in the B16-F10 melanoma model

With the anti-cancer activity of the parent compound established, we next assessed if our optimized analog **53** also improves upon **Nico-52’s** activity *in vivo*. Survival studies were performed with **Nico-52** or **53** three times a week for a maximum possible duration of 8 weeks in the B16-F10 model. A lower 1 mg/kg dose was chosen for both compounds since **Nico-52** displayed weaker tumor inhibition at this dose and any increased potency from **53** would be revealed. Mice treated with 1 mg/kg of **Nico-52** appeared to show increased survival for the cohort relative to the vehicle-treated control (50% DMSO, 50% 1X PBS), but it did not achieve statistical significance (**Figure 8C**, p = 0.1943). However, mice treated with the same concentration (1 mg/kg) of the more potent **53** survived a median of 4 days longer than those treated with the vehicle and achieved statistical significance (**Figure 8D**, p=0.0174). This result further supports that more potent IL-4 inhibitors can lead to improved outcomes in this tumor model.

We also assessed if **Nico-52**’s anti-tumor activity would also impact survival for other tumor models. In a 4T1 syngeneic breast cancer model in Balb/C mice (**Figure S22**), mice treated with 10 mg/kg of **Nico-52** intraperitoneally three times a week survived a median of six days longer than those treated with vehicle control (p = 0.0488, 50% DMSO, 50% 1X PBS). The increased survival with **Nico-52** treatment in the 4T1 breast cancer model suggests that the anti-tumor activity of the compound can be effective against other IL-4 dependent cancers.

## DISCUSSION AND CONCLUSIONS

Our SAR studies of **Nico-52** revealed that the R^1^ position is the most amenable to changes and resulted in nanomolar inhibitors **14** and **15**. R^2^ structural changes in **19** and **20** did not result in improved potency. Modifications to the ortho-hydroquinone in the R^3^ position played a significant role in retaining the potency as seen in **43** where complete abolishment of 3,4-diol resulted in almost complete loss of potency. Removal of the fluoro group of **Nico-52** as seen in analog **15** resulted in the largest improvement in potency (71-fold in THP-1 cells), with the unsubstituted phenyl group at this position being more optimal for IL-4 inhibition and maintained good selectivity against IL-13 and other γ_c_-chain utilizing cytokines. While **15** was a more potent IL-4 inhibitor, it did demonstrate a metabolic liability by being vulnerable to oxidation of the phenyl group in a microsomal stability assay. This liability was addressed, however, by utilizing phenyl bioisosteres to improve metabolic stability as seen with the bicyclopentane analog **53** which restored metabolic stability with a potency of 690 nM, positioning the amino nicotinonitrile scaffold for *in vivo* studies. *In vivo* evaluation of **Nico-52** and its lead analogs in syngeneic tumor mouse models revealed tumor-suppressive effects and improved survival. **Nico-52** did not directly affect B16-F10 melanoma cell viability *in vitro* but acted instead through IL-4 dependent immunomodulation of the tumor microenvironment, shifting the balance of tumor-associated macrophages (TAMs) towards the M1 tumor-killing phenotype and away from the immunosuppressive M2 phenotype.

In summary, our studies of the **Nico-52** scaffold revealed insights into designing enhanced inhibitors of the soluble cytokine IL-4. As **Nico-52** was the first reported inhibitor of IL-4, our analogs **14** and **15** are now the first reported nanomolar IL-4 inhibitors. Furthermore, these results contribute to further characterizing and understanding chemotypes capable of inhibiting protein-protein interactions with no defined binding grooves/pockets like the interaction between IL-4 and its receptors.^79^ Blocking IL-4-induced immune responses has improved lung function in mouse models of pulmonary inflammation^89,90^ and impacts macrophage polarization in tumor models.^87^ The development of potent small-molecule inhibitors is a critical step to achieving orally bioavailable therapies that target these cytokine-mediated diseases. Finally, the advent of a small molecule inhibitor of soluble IL-4 with nanomolar potency can complement other cytokine-targeted therapeutic modalities.

## MATERIALS AND METHODS

### General Information for Nico-52 Analog Synthesis

The Perkin Elmer Signal notebook was used for experimental procedure planning. All reactions were performed in 1-dram vials unless otherwise noted. Analytical thin layer chromatography (TLC) was performed using 0.25 mm silica gel 60-F plates (Silicycle, Inc.). All compounds were purified using a CombiFlash automated column from Teledyne ISCO on RediSep Rf or Rf Gold silica Flash columns, 20-40 microns unless otherwise noted.

All ^1^H NMR spectra were recorded at 400 or 500 MHz at ambient temperature with CDCl_3_, CD_3_OD, or DMSO (d_6_) as the solvent. Chemical shifts are recorded in parts per million (ppm) relative to CDCl_3_ (^1^H, δ 7.26; ^13^C, δ 77.1), CD_3_OD (^1^H, δ 3.31; ^13^C, δ 49.0), or (CD_3_)_2_SO (^1^H, δ 2.50; ^13^C, δ 39.5), unless otherwise stated. Data for ^1^H NMR are reported as follows: chemical shift, multiplicity (s = singlet; d = doublet; t = triplet; q = quartet; m = multiplet; br = broad; ovrlp = overlapping), coupling constants (*J* values), and integration. Analytical LCMS was performed on a Waters Acquity UPLC (Ultra Performance Liquid Chromatography (Waters MassLynx Version 4.1) with a Binary solvent manager, SQ mass spectrometer, Water 2996 PDA (PhotoDiode Array) detector, and ELSD (Evaporative Light Scattering Detector).

### Synthesis of Nico-52 and Nico-52 derivatives

In a 4-dram pressure seal vial, a solution of ammonium acetate (963.5 mg, 10 Eq, 12.5 mmol) in 15 mL ethanol was added with R^1^ ketone derivative (1.0 Eq, 1.25 mmol) and stirred for 5 minutes. Then R^3^ aldehyde derivative (1.0 Eq, 1.25 mmol) was added and stirred for 5 minutes. Lastly, malononitrile (82.58 mg, 1.0 Eq, 1.25 mmol) or ethyl 2-cyanoacetate (141.4 mg, 133 μl, 1.0 Eq, 1.25 mmol) was added, and the reaction vial was sealed and heated at 90 °C for 10-14 hours. After completion, the reaction was cooled to room temperature, the vial seal was removed and the stirring stopped. The vial was allowed to slowly evaporate the solvent in the hood until a brown-yellow precipitate formed or until half the solvent remained. If a precipitate formed, filtration was performed and the solid was washed with either cold ethanol or toluene and air dried. If the precipitate was not pure by NMR, recrystallization from either MeOH or 1:10 DMF: EtOH.

If no precipitate formed, the reaction was loaded onto celite in a pre-column cartridge that contained silica gel and placed under vacuum for 48 hours prior to column chromatography. For all column chromatography, 100% DCM was run for 10 minutes, then a 35-minute ramp to 15 percent MeOH, and then a 10-minute flush at 40 percent MeOH was run. UPLC/MS was performed to identify the product and the solvent was evaporated to afford the Nico-52 derivative product. NMR in DMSO-d was used to determine purity and if necessary HPLC was performed for additional purification using an ACN:H_2_O gradient. Nico-52: 4-(3,4-dihydroxyphenyl)-6-(4fluorophenyl)-2-imino-1,2-dihydropyridine-3-carbonitrile. yellow-brown solid, Yield: 13% (150 mg). 1H NMR (500 MHz, DMSO-d) δ 9.47 (s, 1H), 9.27 (s, 1H), 8.20 – 8.14 (m, 2H), 7.74 – 7.64 (m, 1H), 7.35 – 7.27 (m, 2H), 7.18 (s, 1H), 7.09 (d, J = 2.3 Hz, 1H), 7.00 (d, J = 2.2 Hz, 0H), 6.88 (d, J = 8.2 Hz, 3H). MS (EI) m/z: 322 [M + H]^+^.

#### Compound 1

2-amino-4-(3,4-dihydroxyphenyl)-6-(3-fluorophenyl)nicotinonitrile. ^1^H NMR (500 MHz, DMSO) δ 7.94 (m, 2H), 7.55 – 7.47 (m, 1H), 7.32 – 7.27 (t, 1H), 7.23 (s, 1H), 7.08 (s, 1H), 6.99 (d, *J* = 5.9 Hz, 1H), 6.94 (s, 2H), 6.86 (d, J = 8.2 Hz, 1H). UPC2-MS (ES+) m/z calculated for C_18_H_13_FN_3_O_2_, [M + H]^+^ 322.

#### Compound 2

2-amino-4-(3,4-dihydroxyphenyl)-6-(2-fluorophenyl)nicotinonitrile. ^1^H NMR (500 MHz, DMSO) δ 7.92 (td, *J* = 8.0, 1.9 Hz, 1H), 7.55 – 7.47 (m, 1H), 7.38 – 7.27 (m, 2H), 7.04 (d, *J* = 2.2 Hz, 1H), 7.00 – 6.93 (m, 4H), 6.87 (d, *J* = 8.2 Hz, 1H). UPC^2^-MS (ES+) m/z calculated for C_18_H_13_FN_3_O_2_, [M + H]^+^ 322.

#### Compound 3

6-(4-bromophenyl)-4-(3,4-dihydroxyphenyl)-2-imino-2,3-dihydropyridine-3-carbonitrile. yellow-brown solid. ^1^H NMR (500 MHz, CDCL3) δ 8.04 (d, J = 8.6 Hz, 2H), 7.66 (d, J = 8.6 Hz, 2H), 7.18 (s, 1H), 7.05 (d, J = 2.2 Hz, 1H), 6.96 (dd, J = 8.2, 2.3 Hz, 1H), 6.84 (d, J = 8.2 Hz, 1H). MS (EI) m/z: 382 [M + H]^+^.

#### Compound 4

4-(3,4-dihydroxyphenyl)-6-(4-hydroxyphenyl)-2-imino-1,2-dihydropyridine-3-carbonitrile. yellow-brown solid, Yield: 2% (8 mg). ^1^H NMR (500 MHz, DMSO-d) δ 8.01 – 7.95 (m, 2H), 7.14 – 7.06 (m, 2H), 6.99 (dd, J = 8.1, 2.3 Hz, 1H), 6.93 – 6.84 (m, 3H), 6.78 (s, 2H). MS (EI) m/z: 320 [M + H]^+^.

#### Compound 5

4-(3,4-dihydroxyphenyl)-2-imino-6-(p-tolyl)-1,2-dihydropyridine-3-carbonitrile. yellow-brown solid, 6% (22 mg). ^1^H NMR (500 MHz, DMSO-d) δ 9.48 (s, 3H), 8.16 – 8.11 (m, 0H), 8.04 – 7.98 (m, 2H), 7.29 (d, J = 8.0 Hz, 2H), 7.15 (s, 1H), 7.08 (d, J = 2.2 Hz, 1H), 6.98 (dd, J = 8.2, 2.2 Hz, 1H), 6.87 (d, J = 8.5 Hz, 3H), 2.36 (s, 3H). MS (EI) m/z: 318 [M + H]^+^.

#### Compound 6

4-(3,4-dihydroxyphenyl)-2-imino-6-(4-methoxyphenyl)-2,3-dihydropyridine-3-carbonitrile. yellow solid, ^1^H NMR (500 MHz, CDCL_3_) δ 8.05 (d, *J* = 8.9 Hz, 2H), 7.09 (s, 1H), 7.04 (d, *J* = 2.2 Hz, 1H), 7.00 (d, *J* = 8.9 Hz, 2H), 6.94 (dd, *J* = 8.1, 2.2 Hz, 1H), 6.84 (d, *J* = 8.2 Hz, 1H), 3.79 (s, 3H).MS (EI) m/z: 334 [M + H]^+^.

#### Compound 7

4-(3,4-dihydroxyphenyl)-6-imino-1,6-dihydro-[2,3’-bipyridine]-5-carbonitrile. yellow-brown solid Yield: 3% (13 mg).^1^H NMR (500 MHz, DMSO-d) δ 9.41 (d, J = 2.1 Hz, 1H), 8.86 (t, J = 7.9 Hz, 2H), 7.89 (dd, J = 8.1, 5.3 Hz, 1H), 7.41 (s, 1H), 7.30 (s, 1H), 7.22 (d, J = 5.0 Hz, 1H), 7.12 (d, J = 2.0 Hz, 2H), 7.06 – 7.00 (m, 2H), 6.90 (d, J = 8.2 Hz, 1H), 6.78 – 6.61 (m, 2H). MS (EI) m/z: 305 [M + H]^+^.

#### Compound 8

2-amino-4-(3,4-dihydroxyphenyl)-6-(furan-2-yl)nicotinonitrile. Light yellow solid, Yield: 7.09% (30.1mg).^1^H NMR (500 MHz, dmso) δ 7.86 (s, 1H), 7.16 (d, *J* = 3.7 Hz, 1H), 7.02 (d, *J* = 2.1 Hz, 1H), 6.96 (s, 1H), 6.93 (dd, *J* = 8.2, 2.2 Hz, 1H), 6.88 (s, 1H), 6.85 (d, *J* = 8.2 Hz, 1H). MS (EI) m/z: 294 [M + H]^+^.

#### Compound 9

4-(3,4-dihydroxyphenyl)-2-imino-6-(pyrazin-2-yl)-2,3-dihydropyridine-3-carbonitrile. Light yellow solid,Yield: .26% (1.7mg).^1^H NMR (500 MHz, CDCL_3_) δ 7.91 (dd, *J* = 8.6, 5.3 Hz, 2H), 7.34 (t, *J* = 8.7 Hz, 2H), 7.29 (s, 1H), 7.07 (d, *J* = 8.2 Hz, 1H), 6.78 (s, 1H). MS (EI) m/z: 306 [M + H]^+^.

#### Compound 10

6-([1,1’-biphenyl]-4-yl)-4-(3,4-dihydroxyphenyl)-2-imino-1,2-dihydropyridine-3-carbonitrile. yellow-brown solid, Yield: 1% (5 mg).^1^H NMR (500 MHz, CD_3_OD) δ 8.15 – 8.09 (m, 2H), 7.73 – 7.69 (m, 2H), 7.69 – 7.64 (m, 2H), 7.45 (dd, J = 8.4, 6.9 Hz, 3H), 7.38 – 7.32 (m, 1H), 7.20 (s, 1H), 7.14 (d, J = 2.3 Hz, 1H), 7.05 (dd, J = 8.2, 2.2 Hz, 1H), 6.91 (d, J = 8.2 Hz, 1H). MS (EI) m/z: 380 [M + H]^+^.

#### Compound 11

4-(3,4-dihydroxyphenyl)-2-imino-6-(naphthalen-2-yl)-1,2-dihydropyridine-3-carbonitrile.yellow-brown solid,Yield: 1% (4 mg). ^1^H NMR (500 MHz, CD_3_OD) δ 8.59 – 8.49 (m, 1H), 8.20 – 8.10 (m, 1H), 8.02 – 7.84 (m, 4H), 7.54 – 7.48 (m, 2H), 7.30 (s, 1H), 7.16 (d, J = 2.2 Hz, 1H), 7.07 (dd, J = 8.2, 2.2 Hz, 1H), 6.92 (d, J = 8.2 Hz, 1H). MS (EI) m/z: 354 [M + H]^+^.

#### Compound 12

2-amino-6-cyclohexyl-4-(3,4-dihydroxyphenyl)nicotinonitrile. ^1^H NMR (500 MHz, DMSO) δ 9.32 (s, 2H), 6.96 (d, *J* = 2.2 Hz, 1H), 6.89 – 6.78 (m, 2H), 6.66 (s, 2H), 6.47 (s, 1H), 2.48 (d, *J* = 2.3 Hz, 1H), 1.81 – 1.71 (m, 4H), 1.67 (d, *J* = 12.8 Hz, 1H), 1.45 (qd, *J* = 13.0, 3.7 Hz, 2H), 1.36 – 1.25 (m, 2H), 1.23 – 1.13 (m, 1H). UPC^2^-MS (ES+) m/z calculated for C18H19N3O2, [M + H]^+^ 310.

#### Compound 13

2-amino-4-(3,4-dihydroxyphenyl)-6-methylnicotinonitrile. Light yellow solid, 1.65% (10.3 mg). ^1^H NMR (500 MHz, DMSO) δ 6.94 (d, J = 2.1 Hz, 1H), 6.86 (dd, J = 8.4, 2.0 Hz, 1H), 6.82 (d, J = 8.2 Hz, 1H), 6.53 (s, 1H), 2.25 (s, 3H). MS (EI) m/z: 242 [M + H]^+^.

#### Compound 14

6-(4-aminophenyl)-4-(3,4-dihydroxyphenyl)-2-imino-1,2-dihydropyridine-3-carbonitrile. yellow-brown solid, Yield: 3% (11 mg).^1^H NMR (500 MHz, DMSO-d) δ 9.36 (s, 2H), 7.87 – 7.81 (m, 2H), 7.03 (d, J = 2.2 Hz, 1H), 6.98 (s, 1H), 6.94 (dd, J = 8.2, 2.2 Hz, 1H), 6.86 (d, J = 8.2 Hz, 1H), 6.64 (s, 2H), 6.63 – 6.57 (m, 2H), 5.62 (s, 2H). MS (EI) m/z: 319 [M + H]^+^.

#### Compound 15

4-(3,4-dihydroxyphenyl)-2-imino-6-phenyl-1,2-dihydropyridine-3-carbonitrile. yellow-brown solid, Yield: 6% (24 mg). ^1^H NMR (500 MHz, DMSO-d) δ 9.44 (s, 1H), 9.26 (s, 1H), 8.13 – 8.06 (m, 2H), 7.48 (dq, J = 8.8, 3.0, 2.4 Hz, 4H), 7.18 (s, 1H), 7.08 (d, J = 2.2 Hz, 1H), 6.99 (dd, J = 8.2, 2.3 Hz, 1H), 6.89 (s, 2H). MS (EI) m/z: 304 [M + H]^+^.

#### Compound 16

6-(4-(1H-imidazol-1-yl)phenyl)-2-amino-4-(3,4-dihydroxyphenyl)nicotinonitrile. Light yellow solid, Yield: 2.94% (15.7mg). ^1^H NMR (500 MHz, dmso) δ 8.37 (s, 1H), 8.25 (d, *J* = 8.7 Hz, 2H), 7.86 – 7.83 (m, 1H), 7.78 (d, *J* = 8.7 Hz, 2H), 7.25 (s, 1H), 7.13 (t, *J* = 1.1 Hz, 1H), 7.09 (d, *J* = 2.2 Hz, 1H), 6.99 (dd, *J* = 8.2, 2.2 Hz, 1H), 6.87 (d, *J* = 8.1 Hz, 1H). MS (EI) m/z: 370 [M + H]^+^.

#### Compound 17

2-amino-4-(3,4-dihydroxyphenyl)-6-(4-(dimethylamino)phenyl)nicotinonitrile. Light yellow solid, Yield: 4.15% (20.8mg).^1^H NMR (500 MHz, dmso) δ 7.96 (d, *J* = 8.8 Hz, 2H), 6.93 (dd, *J* = 8.2, 2.2 Hz, 1H), 6.85 (d, *J* = 8.2 Hz, 1H), 6.74 (d, *J* = 8.8 Hz, 2H), 6.72 (d, *J* = 2.1 Hz, 1H), 6.69 (s, 1H), 2.96 (s, 6H). MS (EI) m/z: 347 [M + H]^+^.

#### Compound 19

6-(4-fluorophenyl)-2-imino-4-phenyl-1,2-dihydropyridine-3-carbonitrile. yellow-brown solid, Yield: 5% (19 mg). ^1^H NMR (500 MHz, DMSO-d) δ 8.24 – 8.15 (m, 2H), 7.70 – 7.62 (m, 2H), 7.59 – 7.43 (m, 3H), 7.34 – 7.21 (m, 3H), 7.01 (s, 2H). MS (EI) m/z: 290 [M + H]^+^.

#### Compound 20

6-(4-fluorophenyl)-2-imino-4-(5-methylfuran-2-yl)-1,2-dihydropyridine-3-carbonitrile. yellow-brown solid, Yield: 11% (39 mg). ^1^H NMR (500 MHz, DMSO-d) δ 8.18 – 8.10 (m, 2H), 7.44 (s, 1H), 7.41 (d, J = 3.4 Hz, 1H), 7.36 – 7.27 (m, 2H), 6.93 (s, 2H), 6.39 (dd, J = 3.5, 1.1 Hz, 1H), 2.40 (s, 3H). MS (EI) m/z: 294 [M + H]^+^.

#### Compound 21

6-(4-fluorophenyl)-2-imino-4-(5-methylthiophen-2-yl)-1,2-dihydropyridine-3-carbonitrile. yellow-brown solid, Yield: 6% (22 mg). ^1^H NMR (500 MHz, CD_3_OD) δ 7.74 – 7.44 (m, 3H), 7.36 (d, J = 3.7 Hz, 1H), 7.20 (dt, J = 41.7, 8.4 Hz, 2H), 6.88 – 6.62 (m, 2H), 2.42 (d, J = 11.9 Hz, 3H). MS (EI) m/z: [M + H]^+^.

#### Compound 22

6-(4-fluorophenyl)-4-(4-hydroxyphenyl)-2-oxo-1,2-dihydropyridine-3-carbonitrile. yellow solid, Yield: 9% (34 mg). ^1^H NMR (500 MHz, DMSO-d) δ 12.65 (s, 1H), 10.12 (s, 1H), 7.99 – 7.92 (m, 2H), 7.67 – 7.60 (m, 2H), 7.41 – 7.33 (m, 2H), 6.96 – 6.89 (m, 2H), 6.77 (s, 1H). MS (EI) m/z: 307 [M + H]^+^.

#### Compound 23

2-amino-6-(4-fluorophenyl)-4-(2-hydroxyphenyl)nicotinonitrile. yellow-brown solid, Yield: 8% (31 mg). ^1^H NMR (500 MHz, DMSO-d) δ 8.57 (dd, J = 8.1, 1.5 Hz, 1H), 8.42 – 8.33 (m, 2H), 8.01 (s, 1H), 7.66 (ddd, J = 8.4, 7.3, 1.5 Hz, 1H), 7.47 – 7.32 (m, 4H). MS (EI) m/z: 306 [M + H]^+^.

#### Compound 24

6-(4-fluorophenyl)-4-(4-hydroxy-3-methoxyphenyl)-2-imino-1,2-dihydropyridine-3-carbonitrile. yellow-brown solid, Yield: 9% (37 mg). ^1^H NMR (500 MHz, DMSO-d) δ 9.52 (s, 1H), 8.25 – 8.14 (m, 2H), 7.36 – 7.29 (m, 2H), 7.27 (d, J = 1.9 Hz, 2H), 7.22 – 7.10 (m, 1H), 6.98 – 6.85 (m, 4H), 3.86 (s, 3H). MS (EI) m/z: 336 [M + H]^+^.

#### Compound 25

6-(4-fluorophenyl)-4-(3-hydroxy-4-methoxyphenyl)-2-imino-1,2-dihydropyridine-3-carbonitrile. yellow-brown solid, Yield: 7% (28 mg). ^1^H NMR (500 MHz, DMSO-d) δ 8.21 – 8.15 (m, 2H), 7.36 – 7.29 (m, 2H), 7.21 (s, 1H), 7.18 – 7.10 (m, 2H), 7.09 – 7.04 (m, 1H), 6.96 (s, 2H), 3.85 (s, 3H). MS (EI) m/z: 336 [M + H]^+^.

#### Compound 26

6’-(4-fluorophenyl)-2’-imino-1’,2’-dihydro-[3,4’-bipyridine]-3’-carbonitrile. yellow-brown solid,Yield: 6% (21 mg). ^1^H NMR (500 MHz, DMSO-d) δ 8.88 (s, 1H), 8.79 – 8.70 (m, 1H), 8.33 – 8.20 (m, 2H), 8.13 (dt, J = 7.9, 2.0 Hz, 1H), 7.60 (dd, J = 7.9, 4.8 Hz, 1H), 7.39 (d, J = 3.8 Hz, 1H), 7.37 – 7.31 (m, 2H), 7.15 (s, 2H). MS (EI) m/z: 291 [M + H]^+^.

#### Compound 27

5-(3-cyano-6-(4-fluorophenyl)-2-imino-1,2-dihydropyridin-4-yl)-2-hydroxybenzoic acid. yellow-brown solid, Yield: 2% (8.8 mg). ^1^H NMR (500 MHz, DMSO-d) δ 8.43 – 8.35 (m, 2H), 8.14 (s, 1H), 8.22 – 8.03 (m, 1H), 7.39 – 7.30 (m, 3H), 6.99 (dd, J = 122.1, 8.5 Hz, 1H). MS (EI) m/z: 350 [M + H]^+^.

#### Compound 28

4-(3-bromo-4-hydroxyphenyl)-6-(4-fluorophenyl)-2-imino-1,2-dihydropyridine-3-carbonitrile. yellow-brown solid, Yield: 10% (47 mg). ^1^H NMR (500 MHz, DMSO-d) δ 10.82 (s, 1H), 8.24 – 8.16 (m, 2H), 7.85 (d, J = 2.3 Hz, 1H), 7.55 (dd, J = 8.4, 2.3 Hz, 1H), 7.31 (t, J = 8.8 Hz, 2H), 7.10 (d, J = 8.4 Hz, 1H), 7.00 (s, 2H). MS (EI) m/z: 385 [M + H]^+^.

#### Compound 29

6-(4-fluorophenyl)-4-(4-hydroxy-3-iodophenyl)-2-imino-1,2-dihydropyridine-3-carbonitrile. yellow-brown solid, Yield: 8% (43 mg). ^1^H NMR (500 MHz, DMSO-d) δ 10.88 (s, 1H), 9.98 (d, J = 4.4 Hz, 1H), 8.19 (ddd, J = 10.9, 6.6, 3.7 Hz, 2H), 8.01 (d, J = 2.3 Hz, 1H), 7.59 – 7.51 (m, 1H), 7.29 (t, J = 8.8 Hz, 3H), 7.03 (d, J = 8.4 Hz, 1H), 6.98 (s, 2H). MS (EI) m/z: 432 [M + H]^+^.

#### Compound 30

4-(3,4-dihydroxyphenyl)-6-(4-fluorophenyl)-2-oxo-1,2-dihydropyridine-3-carbonitrile. yellow-brown solid, Yield: 9% (37 mg).1H NMR (500 MHz, DMSO-d) δ 12.65 (s, 1H), 9.67 (s, 1H), 9.36 (s, 1H), 7.95 (dd, J = 8.6, 5.3 Hz, 2H), 7.38 – 7.33 (m, 2H), 7.17 (d, J = 2.3 Hz, 1H), 7.09 (dd, J = 8.2, 2.3 Hz, 1H), 6.89 (d, J = 8.3 Hz, 1H), 6.73 (s, 1H). MS (EI) m/z: 323 [M + H]^+^.

#### Compound 31

4-(3,4-dihydroxyphenyl)-6-(4-fluorophenyl)-2-imino-5-methyl-1,2-dihydropyridine-3-carbonitrile. yellow-brown solid, Yield: 7% (29 mg).1H NMR (500 MHz, DMSO-d) δ 7.56 (ddd, J = 8.9, 5.6, 2.7 Hz, 2H), 7.33 – 7.22 (m, 2H), 6.84 (d, J = 8.0 Hz, 1H), 6.71 (d, J = 2.2 Hz, 1H), 6.67 (s, 2H), 6.61 (dd, J = 8.1, 2.1 Hz, 1H), 1.85 (s, 3H). MS (EI) m/z: 336 [M + H]^+^.

#### Compound 42

6-([1,1’-biphenyl]-4-yl)-4-(3,4-dihydroxyphenyl)-2-oxo-2,3-dihydropyridine-3-carbonitrile. yellow-brown solid. 1H NMR (500 MHz, CDCL3) δ 7.97 (d, J = 8.0 Hz, 2H), 7.82 (d, J = 8.6 Hz, 1H), 7.75 (d, J = 7.1 Hz, 1H), 7.49 (dd, J = 8.4, 7.0 Hz, 2H), 7.44 – 7.37 (m, 1H), 7.17 (d, J = 2.3 Hz, 1H), 7.09 (dd, J = 8.2, 2.3 Hz, 1H), 6.88 (d, J = 8.2 Hz, 1H), 6.76 (s, 1H). MS (EI) m/z: 381 [M + H]^+^.

#### Compound 43

5-(2-amino-6-(4-aminophenyl)-3-cyanopyridin-4-yl)-2-hydroxybenzoic acid. ^1^H NMR (500 MHz, DMSO) δ 7.95 (s, 1H), 7.84 (d, *J* = 8.7 Hz, 2H), 7.45 (d, *J* = 8.4 Hz, 1H), 6.98 (s, 1H), 6.72 (d, *J* = 10.0 Hz, 1H), 6.63 – 6.57 (m, 4H), 5.61 (s, 2H). UPC^2^-MS (ES+) m/z calculated for C_19_H_14_N_4_O_3_, [M + H]^+^ 347.

#### Compound 44

4-(3,4-dichlorophenyl)-6-(4-fluorophenyl)-2-oxo-1,2-dihydropyridine-3-carbonitrile. yellow solid, Yield: 23% (101 mg).^1^H NMR (500 MHz, DMSO-d) δ 12.92 (s, 1H), 8.05 (d, J = 2.1 Hz, 1H), 8.00 (t, J = 6.9 Hz, 2H), 7.86 (d, J = 8.4 Hz, 1H), 7.74 (dd, J = 8.4, 2.2 Hz, 1H), 7.45 – 7.35 (m, 2H), 6.93 (s, 1H). MS (EI) m/z: 360 [M + H]^+^.

#### Compound 45

6-(4-fluorophenyl)-2-oxo-4-phenyl-1,2-dihydropyridine-3-carbonitrile. yellow solid, Yield: 14% (50 mg). ^1^H NMR (500 MHz, DMSO-d) δ 12.82 (s, 1H), 7.99 (ddd, J = 8.5, 5.3, 2.4 Hz, 2H), 7.74 (dt, J = 7.5, 2.6 Hz, 2H), 7.57 (q, J = 2.8 Hz, 3H), 7.38 (td, J = 8.9, 2.4 Hz, 2H), 6.95 – 6.75 (m, 1H). MS (EI) m/z: 291 [M + H]^+^.

#### Compound 46

6-(4-fluorophenyl)-2-imino-4-(tetrahydro-2H-pyran-3-yl)-1,2-dihydropyridine-3-carbonitrile. yellow-brown solid, Yield: 2% (8 mg). ^1^H NMR (500 MHz, DMSO-d) δ 12.66 (s, 1H), 7.91 (s, 2H), 7.45 – 7.32 (m, 2H), 6.80 (s, 1H), 3.95 – 3.78 (m, 2H), 3.45 (td, J = 10.9, 3.9 Hz, 1H), 3.09 – 2.96 (m, 1H), 2.05 – 1.85 (m, 2H), 1.74 – 1.57 (m, 2H). MS (EI) m/z: 298 [M + H]^+^.

#### Compound 47

6-(4-fluorophenyl)-4-(5-methylfuran-2-yl)-2-oxo-1,2-dihydropyridine-3-carbonitrile. yellow solid, Yield: 10% (36 mg). ^1^H NMR (500 MHz, DMSO-d) δ 12.51 (s, 1H), 7.99 – 7.83 (m, 2H), 7.61 (d, J = 3.5 Hz, 1H), 7.47 – 7.30 (m, 2H), 6.96 (s, 1H), 6.48 (dd, J = 3.6, 1.1 Hz, 1H), 2.42 (s, 3H). MS (EI) m/z: 295 [M + H]^+^.

#### Compound 48

6-(4-fluorophenyl)-2-imino-4-(naphthalen-2-yl)-1,2-dihydropyridine-3-carbonitrile. yellow-brown oil, Yield: 7% (31 mg). ^1^H NMR (500 MHz, DMSO-d) δ 12.85 (s, 1H), 8.33 (d, J = 1.8 Hz, 1H), 8.10 (d, J = 8.6 Hz, 1H), 8.08 – 7.98 (m, 4H), 7.83 (dd, J = 8.5, 1.9 Hz, 1H), 7.64 (tt, J = 6.9, 5.2 Hz, 2H), 7.47 – 7.30 (m, 2H), 6.98 (s, 1H). MS (EI) m/z: 340 [M + H]^+^.

#### Compound 49

6-(4-fluorophenyl)-2-hydroxy-4-(2-hydroxyphenyl)nicotinonitrile. yellow solid, Yield: 21% (79 mg). ^1^H NMR (500 MHz, DMSO-d) δ 8.56 (dd, J = 8.1, 1.5 Hz, 1H), 8.41 – 8.33 (m, 2H), 8.01 (s, 1H), 7.73 – 7.62 (m, 1H), 7.47 – 7.33 (m, 4H). MS (EI) m/z: 307 [M + H]^+^.

#### Compound 50

6-(4-fluorophenyl)-4-(3-hydroxyphenyl)-2-oxo-1,2-dihydropyridine-3-carbonitrile. yellow solid, Yield: 11% (43 mg). ^1^H NMR (500 MHz, DMSO-d) δ 12.78 (s, 1H), 9.83 (s, 1H), 7.97 (dd, J = 8.6, 5.3 Hz, 2H), 7.36 (q, J = 8.3, 7.8 Hz, 3H), 7.12 (dt, J = 7.7, 1.3 Hz, 1H), 7.08 (t, J = 2.1 Hz, 1H), 6.95 (ddd, J = 8.2, 2.5, 0.9 Hz, 1H), 6.82 – 6.78 (m, 1H). MS (EI) m/z: 307 [M + H]^+^.

#### Compound 51

6-(4-fluorophenyl)-4-(4-hydroxyphenyl)-2-oxo-1,2-dihydropyridine-3-carbonitrile. yellow solid, Yield: 9% (34 mg). ^1^H NMR (500 MHz, DMSO-d) δ 12.65 (s, 1H), 10.12 (s, 1H), 7.99 – 7.92 (m, 2H), 7.67 – 7.60 (m, 2H), 7.41 – 7.33 (m, 2H), 6.96 – 6.89 (m, 2H), 6.77 (s, 1H). MS (EI) m/z: 307 [M + H]^+^.

#### Compound 52

6-(4-fluorophenyl)-4-(4-hydroxy-3-methoxyphenyl)-2-oxo-1,2-dihydropyridine-3-carbonitrile. yellow solid, Yield: 10% (42 mg). ^1^H NMR (500 MHz, DMSO-d) δ 12.65 (s, 1H), 9.72 (s, 1H), 8.00 – 7.92 (m, 2H), 7.42 – 7.32 (m, 3H), 7.24 (dd, J = 8.2, 2.2 Hz, 1H), 6.93 (d, J = 8.2 Hz, 1H), 6.83 (s, 1H), 3.86 (s, 3H). MS (EI) m/z: 337 [M + H]^+^.

#### Compound 53

6-(bicyclo[1.1.1]pentan-1-yl)-4-(3,4-dihydroxyphenyl)-2-imino-2,3-dihydropyridine-3-carbonitrile. Light yellow solid, Yield: 7.4% (10.0mg). ^1^H NMR (500 MHz, CDCL_3_) δ 7.91 (s, 2H), 7.54 (dd, *J* = 8.6, 1.7 Hz, 2H) 7.31 (dd, *J* = 2.0, 0.3 Hz, 1H), 7.02 (dd, *J* = 8.84, .5 Hz, 1H), 7.05 (sept, *J* = 3.2 Hz, 1H), 2.5-1.7 (m, 6H). MS (EI) m/z: 294 [M + H]^+^.

#### Compound 54

2-amino-6-cyclopropyl-4-(3,4-dihydroxyphenyl) nicotinonitrile. Light yellow solid, Yield: 33.1% (128mg). 1H NMR (500 MHz, dmso) δ 6.95 (d, J = 2.1 Hz, 1H), 6.85 (dd, J = 8.1, 2.1 Hz, 1H), 6.82 (d, J = 8.2 Hz, 1H), 6.55 (s, 1H), 1.98 (p, J = 6.4 Hz, 1H), 0.93 (s, 2H), 0.92 (s, 2H). MS (EI) m/z: 268 [M + H]^+^.

#### Compound 55

2-amino-6-cyclobutyl-4-(3,4-dihydroxyphenyl) nicotinonitrile. Light yellow solid, Yield: 7.44% (30.3mg). ^1^H NMR (500 MHz, dmso) δ 6.95 (d, *J* = 2.1 Hz, 1H), 6.85 (d, *J* = 2.2 Hz, 1H), 6.83 (s, 1H), 6.46 (s, 1H), 3.49 (p, *J* = 8.7 Hz, 1H), 2.24 – 2.16 (m, 4H), 2.01 – 1.90 (m, 2H). MS (EI) m/z: 282 [M + H]^+^.

### Reagents

Tag-free HEK-293 expressed IL-4 for HEK-Blue IL-4/IL-13, THP-1, and Ramos inhibition assays purchased from Acros Biosystems (IL4-H4218). Tag-free mouse IL-4 (574302) for Raw 264.7 macrophage Inhibition assay was purchased from BioLegend.

Blasticidin, Zeocin, Normocin, and Quanti-Blue were purchased from Invivogen. pSTAT6 (690102) and STAT6 (657901) antibodies for THP-1 and Ramos Inhibition assays were purchased from BioLegend. pSTAT6 antibody (PI700247) for Raw Macrophage Inhibition assay was purchased from Fisher Scientific. All other chemicals and reagents were purchased from Fisher Scientific and Sigma Aldrich.

### HEK -Blue IL-4/IL-13 Inhibition assay

HEK-Blue IL-4/IL-13 reporter cell lines were purchased from Invivogen and cultured in DMEM with 10% Heat Inactivated Fetal Bovine Serum (FBS), 100U/ml-100µg/ml Pen-Strep, 10 µg/ml blasticidin, 100 µg/ml Normocin and 100 µg/ml Zeocin. Cells were tested for SEAP production induced by IL-4 alone prior to small molecule screening. It was determined to be 0.3 ng/ml. The EC_25_ for IL-4 induced SEAP production (0.1 ng/ml final well concentration) was chosen for screening small molecules to determine their inhibitory activity.

22.5 µl of small molecule in DMEM 4% DMSO was added to 22.5 µl of 1 ng/ml IL-4 in DMEM in a non-binding 96-well reaction plate for 30 minutes at room temperature. The reaction plate was further incubated for 15 min at 37_°_C. The HEK-Blue IL-4/IL-13 reporter cell line was used between passages 11-17. Cells were washed with 1X PBS, trypsinized using 0.25% Trypsin-EDTA (1X) for 2 minutes, and centrifuged for 5 minutes at 250 x g. Cells were resuspended in DMEM 10% FBS and 100 U/ml-100 µg/ml Pen-Strep at a concentration of 3.1×10^5^ cells/ml. 160 µl of cell suspension was added to each well of a 96-well plate, avoiding edges, and incubated for 30 minutes in a 37_°_C/5% CO_2_ incubator. Lastly, 40 µl of IL-4 with small molecule/ vehicle (DMEM, 0.4% DMSO final concentration) solution was added to the cells preincubated in a 96-well plate. Measurements were performed in triplicate. The plate was incubated for ∼22-24 hours at 37_°_C/5% CO_2_. Quanti-Blue dye was prepared as per the manufacturer’s recommendation and 160 µl was dispensed in each well of a 96-well plate (avoid edges to avoid evaporation), 40 µl of cell supernatant was added to the same plate and incubated for 3 hours at 37_°_C/5% CO_2_. Optical density was then measured at 650 nm using Molecular Devices SpectraMax M5 Microplate reader. Data was analyzed and plotted using GraphPad Prism 9.2.0. Hill coefficients for Nico-52, 14, and 15 were also determined using the dose-response evaluation in the HEK-Blue IL-4/IL-13 reporter cell line using GraphPad Prism 9.2.0.

HEK-Blue IL-4/IL-13 reporter cell lines were also used to screen potent Nico-52 analogs against IL-13 similarly. Prior to screening, IL-13 induced SEAP production was quantified with IL-13 concentrations ranging from 0.2-100 µg/ml with 0.04% DMSO in DMEM (final well concentration). 8 ng/ml of IL-13 was incubated with increasing concentrations of analog 14 and 15 with 4% DMSO in DMEM for 30 minutes at room temperature and 15 minutes at 37_°_C. The solution was added to preincubated cells in a 96-well culture plate. The supernatant was then incubated with Quanti-Blue dye for 3 hours and absorbance was measured at 650 nm. Data was analyzed and plotted using GraphPad Prism 9.2.0 .

### Differential Scanning Fluorimetry

Potent Nico-52 analogs were tested for selectivity across cytokines IL-2, IL-7, IL-21 and IL-13. 1 µg of cytokine in PBS was added to each well in 384-well plates followed by the addition of 8X Spiro-Orange dye proceeded with PBS, DMSO, or potent analogs. Potent analogs were screened at a final concentration of 2 µM of small molecule in 0.2% DMSO. DSF was performed using a BioRad CFX384. Melting temperature shifts were calculated by subtracting the nadir of the melting curve of cytokine with the vehicle (1XPBS, 0.2% DMSO) from the nadir of the melting curve of the cytokine with potent analogs. A cut-off of >0.5_°_C was used for binding determination. Plots were made and analyzed using GraphPad Prism 9.2.0.

### THP-1 STAT-6 Phosphorylation assay

THP-1 cells were received as a gift from the laboratory of Dr. Mark Grinstaff and maintained in RPMI-1640 with 10% heat-inactivated fetal bovine serum and 50 µM beta-mercaptoethanol. Cells used for assays were between passages 6 and 10. 10 ng/mL IL-4 was preincubated with compound (vehicle (RPMI-1640, 2% DMSO) or Nico-52 potent analogs) for 30-45 minutes at 37°C. 1 mL of each solution was then added to a pellet of 2×10^6^ cells/ml THP-1 cells and pipetted up and down. THP-1 cells were incubated for 30 minutes in the incubator at 37 °C/5% CO_2_. Cells were then spun down and lysed with 200 µL ice-cold RIPA buffer complete with phosphatase and protease inhibitors. Cell lysates were incubated on ice for 30 minutes, spun down and the supernatants were stored at -20°C until western blot analysis. Undiluted cell lysates were run on SDS-PAGE and transferred to Immunoblot PVDF membrane using Biorad equipment. Membranes were incubated with 1:1000 primary antibody overnight at 4 °C. Membranes were then incubated with 1:50,000 dilution secondary for 1 hour at room temperature. Membranes were then exposed to Femto ECL for 5 minutes and imaged using a Bio-Rad Gel Doc. The brightness/contrast was adjusted in FIJI and the area under the curve of each lane was quantified. The phosphorylated STAT-6 to STAT-6 ratio was calculated and normalized to the amount of total STAT-6. The data was plotted and analyzed using GraphPad Prism 9.2.0.

### Ramos STAT-6 Phosphorylation assay

Ramos cells were purchased from ATCC and maintained in RPMI-1640 with 10% heat-inactivated fetal bovine serum. Cells used for assays were between passages 4 and 8. Cell lysate for IL-4 or IL-4 and Nico-52/Potent analogs treated cells were generated similarly to the THP-1 Inhibition assay. Undiluted cell lysates were run SDS-PAGE and transferred to Immunoblot PVDF membrane using Biorad equipment. Membranes were incubated with 1:1000 primary antibody overnight at 4°C. Membranes were then incubated with 1:50,000 dilution secondary for 1 hour at room temperature. Membranes were then exposed to Femto ECL for 5 minutes and imaged using a Bio-Rad Gel Doc. The brightness/contrast was adjusted in FIJI and the area under the curve of each lane was quantified. The phosphorylated STAT-6 to STAT-6 ratio was calculated and normalized to the amount of total STAT-6. The data was plotted and analyzed using GraphPad Prism 9.2.0.

### RAW 264.7 STAT-6 Phosphorylation assay

RAW 264.7 cells were purchased from ATCC and maintained in DMEM with 10% heat-inactivated FBS. Cells used for assays were between passages 3 and 5. Cell lysates for IL-4 or IL-4 and Nico-52 treated cells were generated similarly to the THP-1 Inhibition assay, but murine IL-4 was used at a concentration of 5 ng/ml. The undiluted cell lysates were run SDS-PAGE followed by transfer to Immunoblot PVDF membrane using Biorad equipment. Membranes were incubated with 1:1000 primary antibody overnight at 4 °C. Membranes were then incubated with 1:50,000 dilution secondary for 1 hour at room temperature. Membranes were then exposed to Femto ECL for 5 minutes and imaged using a Bio-Rad Gel Doc. The brightness/contrast was adjusted in FIJI and the area under the curve of each lane was quantified. The phosphorylated STAT-6 to STAT-6 ratio was calculated and normalized to the amount of total STAT-6. The data was plotted and analyzed using GraphPad Prism 8.

### THP-1 pSTAT-6 Immunofluorescence

IL-4 and potent analogs or vehicle (RPMI-1640 with 2% DMSO) were preincubated for 30-45 minutes at 37 °C. After the incubation, IL-4 or IL-4/potent analogs were incubated with 2×10^6^ cells/ml THP-1 cells for 30 min at 37°C/5% CO_2_. Incubated cells were centrifuged for 5 min at 250 x g. After aspirating the supernatant, cells were resuspended in 1X PBS and plated into a MatTek dish coated with Cell Tak adhesive for 30 minutes at room temperature. Cells were fixed with 4% Paraformaldehyde for 1 hour at room temperature. Cells were rinsed twice with 1X PBS and then permeabilized with 0.5% Triton-X for 1 hour at room temperature. Cells were rinsed again with 1X PBS, and blocked for an hour in superblock with 0.05% Tween-20, rinsed twice, and then incubated with anti-pSTAT-6 1:100 diluted in superblock 0.05% Tween-20 overnight at 4 °C (parafilmed to avoid evaporation). Cells were rinsed thrice with 1X PBS and then incubated with Mouse anti-rabbit IgG(H+L) Secondary Antibody-Alexa Fluor 647 for 1 hour at room temperature in the dark. Cells were rinsed thrice with 1X PBS and then a dilution of 1:100 Hoechst 33342 dye was added to the cells for 15-20 minutes at room temperature in the dark. Cells were rinsed twice with 1X PBS and left in 1 ml of 1X PBS. Confocal Imaging was done using an Olympus FV10i and images were processed using FIJI.

### UPLC-MS analysis

UPLC-MS analysis was performed on the Waters ACQUITY system using an ACQUITY UPLC 1.7 μm column and equipped with ELS and diode array detection. Prior to analysis, equal volumes of acetonitrile were added to samples of Nico-52 and its analogs, with centrifugation and filtration of hepatic microsomes and plasma before injection. Samples were run on a 10% acetonitrile/90% water-99% acetonitrile/1%water gradient over 2 minutes. Peaks of total absorbance corresponding to Nico-52 or its analogs were identified by their corresponding masses and quantified by the area under the curve.

### Hepatic Microsomes

0.5 mg/mL Mouse (CD-1) liver microsomes purchased from GIBCO MSMCPL were incubated with 50 μM of each compound from 100 mM 100% DMSO stock solutions (10mM in 20% DMSO, 50% PEG-400 stock solution for analog 53) at 37°C/5% CO2 with 1 mM NADPH for 0 (control) or 60 minutes, and 60 minutes without NADPH (NADPH independent metabolism) in PBS before quenching with equal volume of acetonitrile and analyzed by UPLC-MS, quantified by AUC of total absorbance with stability reported as % remaining compared to control.

### Plasma Stability

Compounds at 50 μM from 100mM 100% DMSO stock solutions were incubated for 0 (control) or 180 minutes in plasma from C57BL/6 mouse or pooled human both purchased from Valley Biomedical at 37 °C/5% CO_2_ before quenching with equal volume of acetonitrile and analyzed by UPLC-MS, quantified by AUC of total absorbance with plasma stability reported as % remaining compared to control.

### PAMPA membrane permeability

PVDF membranes purchased from Bioassay Systems PAMPA-096 were coated with 5 µL of 4% Phosphatidylcholine (Lecithin, MP Biomedicals 0210214780) in dodecane. Donor chambers were filled with 300 µL of the compound at 100 μM from 100 mM 100% DMSO stock solutions (10 mM in 20% DMSO, 50% PEG-400 stock solution for analog 53) in PBS, acceptor chambers were filled with an equal volume of PBS, and then the plate was incubated for 18 hours at 37 ^0^C/ 5% CO2 before quenching with an equal volume of acetonitrile and analyzed by UPLC-MS, quantified by AUC of total absorbance. Permeability was calculated by P_e_ = C x ln (1 – OD_A_/OD_E_) cm/s, where OD_A_ is the absorbance of the acceptor solution minus blank, and OD_E_ the absorbance of the equilibrium standard minus blank, and C = 7.72 x 10^-6^ using an 18 hour incubation time.

### *In vitro* Cytotoxicity and Growth Inhibition

CellTiter-Glo 2.0 purchased from Promega G9242) was used to measure cell viability with the Victor-3 plate reader (PerkinElmer) after 24 hours in concentrations of compound up to 25 μM from 100 mM 100% DMSO stock solutions (10 mM in 20% DMSO, 50% PEG-400 stock solution for analog 53) in DMEM (ATC 30-2002) supplemented with 10% FBS (Corning 35011CV). Vehicle was used as a negative control for both cytotoxicity and growth inhibition, methanol was used as a positive control for cytotoxicity, and 1% FBS was used as a positive control for growth inhibition.

### Mouse models of tumor growth

Female C57BL/6 and Balb/C mice (Jackson Labs) between 4-8 weeks of age at the start of each experiment were used. The abdomens of mice were also shaved with a small electric razor using one-handed manual restraint to allow visualization of the skin and tumor growth. B16-F10 cells (ATCC CRL-6475) were grown in DMEM (ATCC 30-2002) supplemented with 10% FBS (Corning 35011CV) between passages 6 and 10. 4T1 (ATCC CRL-2539) were grown in RPMI (ATCC 30-2001) supplemented with 10% FBS (Corning 35011CV) between passages 3-6.

To prepare inoculations, trypsinized B16-F10 or 4T1 cells were resuspended in sterile saline at a concentration of 1×10^5^ cells in 100 μL of sterile saline. The abdominal inoculation site, between the midline and the hindlegs, was sterilized by wiping with sterile isopropanol prep pads. One-handed manual restraint was employed for inoculation. Needles were inserted bevel up from the posterior end of the mouse at a shallow angle for subcutaneous injection. Visualization of the needle under the skin was confirmed before injection into the abdomen of C57BL/6 (B16-F10) or Balb/C (4T1) mice. “Blebbing” of the mouse skin due to injection confirmed successful subcutaneous inoculation.

### *In vivo* Nico-52 treatment

Working solutions of Nico-52 was formulated in 50% DMSO, 50% 1X PBS for *in vivo* injections. 10 mg/kg, 1 mg/kg, and 0.1 mg/kg doses used working concentrations of 25 mM, 2.5 mM, and 250 μM, respectively. 53 was formulated at 1 mM in a 50% PEG-400, 20% DMSO solution in 1X PBS. Mice were weighed with a scale prior to injection to determine dose. Heating the working solution with either a heat lamp, heat gun, or heating block, was required to solubilize the 25 mM Nico-52 and 1 mM 53 solutions with visual confirmation. These would remain in solution for enough time after cooling for injections.

Injections were performed either intratumorally (directly into the tumor site), by intravenous tail vein injections, or intraperitoneal injections no more frequently than every 48 hours starting the first day after inoculation. Injection sites were sterilized with sterile isopropanol prep pads prior to injection. Intratumoral and Intraperitoneal injections were performed in conjunction with one-handed manual restraint, with the dosing needle bevel up at a 45-degree angle. For intraperitoneal injections, the anterior of the mouse was held 45 degrees lower than their posterior in order to move internal organs away from the injection site in the lower abdomen. Intravenous injections were performed after warming the mice under a heat lamp to dilate the tail vein (10-20 minutes). With tail vein dilation confirmed, mice were restrained with the Tailveiner mechanical restraint (Braintree Scientific). Tails were held in one hand with the thumb applying light pressure to the vein proximal to the injection site in order to aid with venous injection. Needles were inserted into the vein bevel up at a near parallel angle. For all injection routes, the needle was removed immediately if the mouse reacted strongly enough to interfere with injection. Otherwise, the injection proceeded with depressing the plunger with the non-restraining hand.

For the topical formulation, a solution of 25 mM Nico-52 in 100% DMSO was used at 20 mg/kg. Mice were restrained with one-handed manual restraint and treated by gently pipetting and massaging the formulation onto the skin over the tumor site every 48 hours starting the first day after inoculation. Restraint was maintained for at least 2 minutes to allow for sufficient absorption before releasing the mice.

For non-survival studies, all mice were euthanized when any mouse in the study required interventional euthanasia.

### Monitoring and interventional euthanasia

In conjunction with dosing, mice were weighed on a scale, their tumors measured, and their behavior assessed. Digital calipers were utilized to measure tumor length and width during the course of the studies in conjunction with dosing. B16-F10 tumors first appear as circular melanated freckles, which grow into larger ovals, then raised solid tumors. 4T1 tumors first appear as raised bumps, with some red discoloration. Tumor volumes were estimated using the formula V = 0.5 * L * W^2^. In cases where multiple primary tumors presented on an individual mouse, each tumor was measured and their volumes summed. When multiple tumors merged into larger tumors, they were measured as usual.Inoculated mice were monitored and humanely euthanized if tumor burden led to adverse conditions indicating tumor metastasis or organ failure. Symptoms that would lead to euthanization included substantial ulceration of tumors, signs of severe pain (hunched posture), altered behavior (low activity/responsiveness, aggressiveness), impaired mobility, or 20% weight loss over 72 hours. Tumors exceeding 2 cm in length also required interventional euthanasia. Primary euthanasia was performed by CO_2_ gas asphyxiation at a flow rate displacing 30-70% of cage volume per minute until cessation of breathing was observed. Secondary euthanasia was performed by cervical dislocation.

### Tumor excision

Tumors were excised after euthanasia for measuring mass, flow cytometry, or qPCR analysis. To excise a tumor, an excision was made with surgical scissors in the abdominal skin near the tumor site. The skin was then pulled with forceps and inverted over the tumor, revealing it. Surgical scissors were used to separate the tumor from the skin and peritoneum and collected in 6 or 12 well plates. With looser, less dense tumors (often observed with B16-F10), scraping the tumor off of the skin with forceps into the plates was required.

### Flow cytometry

Tumors excised after *in vivo* experiments were collected and dissociated into single cell suspensions using Collagenase IV (0.25 mg/ml, Roche), DNAse (0.1 mg/ml, Roche) and strained using a 40 micron cell strainer (Greiner). Cells were then suspended in PBS with 1% FBS at a cell density of 1M cells per 100 μL and blocked with 1 μg/1M anti-CD16/32 antibody (Clone 93, Invitrogen) for 15 minutes at room temperature. Cells were then stained with the following labeled antibodies at 1 μg/1M for 30 minutes at room temperature: F4/80-PE-Cyanine7 (clone BM8, Invitrogen), CD11b-FITC (clone M1/70, BD Biosciences), CD80-PE (clone 16-10A1, Invitrogen), and CD206-APC (clone MR6F3, Invitrogen). Flow cytometry was performed using the FACSCalibur (BD Biosciences) with a flow rate between 300-1000 events per second.

### Statistical analysis for *in vitro* work

GraphPad Prism 9.2.0 software was used for statistical tests of significance. Significance for IL-4 percent inhibition in Ramos cells by Nico-52, analog 14 and 15 for each concentration was determined by one-way ANOVA with Dunnett’s multiple comparison test. * = p < 0.05, ** = p < 0.01, **** = p < 0.0001, ns = not significant.

### Statistical analysis for *In-vivo* work

GraphPad Prism 9.5.1 software was used for statistical tests of significance. Significance for changes in estimated tumor volume with treatment were determined by two-way ANOVA with Šídák’s multiple comparisons test of the means of tumor volume in each treatment group at the different time points of measurement. Significance for changes in tumor mass, M1, M2, or M2/M1 ratios were determined by student’s unpaired t-test. Significance for survival studies was determined by Log-rank (Mantel-Cox) test for significance. * = p < 0.05, ** = p < 0.01, *** = p < 0.001, **** = p < 0.0001.

## ASSOCIATED CONTENT

### List of Supplemental Materials

**Figure S1.** Nico-52 Analog’s Synthesis Scheme for Investigating Structure-Activity Relationships

**Figure S2.** Analog 15 selectivity evaluation in HEK Blue IL-2 reporter cell line

**Figure S3.** pSTAT-6 Western blot for THP-1 cells for analog 14

**Figure S4.** STAT-6 Western blot for THP-1 cells for analog 14

**Figure S5**. pSTAT-6 Western blot for THP-1 cells for analog 15

**Figure S6**. STAT-6 Western blot for THP-1 cells for analog 15

**Figure S7.** pSTAT-6 Western blot for Ramos cells for Nico-52

**Figure S8.** STAT-6 Western blot for Ramos cells for Nico-52

**Figure S9.** pSTAT-6 Western blot for Ramos cells for analogs 14 and 15

**Figure S10.** STAT-6 Western blot for Ramos cells for analogs 14 and 15

**Figure S11.** pSTAT-6 Western blot for Ramos cells for Nico-52

**Figure S12.** STAT-6 Western blot for Ramos cells for Nico-52

**Figure S13.** pSTAT-6 Western blot for Raw macrophage cells for Nico-52

**Figure S14.** STAT-6 Western blot for Raw macrophage cells for Nico-52

**Figure S15.** pSTAT-6 Western blot for Raw macrophage cells for Analog 14

**Figure S16.** STAT-6 Western blot for Raw macrophage cells for Analog 14

**Figure S17.** pSTAT-6 Western blot for Raw macrophage cells for Analog 15

**Figure S18.** STAT-6 Western blot for Raw macrophage cells for Analog 15

**Figure S19.** Lead analog 53 dose response in HEK Blue IL-4/IL-13 reporter assay

**Figure S20.** Body weight of C57BL/6 mice treated with Nico-52 10 mg/kg intravenously for endpoint study

**Figure S21.** Flow cytometry gating of dissociated B16-F10 tumors 17 days post-inoculation

**Figure S22.** Survival study in 4T1 tumor-bearing Balb/C mice treated IP with Nico-52 at 10 mg/kg three times a week

**Figure S23.** Nico-52 UPLC-MS absorbance AUC standard curve

**Table S1.** Nico-52 R^1^ analogs tested at a 10 µM concentration

**Table S2.** Nico-52 R^2^ analogs tested at a 10 µM concentration

**Table S3.** Nico-52 R^3^ analogs tested at a 10 µM concentration

**Table S4.** Coefficient of determination (r^2^) values for the Nico-52 analogs which were further evaluated for dose response in HEK Blue IL-4/IL-13 reporter cell line.

**Table S5**. Control data for HEK-Blue IL-4/IL-13 inhibition experiments for analog 14 and 15

**Table S6**. Control data for HEK-Blue IL-4/IL-13 inhibition experiments for lead analog 53 HPLC and LC -MS traces of representative analogs

## ACKNOWLEDGMENT

The authors would like to acknowledge the Koehler lab at MIT for use of their Biorad CFX Real-Time PCR System for DSF studies. This publication was supported by the National Center for Advancing Translational Sciences, National Institutes of

Health, through Boston University Clinical & Translational Science Institute Grant Number 1UL1TR001430. Its contents are solely the responsibility of the authors and do not necessarily represent the official views of the National Institutes of Health.

## ABBREVIATIONS

IL-4: Interleukin-4
IL-13: Interleukin-13
TNF-α: Tumor Necrosis Factor-α
IL-1β: Interleukin-1 β
IL-2: Interleukin-2
IL-33: Interleukin-33
IL-33: Interleukin-33
IL-15: Interleukin-15
IL-23: Interleukin-23
IL-17: Interleukin-17
PD-L1: Programmed Cell Death Ligand 1
NSCLC: Non-small cell lung cancer
TLC: Thin layer chromatography
SEAP: Secreted Embryonic Alkaline Phosphatase.

