## Supplemental for "Potent and Selective IL-4 Inhibitors with Anti-Tumor Activity"

**Supplemental Information**

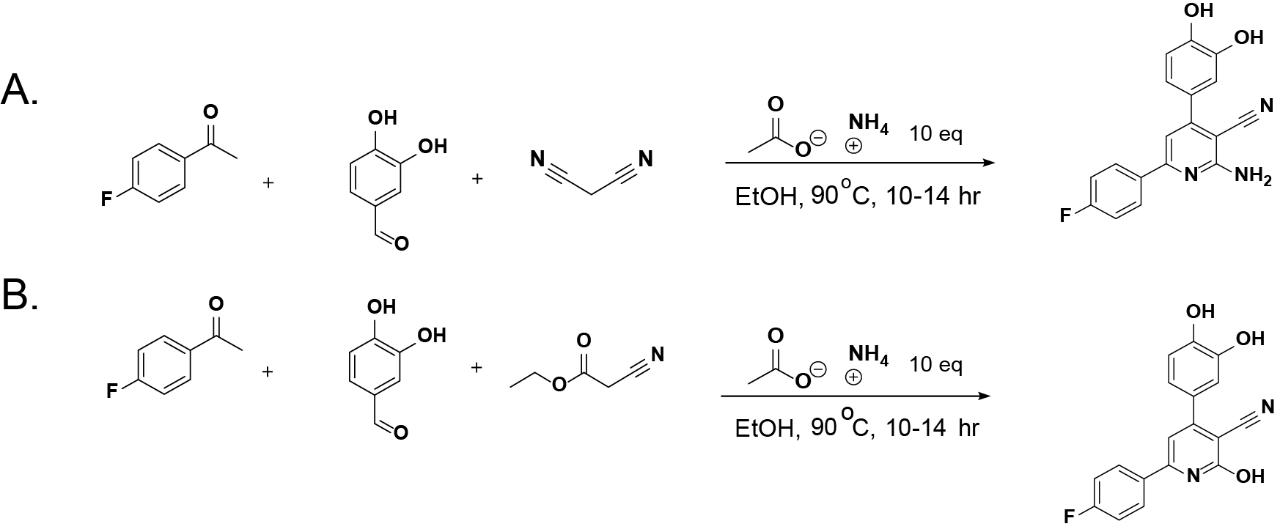
**Supplemental Figure 1. Nico-52 Analog’s Synthesis Scheme for Investigating Structure-Activity Relationships. (A)** Synthesis of **Nico-**52 and amino nicotinonitrile containing R^1^ and R^2^ analogs**. (B)** Synthesis of hydroxy nicotinonitrile **Nico-52** analogs.

**Supplemental Table 1. Nico-52 R^1^ analogs tested at a 10 µM concentration.** For each analog, the structural changes made at R^1^ position are indicated. Blank spaces indicate that the structural position matches that of the parent compound **Nico-52**.

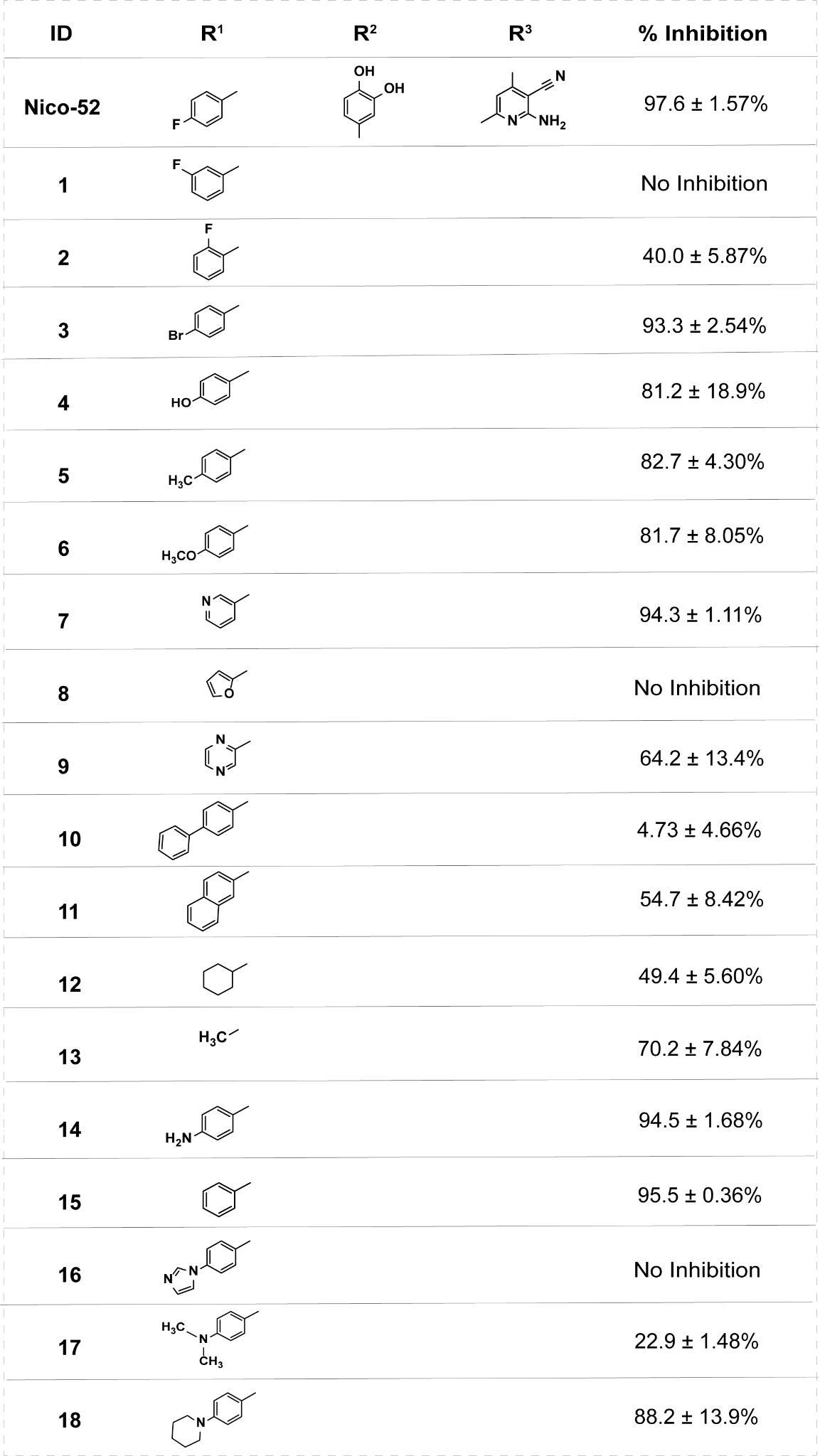

**Supplemental Table 2. Nico-52 R^2^ analogs tested at a 10 µM concentration.** For each analog, the structural changes made at R^2^ position are indicated. Blank spaces indicate that the structural position matches that of the parent compound **Nico-52**.

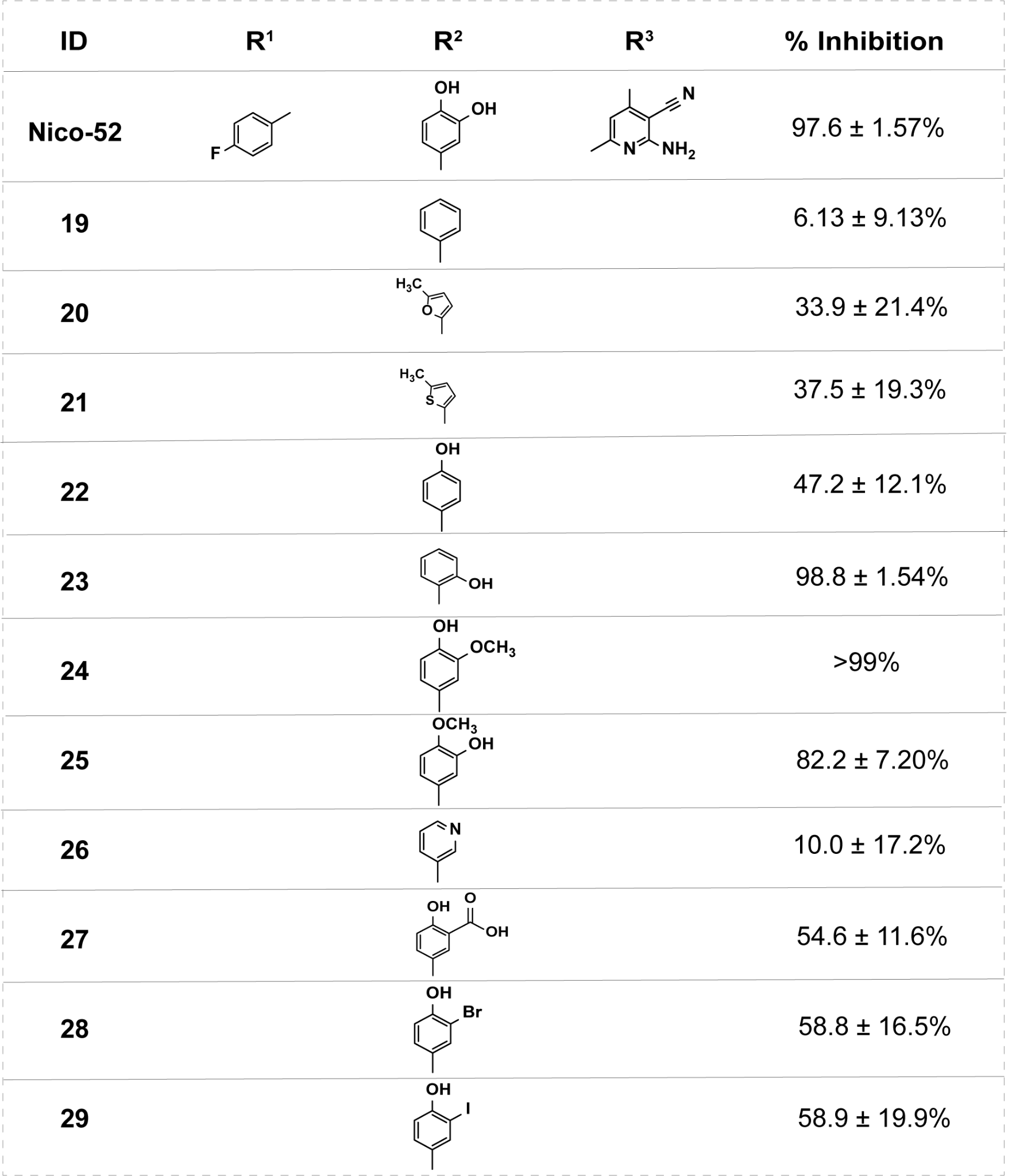

**Supplemental Table 3. Nico-52 R^3^ analogs tested at a 10 µM concentration.** For each analog, the structural changes made at R^3^ position are indicated. Blank spaces indicate that the structural position matches that of the parent compound **Nico-52**.

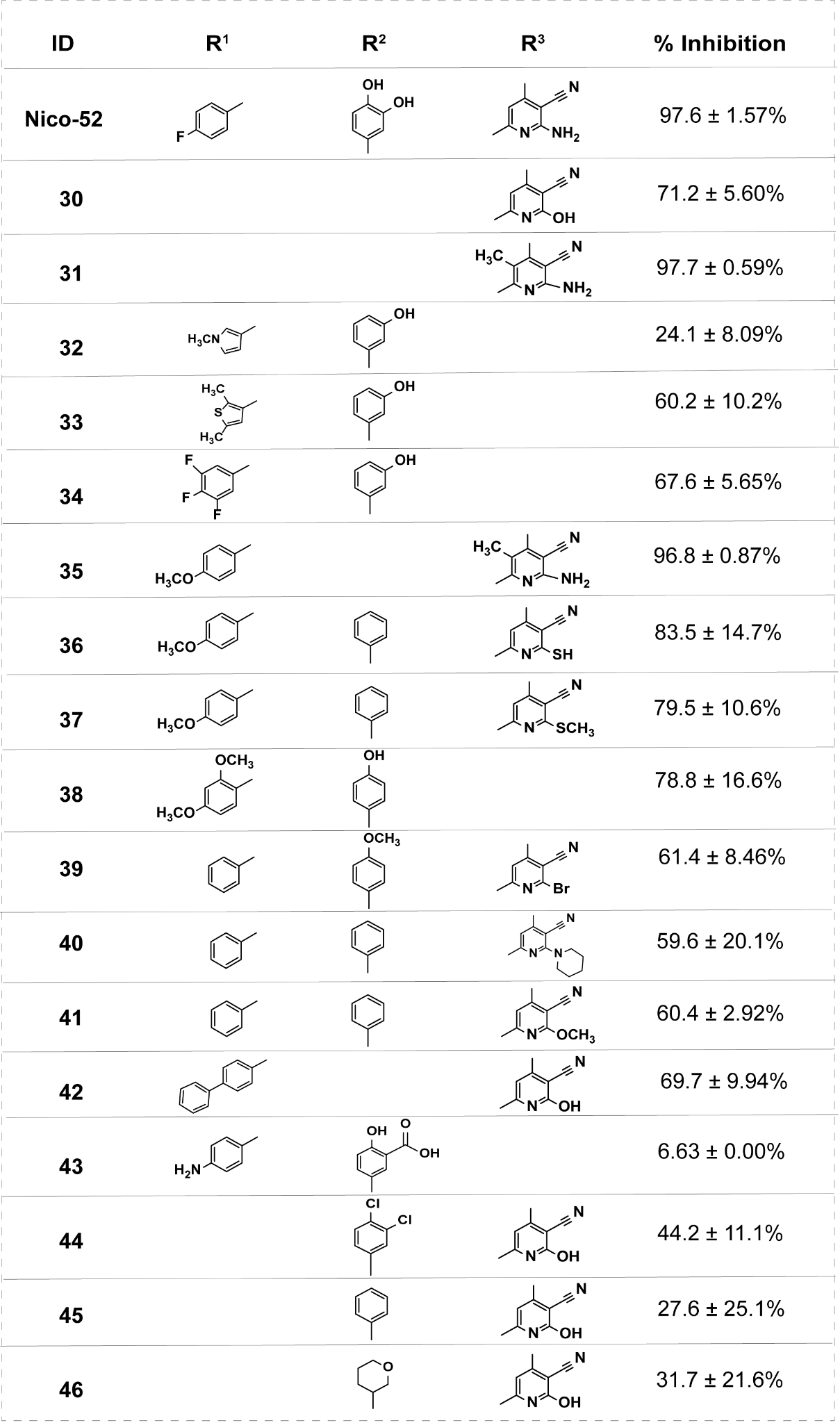

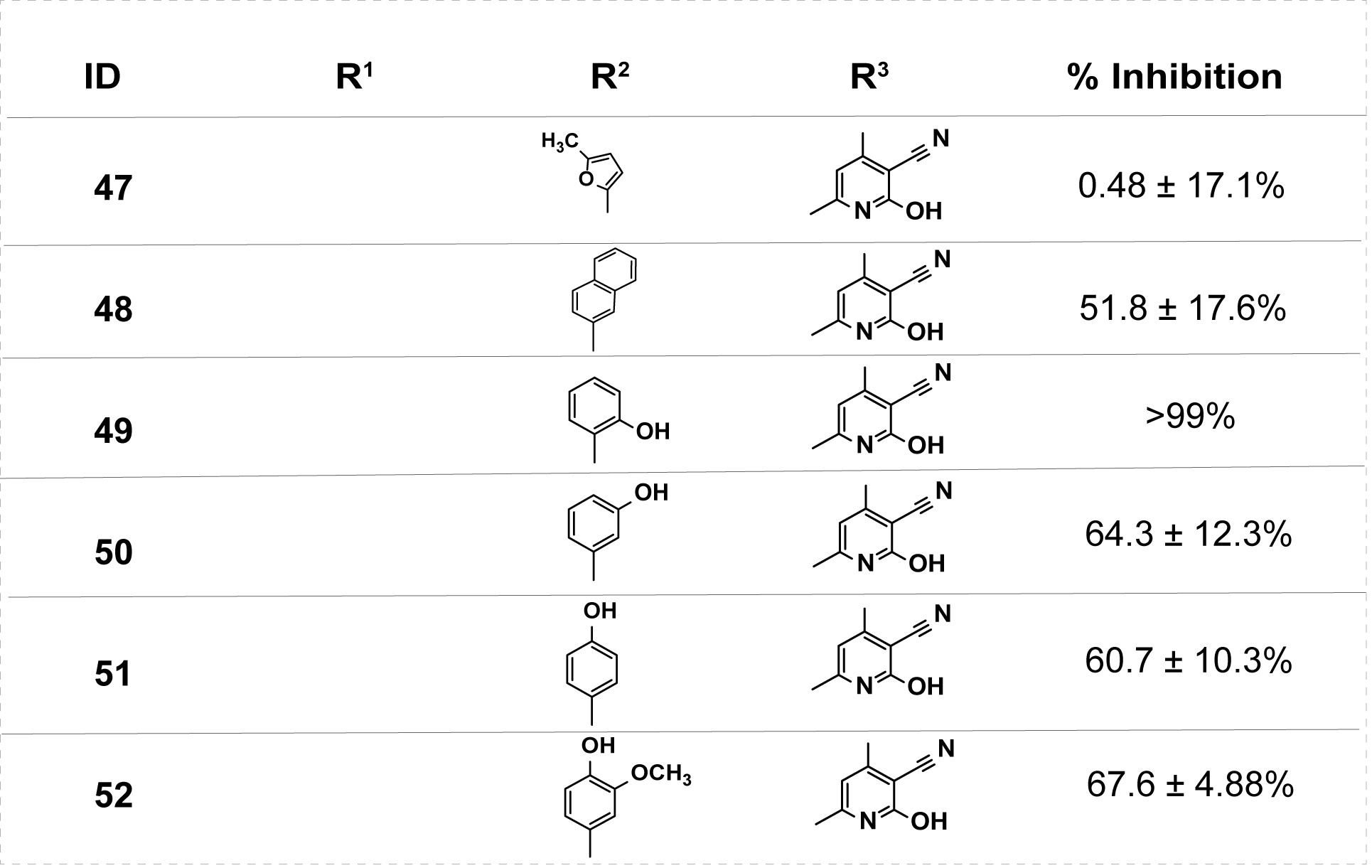

**Supplemental Table 4. Coefficient of determination (r^2^) values for the Nico-52 analogs which were further evaluated for dose response in HEK Blue IL-4/IL-13 reporter cell line.**

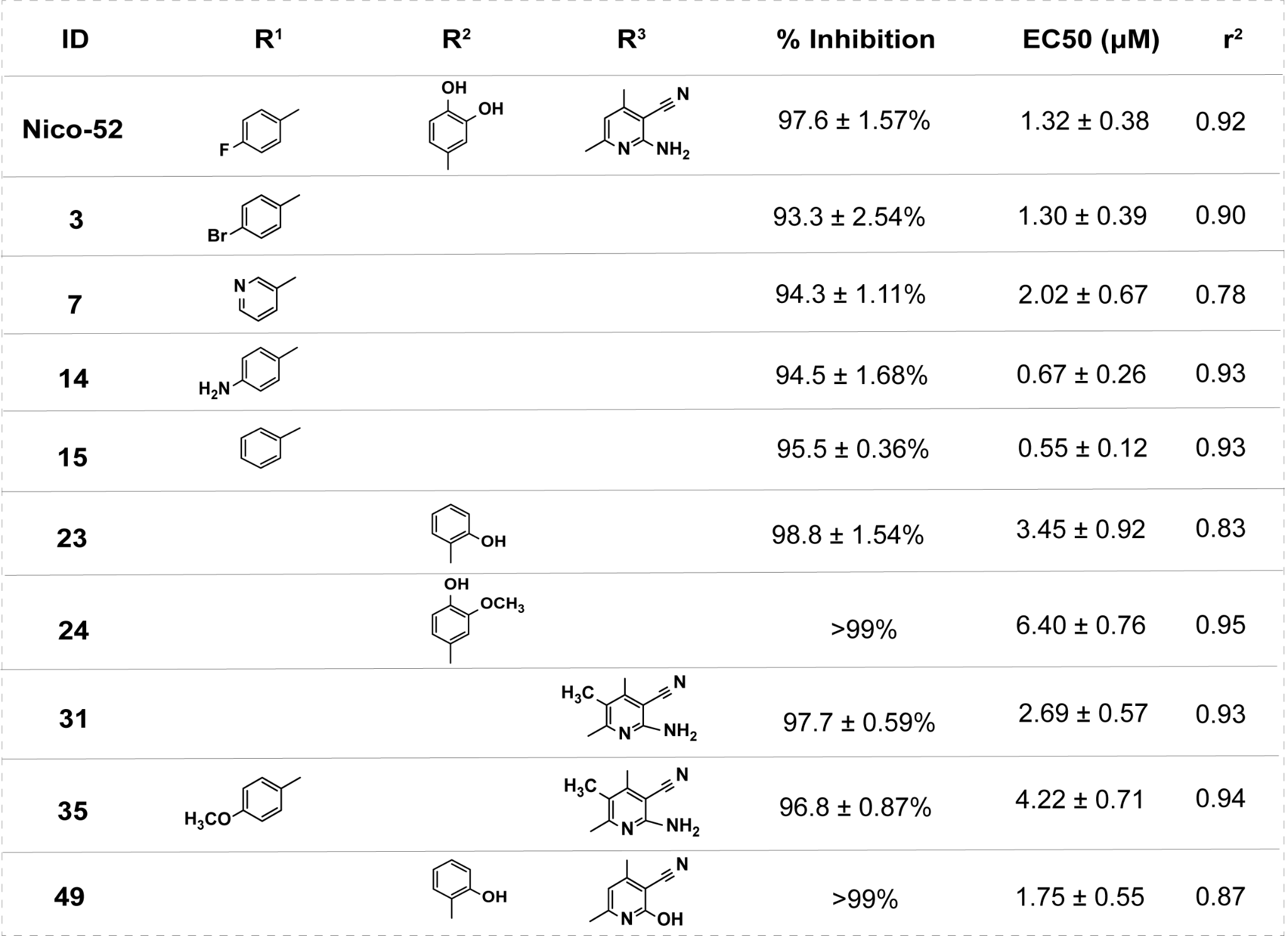

**Supplemental Table 5. Control data for HEK-Blue IL-4/IL-13 inhibition experiments for analog 14 and 15.**

| **Condition** | **Abs650** |
| --- | --- |
| Cells alone | 0.05825 ± 0.00035 |
| Cells with vehicle | 0.05753 ± 0.000791 |
| Cells with IL-4 and vehicle | 0.23874 ± 0.037224 |
| Cells with 10 µM Compound 14 only | 0.05948 ± 0.000642 |
| Cells with 10 µM Compound 15 only | 0.06345 ± 0.001036 |

**Supplemental Figure 2. Analog 15 selectivity evaluation in HEK Blue IL-2 reporter cell line.**

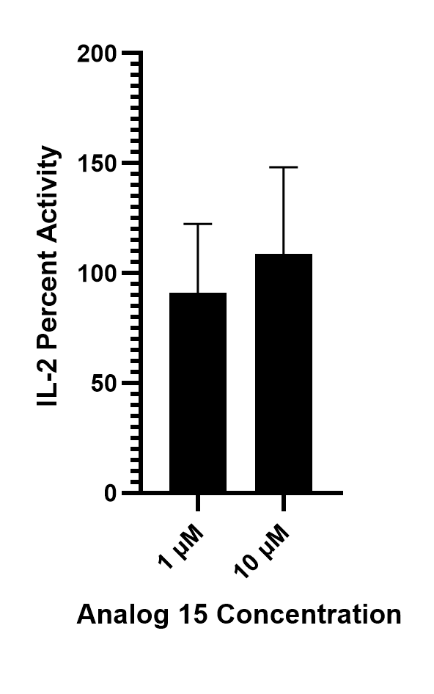
Analog 15 was tested at 1 µM and 10 µM in HEK-Blue IL-2 reporter cells. Treatment did not alter IL-2–induced SEAP production, confirming that Analog 15 selectively targets IL-4.

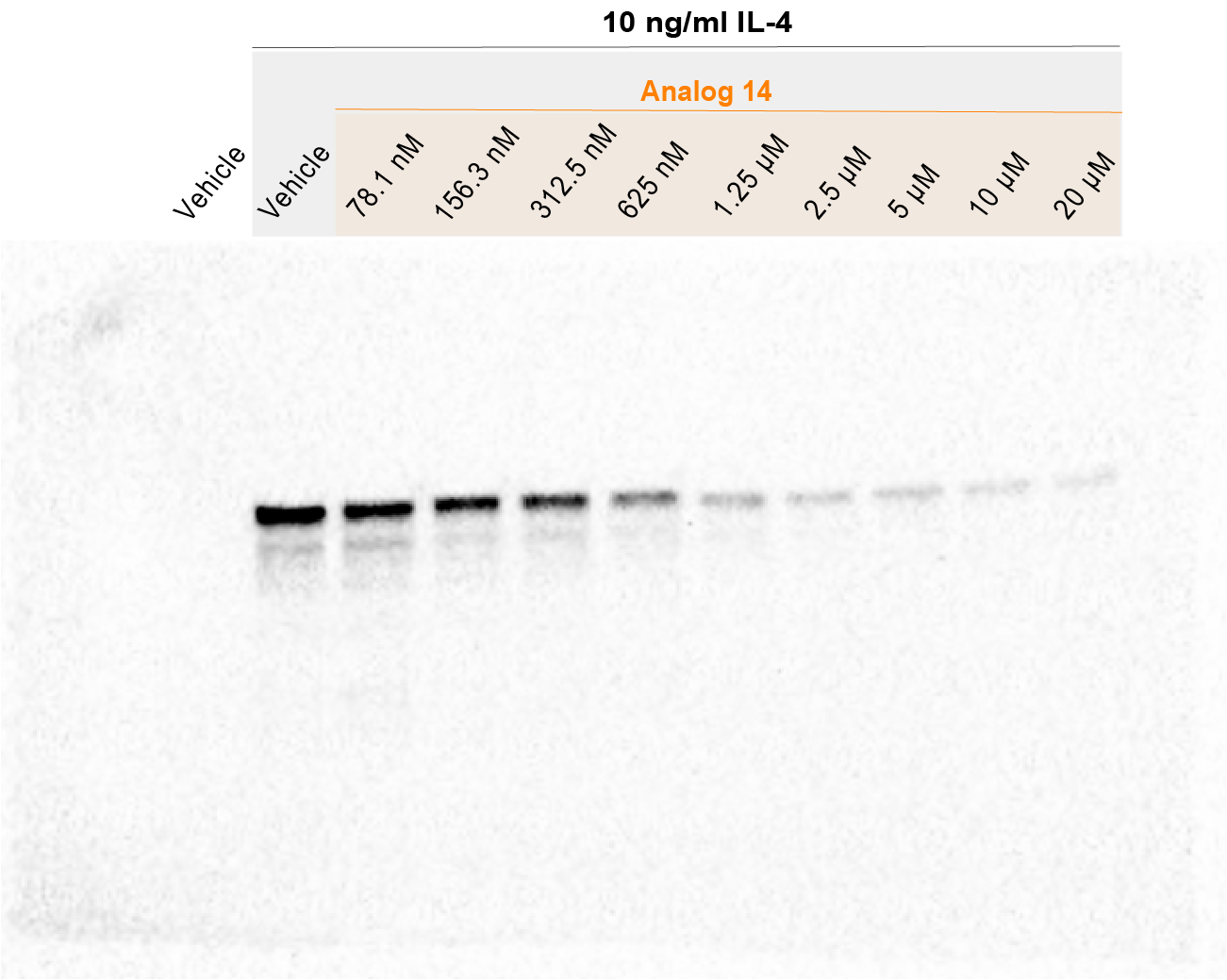

**Supplemental Figure 3. pSTAT-6 Western blot for THP-1 cells for analog 14**

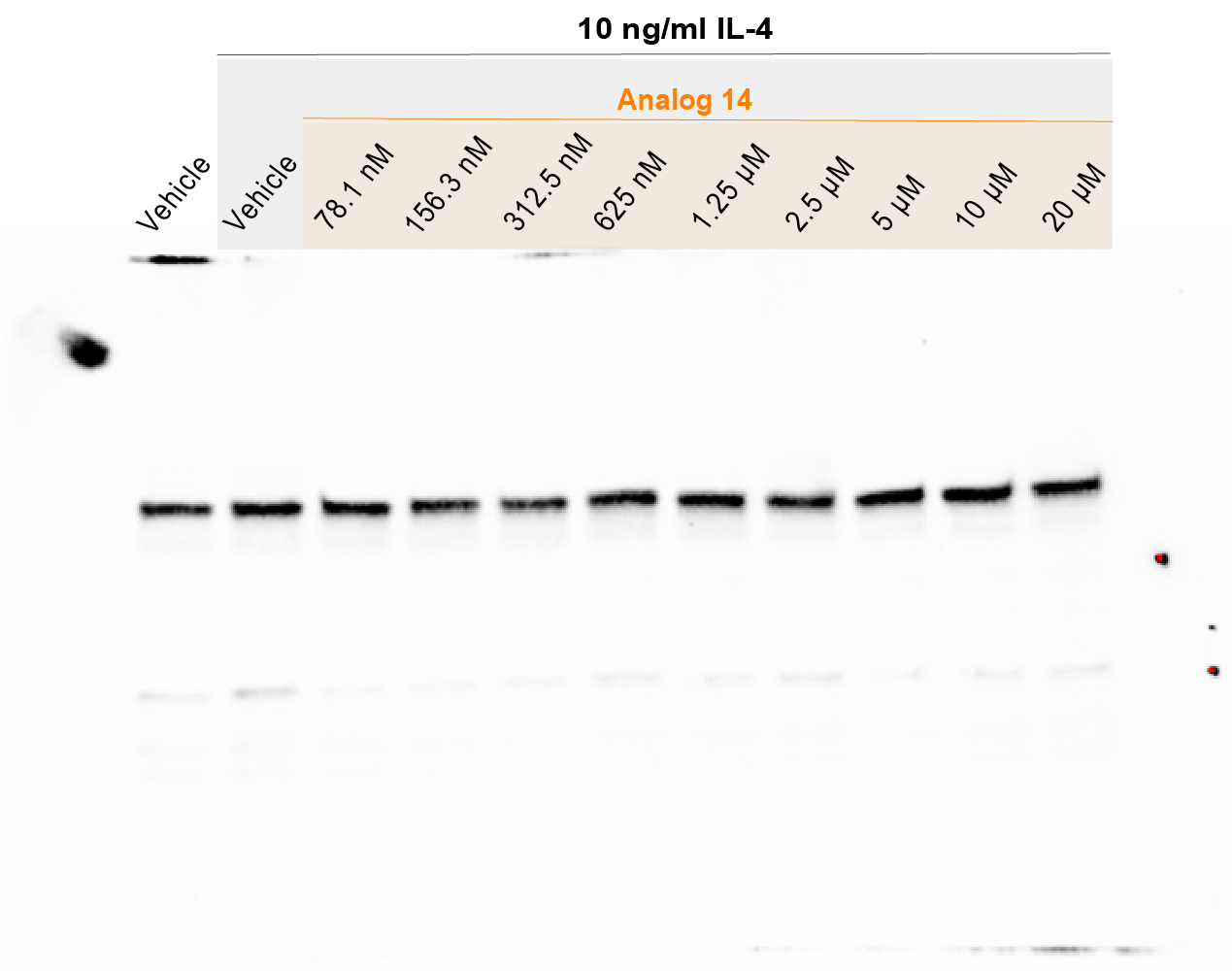

**Supplemental Figure 4. STAT-6 Western blot for THP-1 cells for analog 14**

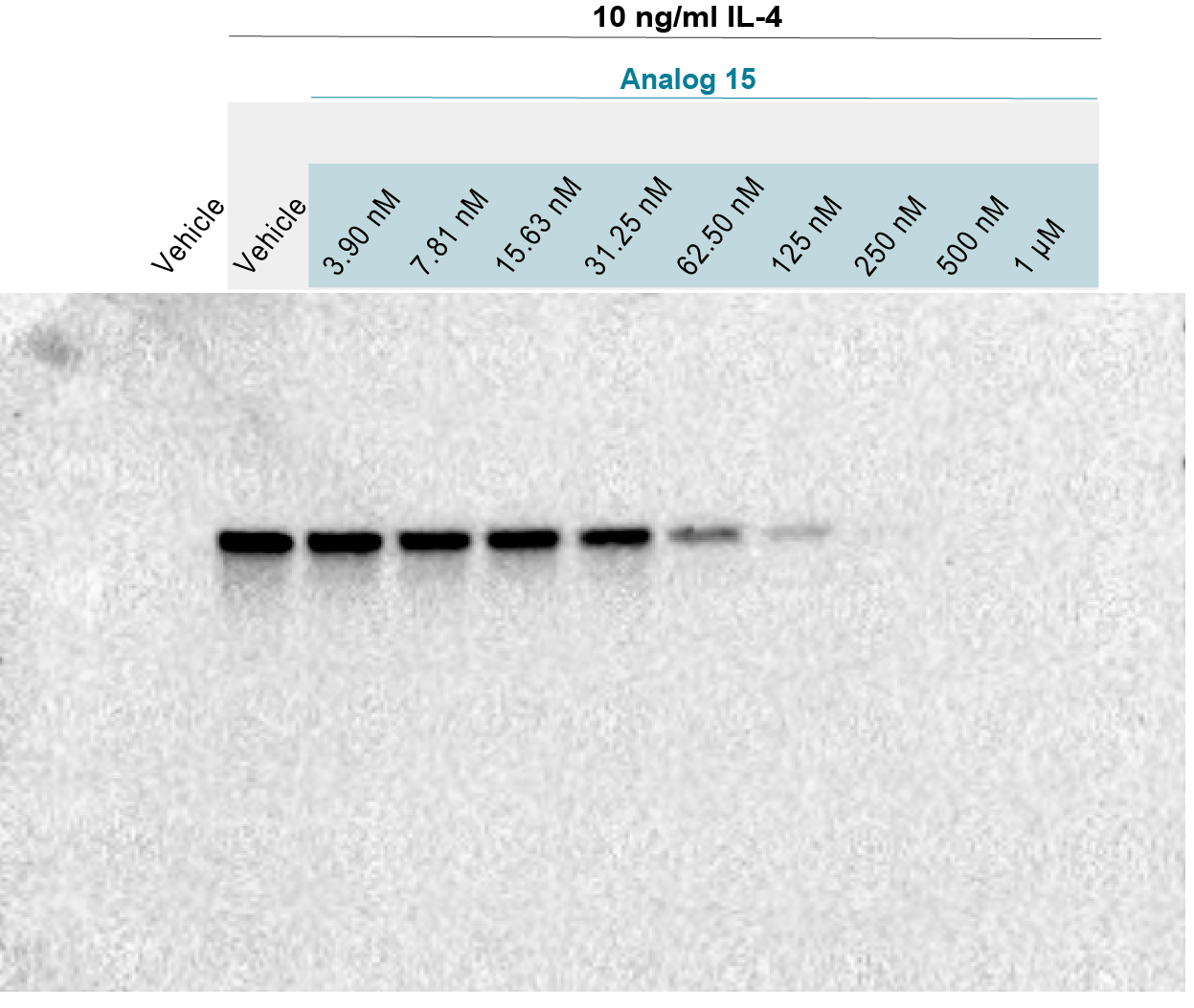

**Supplemental Figure 5. pSTAT-6 Western blot for THP-1 cells for analog 15**

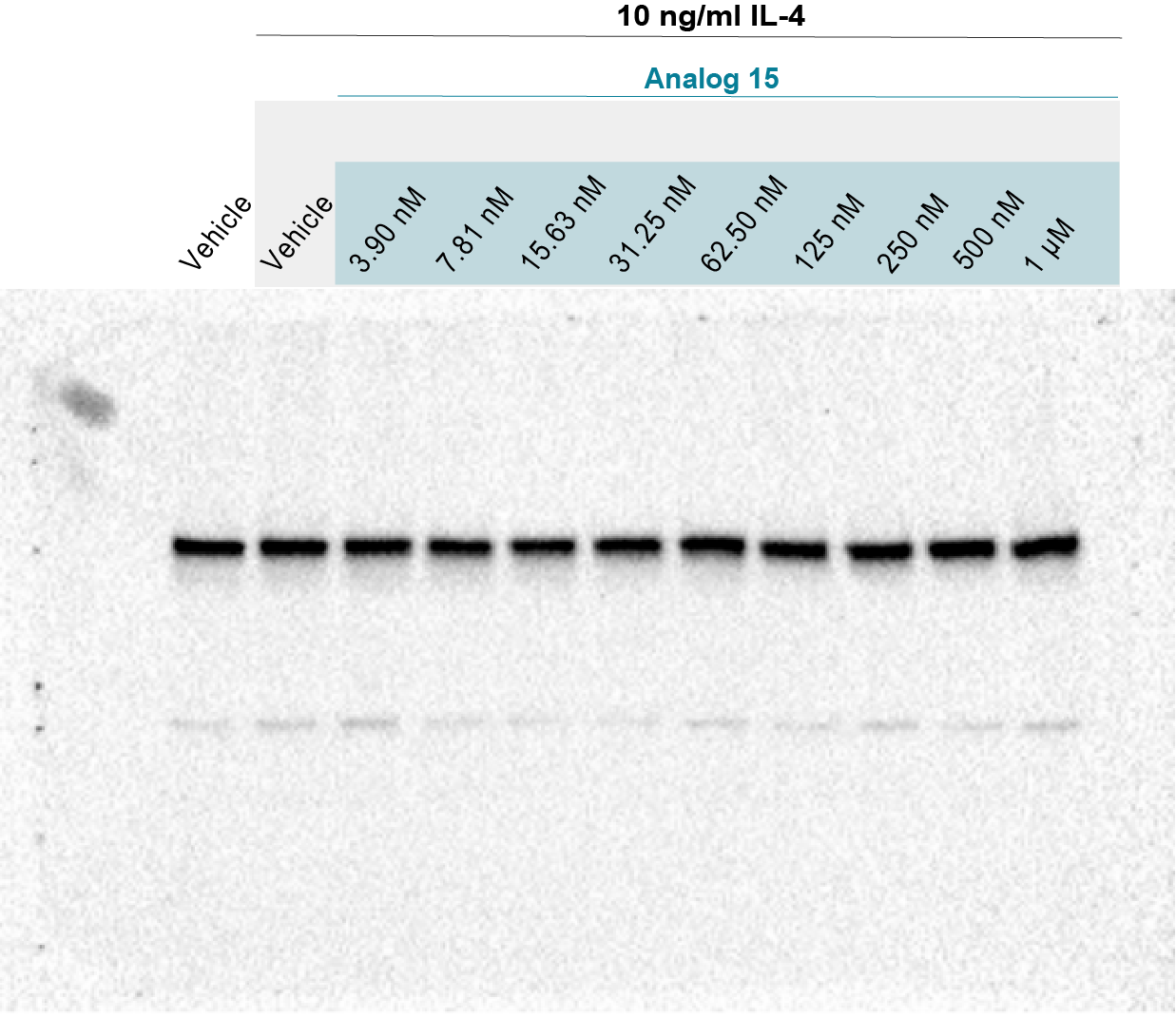

**Supplemental Figure 6. STAT-6 Western blot for THP-1 cells for analog 15**

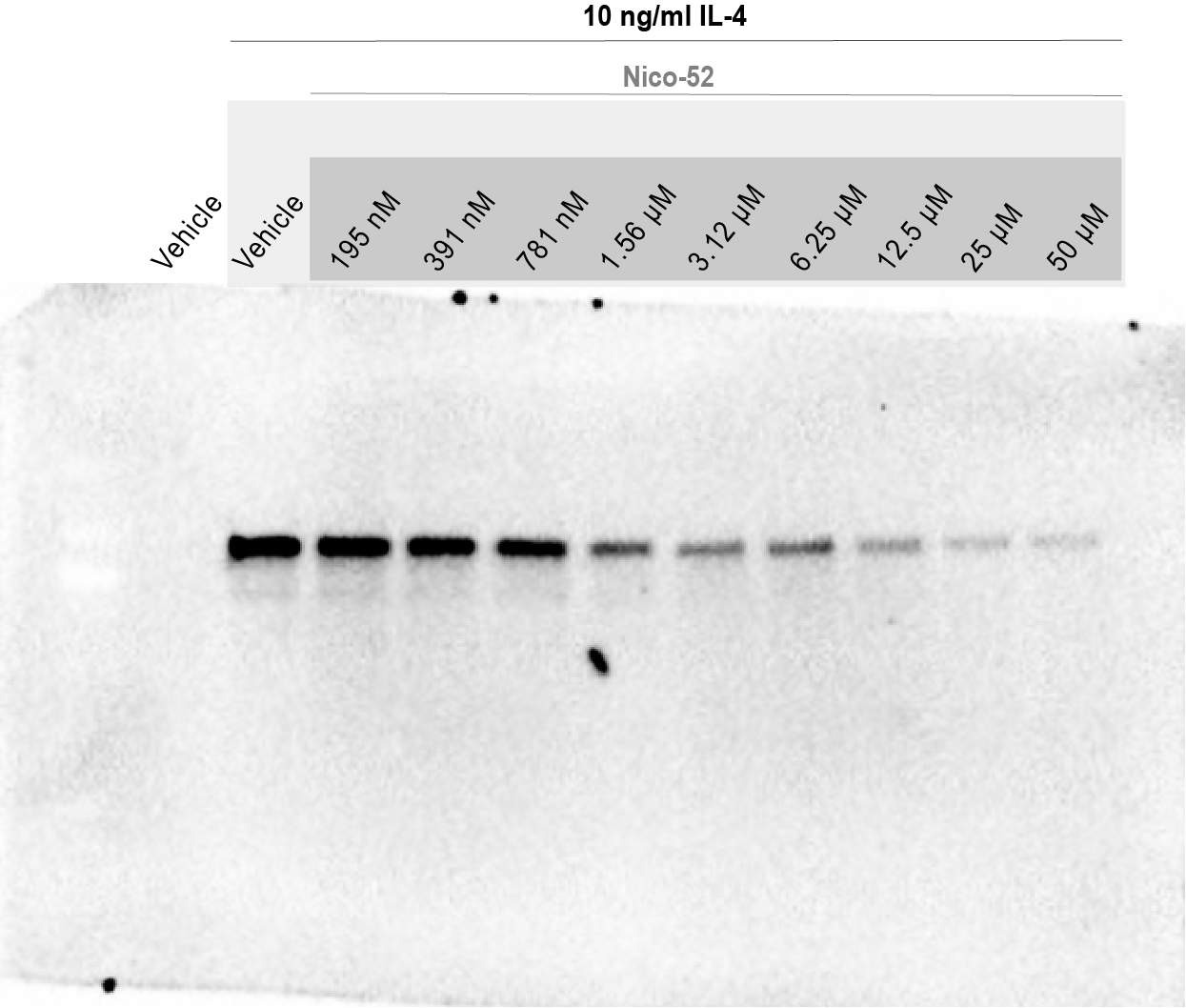

**Supplemental Figure 7. pSTAT-6 Western blot for Ramos cells for Nico-52**

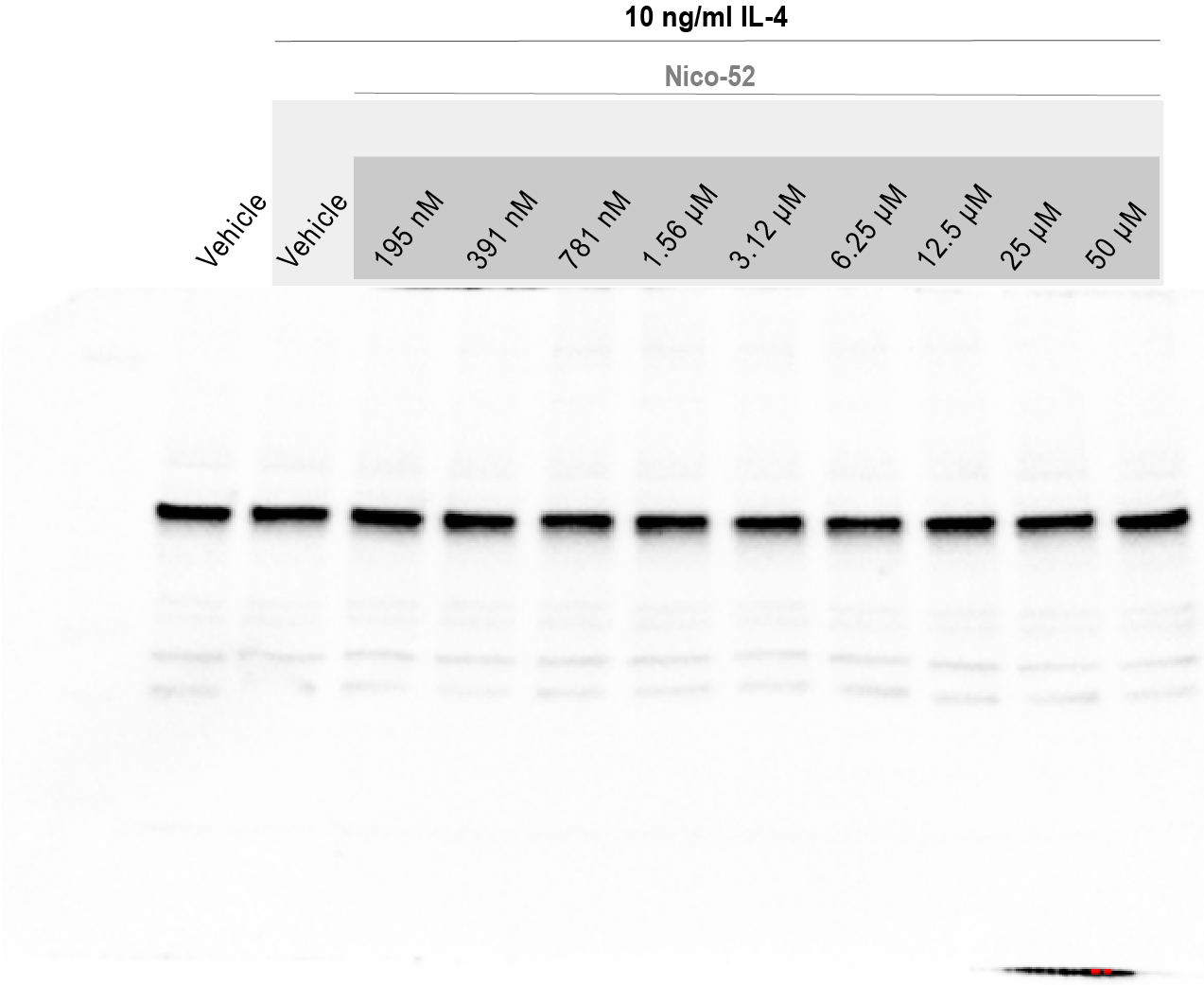

**Supplemental Figure 8. STAT-6 Western blot for Ramos cells for Nico-52**

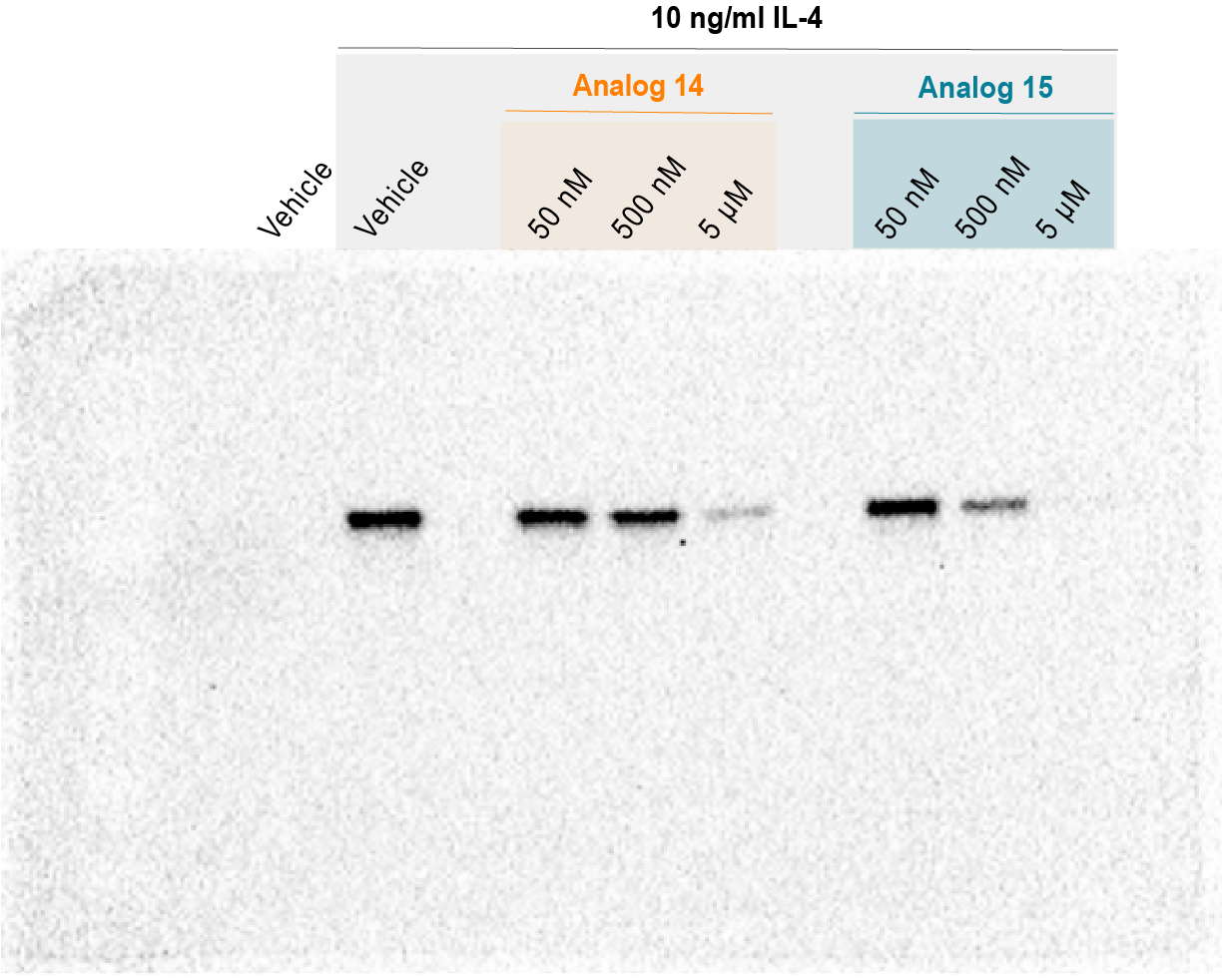

**Supplemental Figure 9. pSTAT-6 Western blot for Ramos cells for analogs 14 and 15**

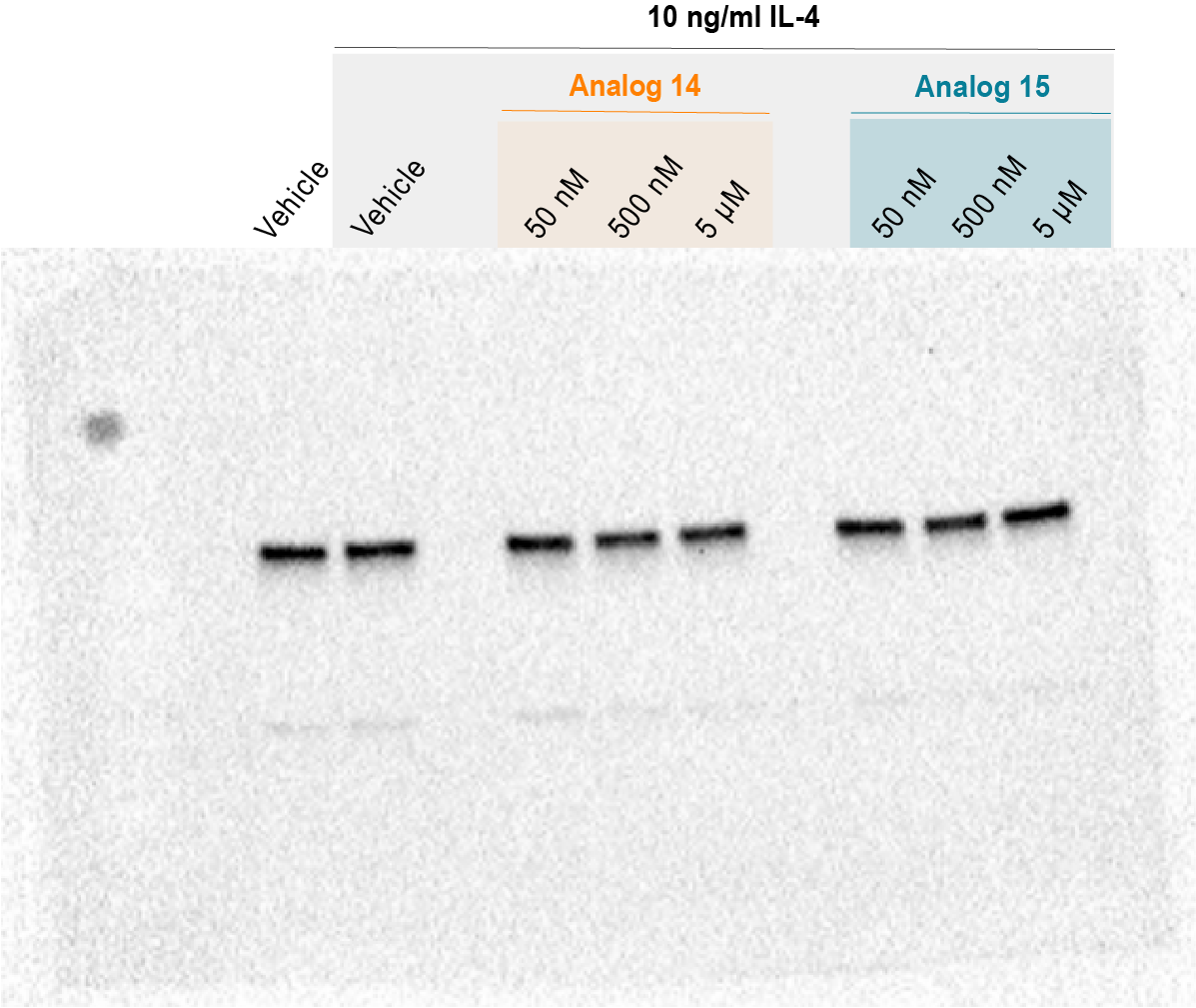

**Supplemental Figure 10. STAT-6 Western blot for Ramos cells for analogs 14 and 15**

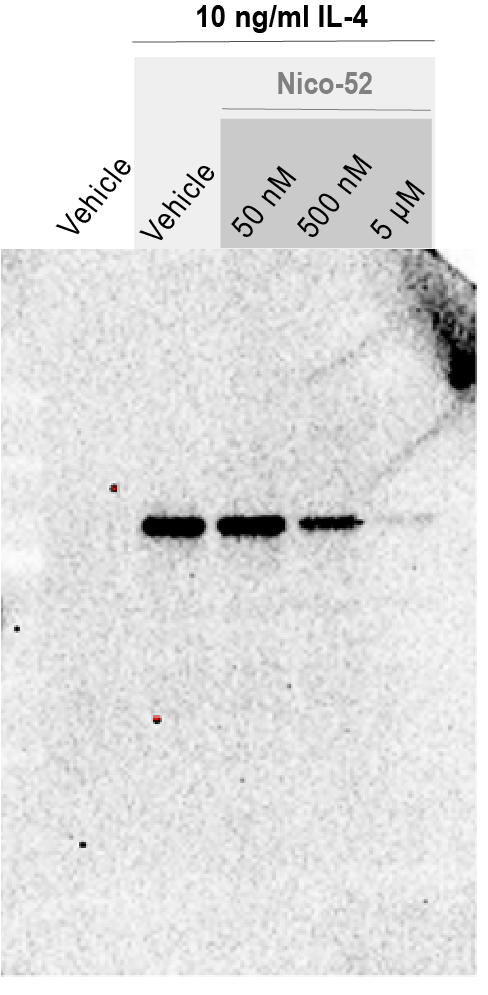

**Supplemental Figure 11. pSTAT-6 Western blot for Ramos cells for Nico-52**

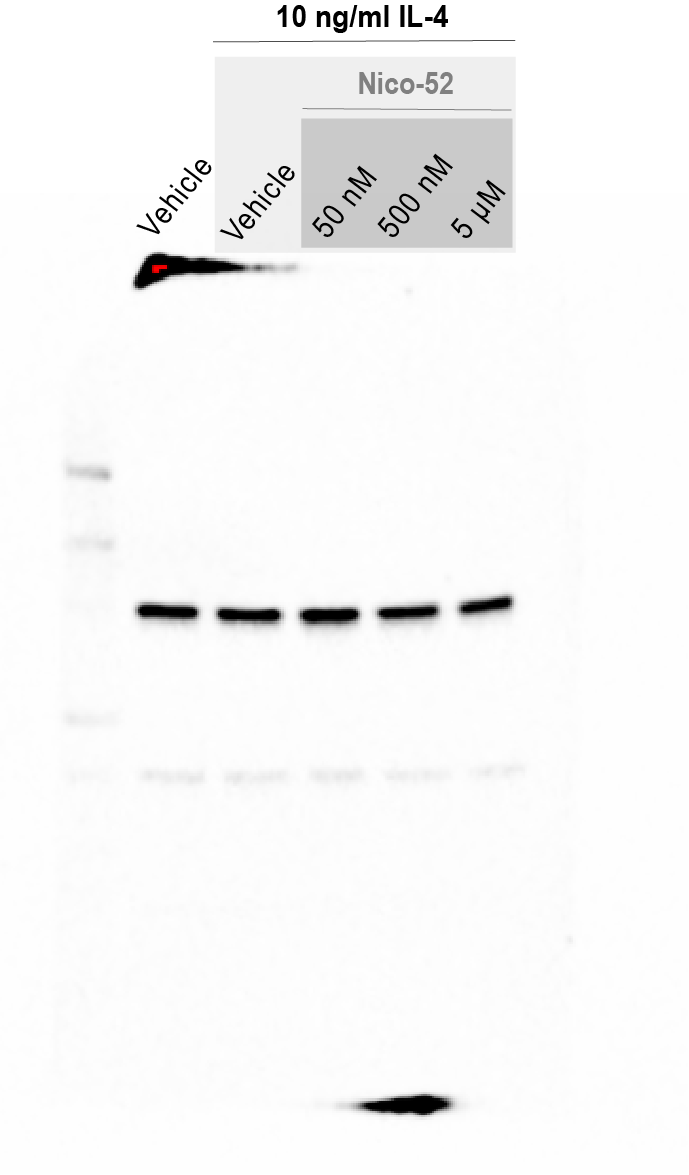

**Supplemental Figure 12. STAT-6 Western blot for Ramos cells for Nico-52**

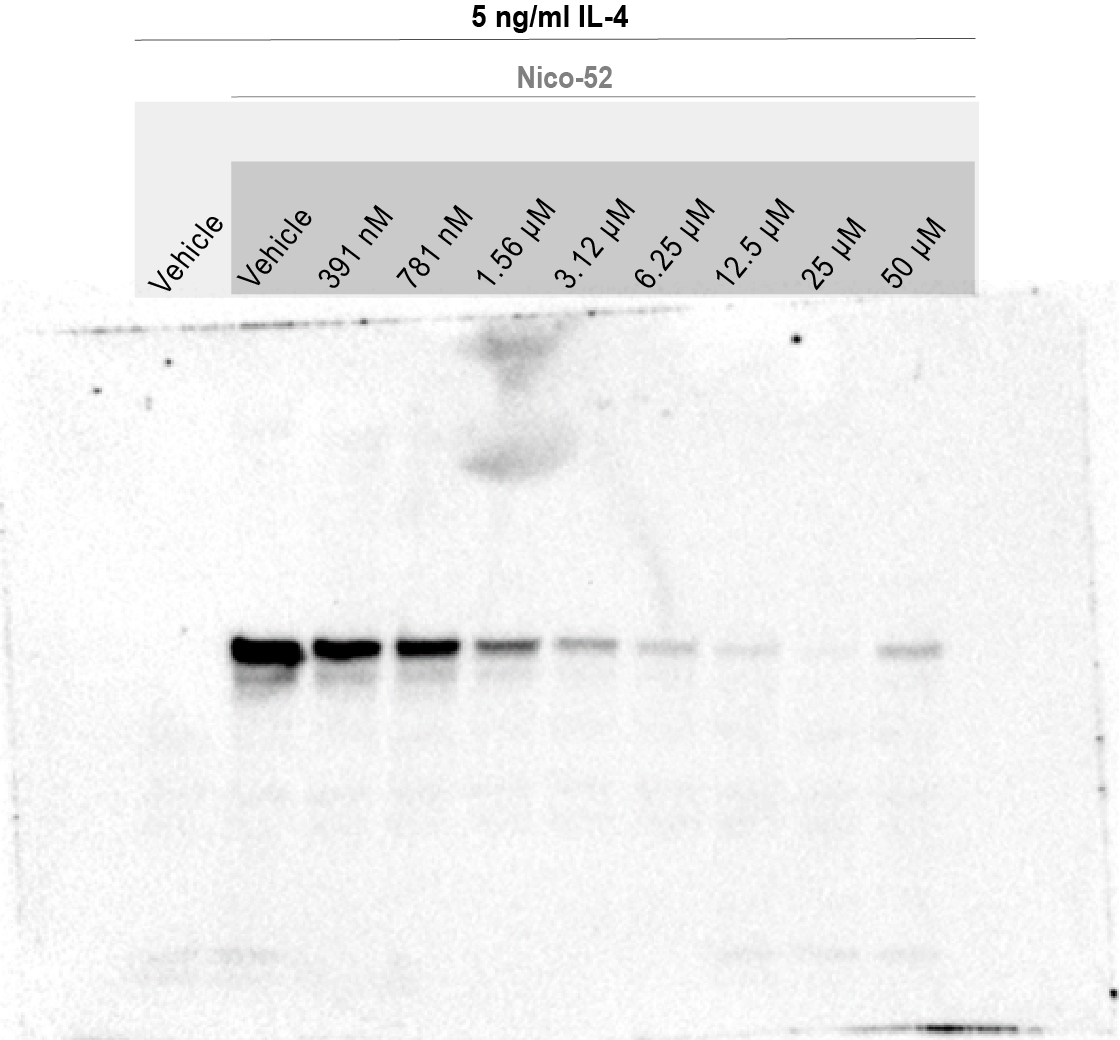

**Supplemental Figure 13. pSTAT-6 Western blot for Raw macrophage cells for Nico-52**

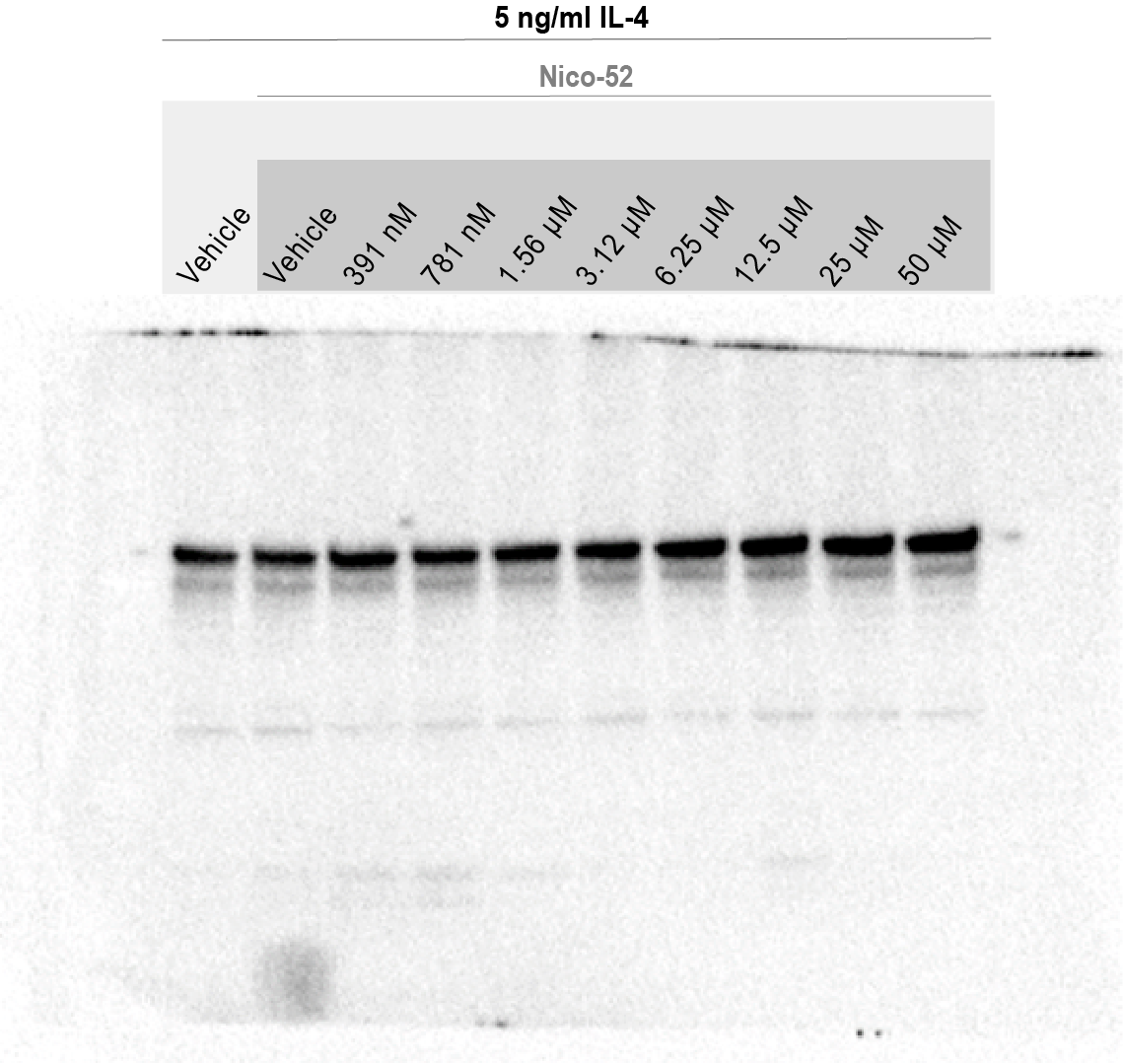

**Supplemental Figure 14. STAT-6 Western blot for Raw macrophage cells for Nico-52**

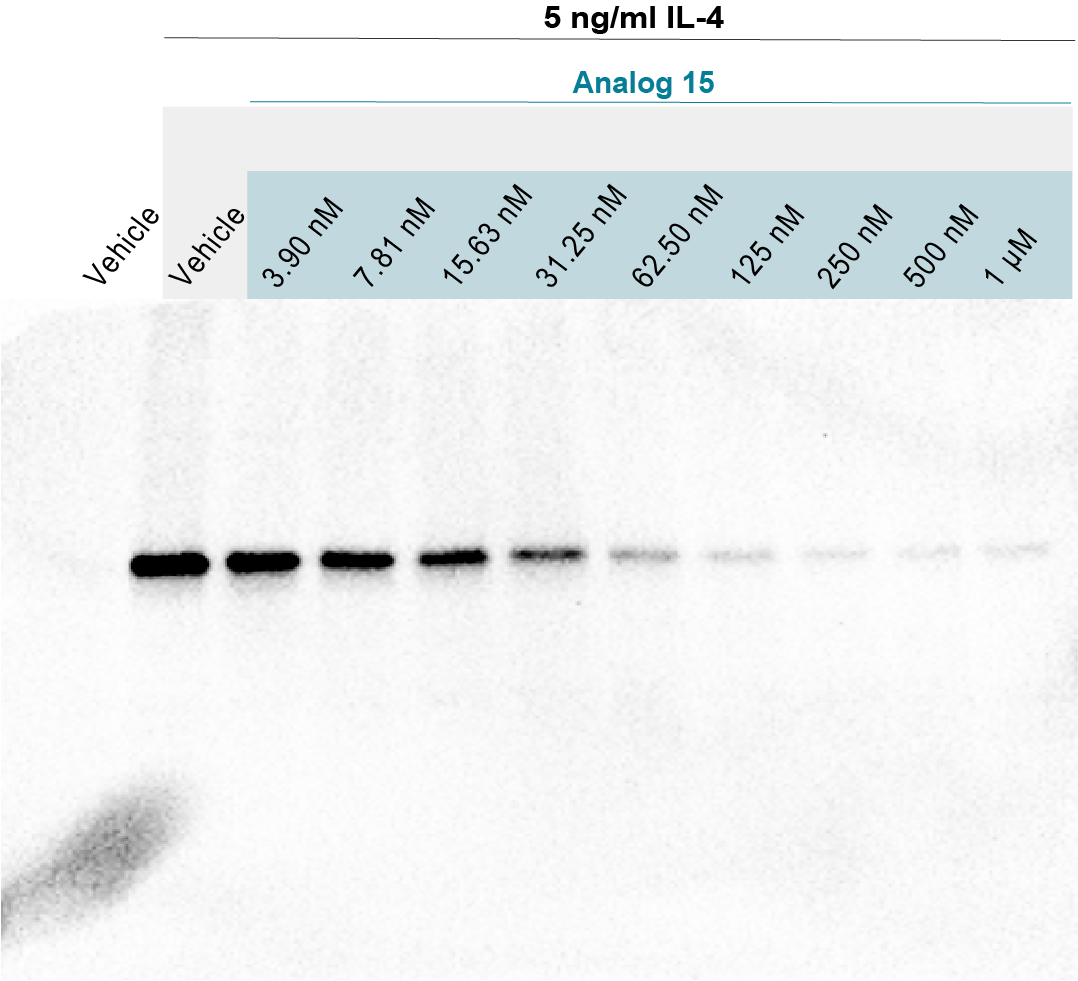

**Supplemental Figure 15. pSTAT-6 Western blot for Raw macrophage cells for**
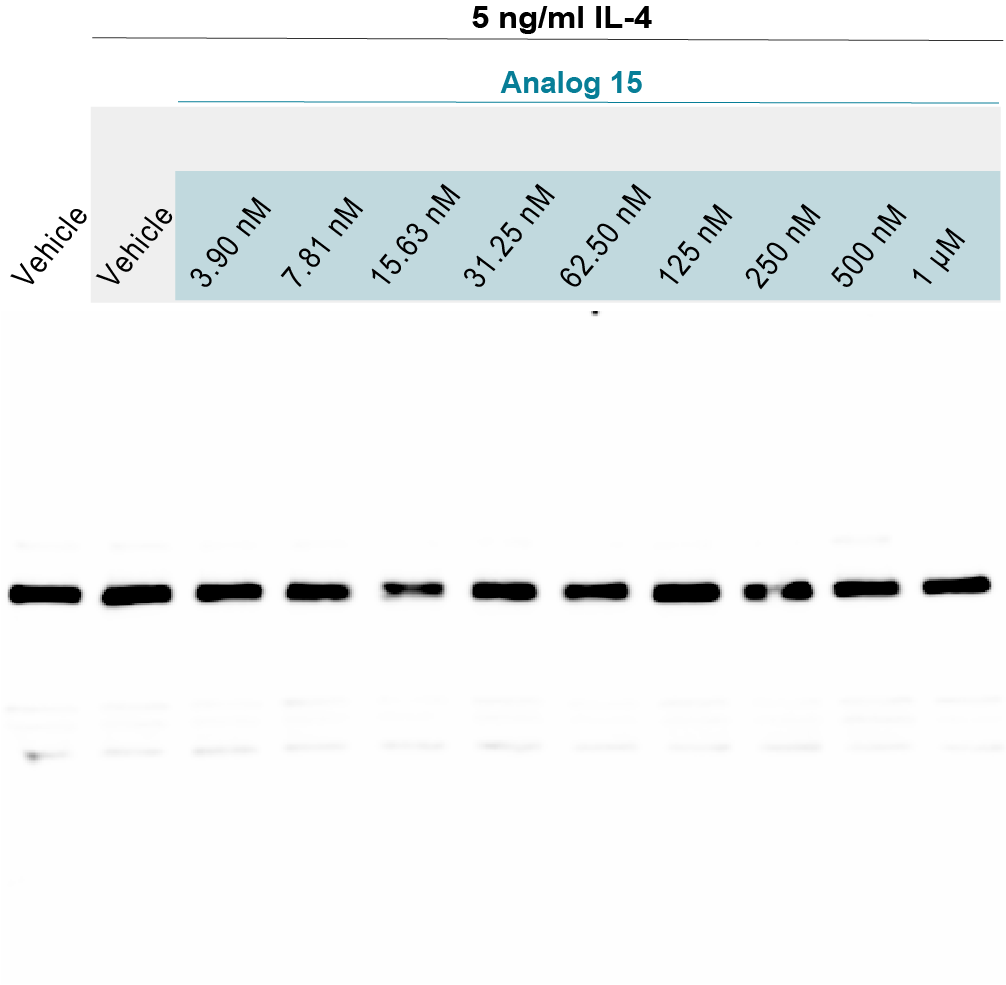
**Analog 15**

**Supplemental Figure 16. STAT-6 Western blot for Raw macrophage cells for Analog 15**

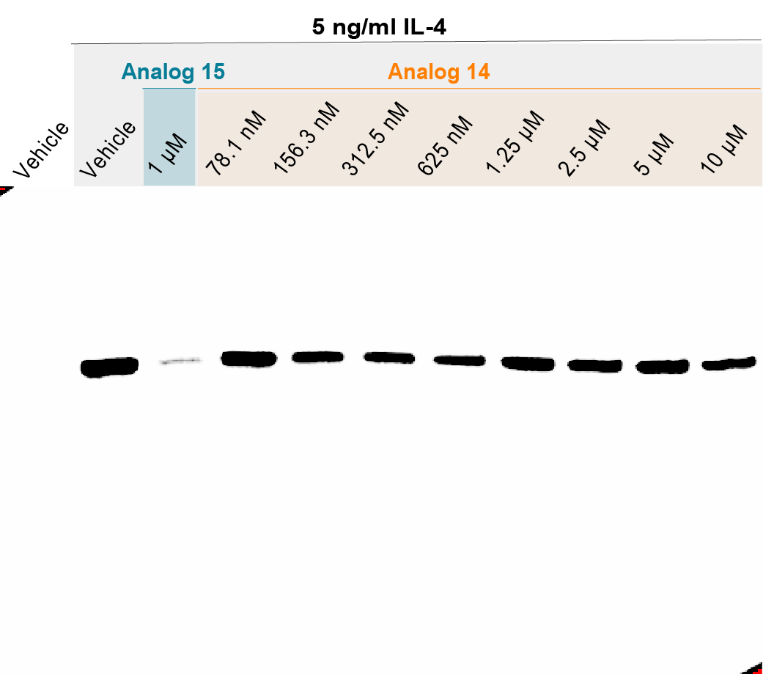
**Supplemental Figure 17. pSTAT-6 Western blot for Raw macrophage cells for Analog 14**

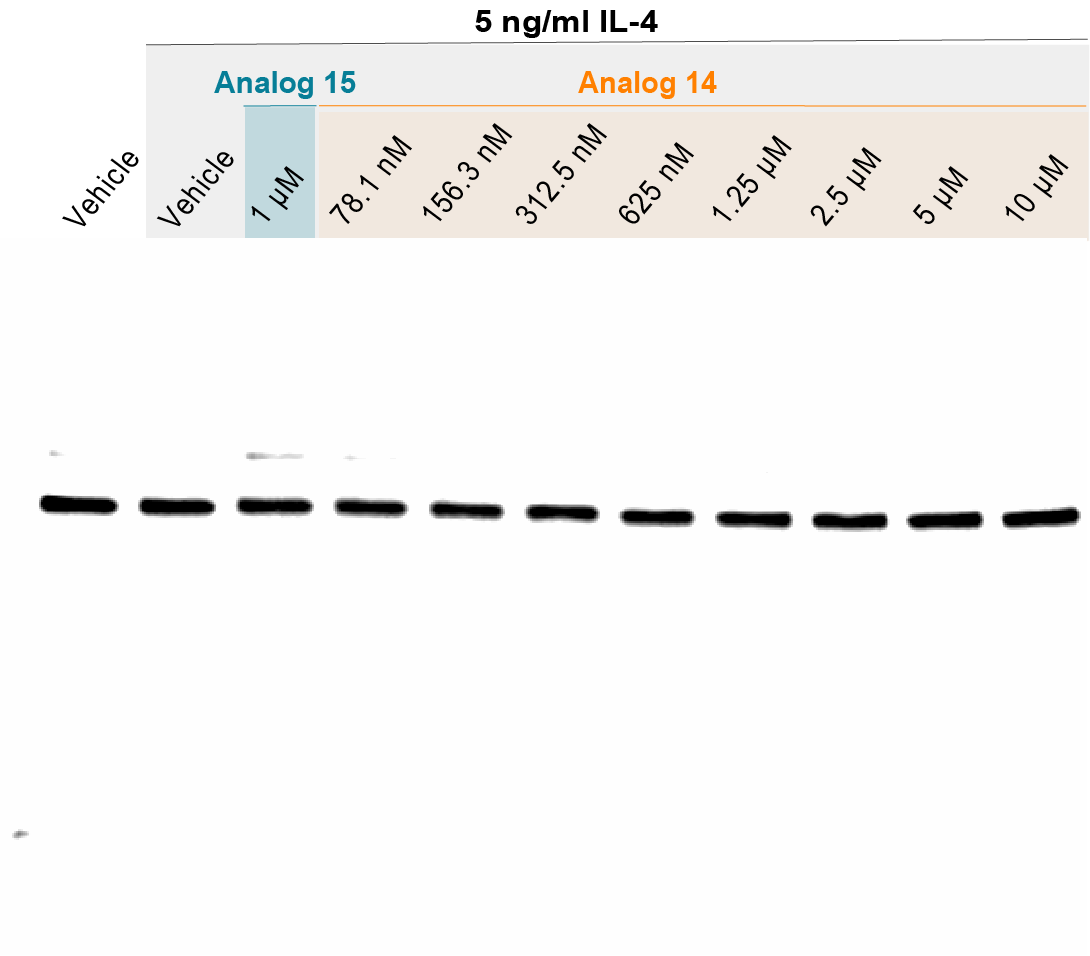

**Supplemental Figure 18. STAT-6 Western blot for Raw macrophage cells for Analog 14**

* A single dose of analog 15 was tested in the same batch of cells to confirm that the reduced potency of analog 14 against mouse IL-4 was not attributable to the cells.

**Supplemental Figure 19. Lead analog 53 dose response in HEK Blue IL-4/IL-13 reporter assay. (***Error bars represent mean ± SD)*

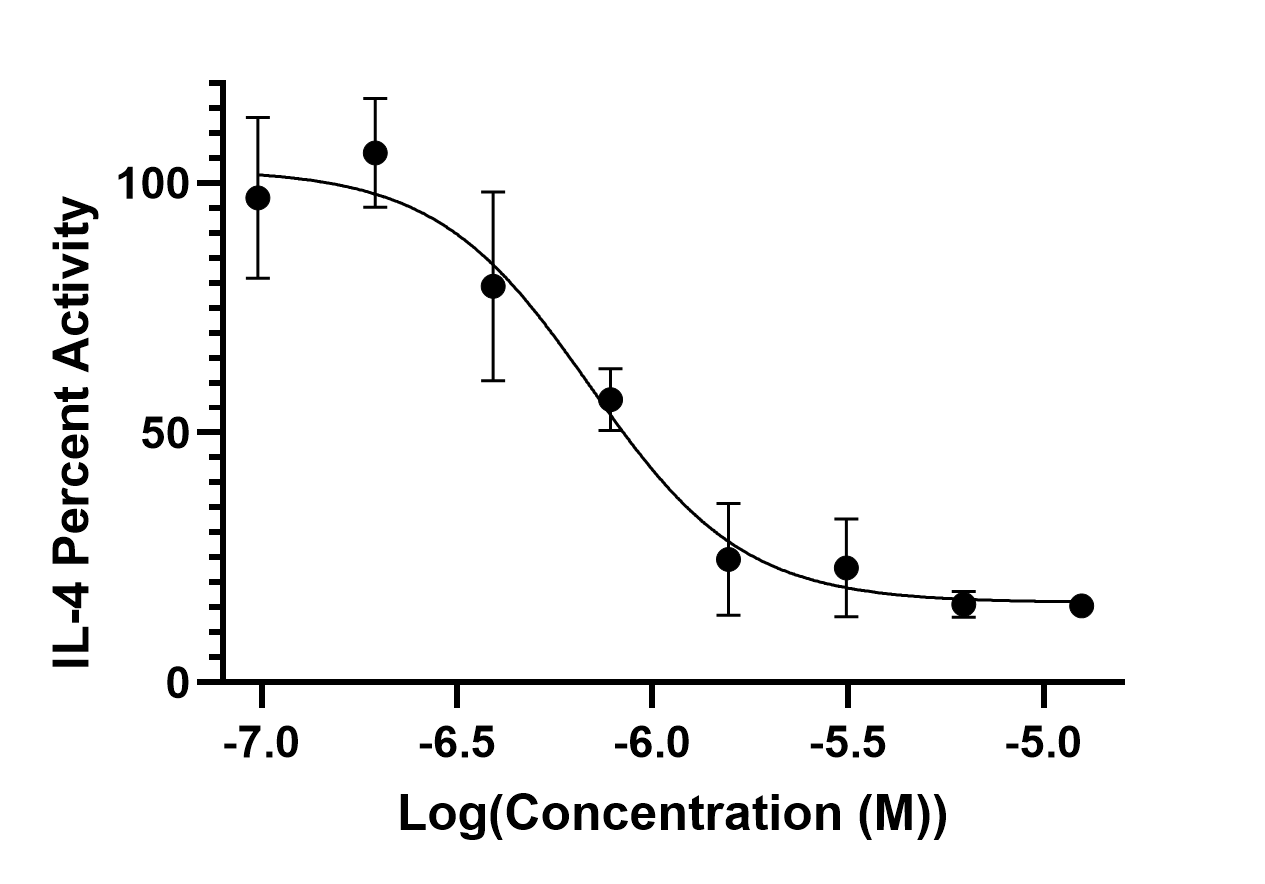

**EC_50_: 690.3 nM**

**Coefficient of determination (r^2^): 0.92**

**Supplemental Table 6. Control data for HEK-Blue IL-4/IL-13 inhibition experiments for lead analog 53.**

| **Condition** | **Abs650** |
| --- | --- |
| Cells alone | 0.06949 ± 0.005601 |
| Cells with vehicle | 0.064758 ± 0.013504 |
| Cells with IL-4 and vehicle | 0.175182 ± 0.037362 |
| Cells with 12.5 µM Compound 53 only | 0.06445 ± 0.002796 |

**Supplemental Figure 20. Body weight of C57BL/6 mice treated with Nico-52 10 mg/kg intravenously for endpoint study.** (*Error bars represent mean ± SEM).*

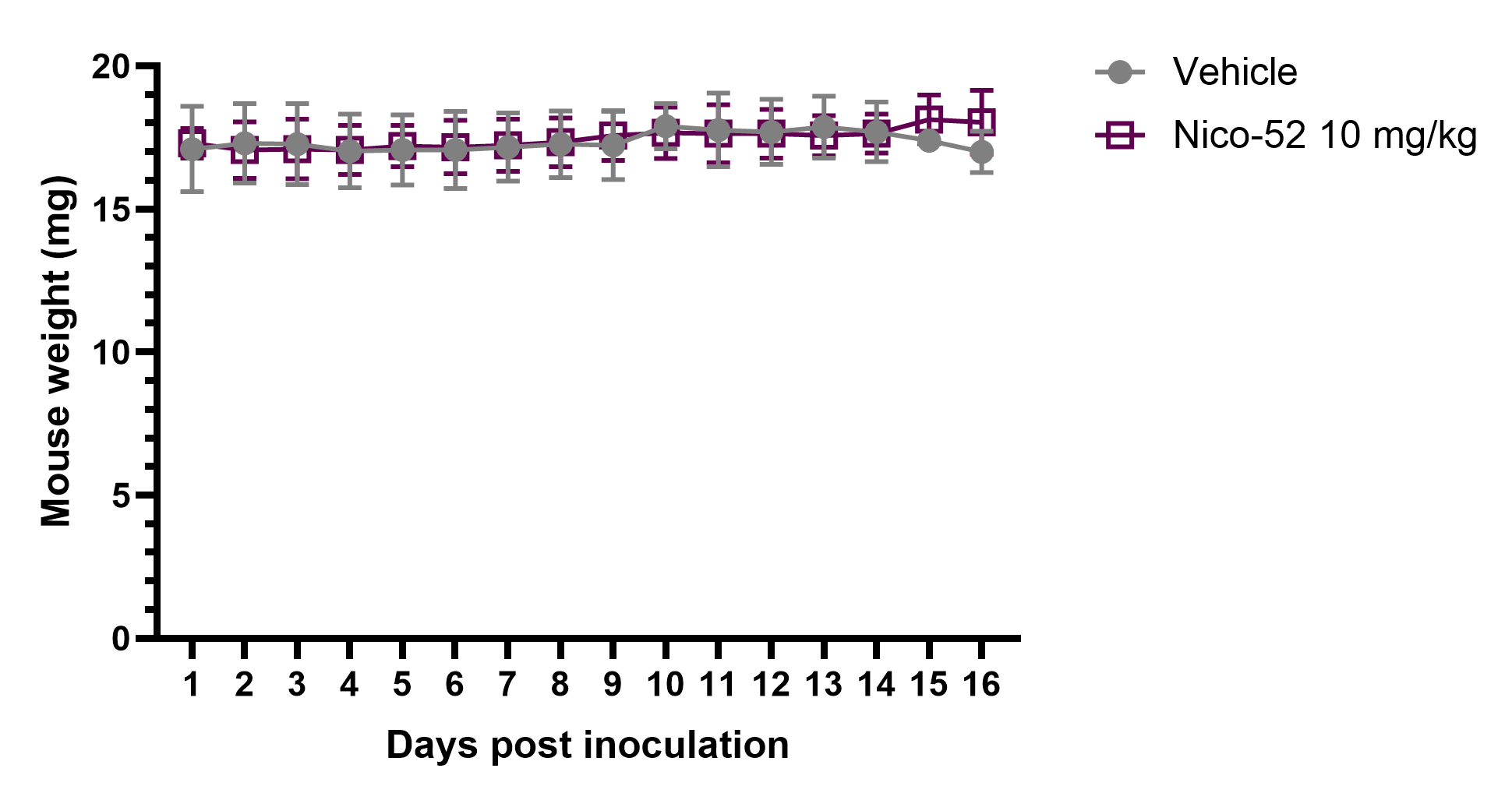

**Supplemental Figure 21. Flow cytometry gating of dissociated B16-F10 tumors 17 days post-inoculation.** FSC-H vs. SSC-H gating was performed. Then, positive events for both F4/80 and CD11b were identified as macrophages, further delineated by staining of CD80 (M1 associated) and CD206 (M2 Associated).

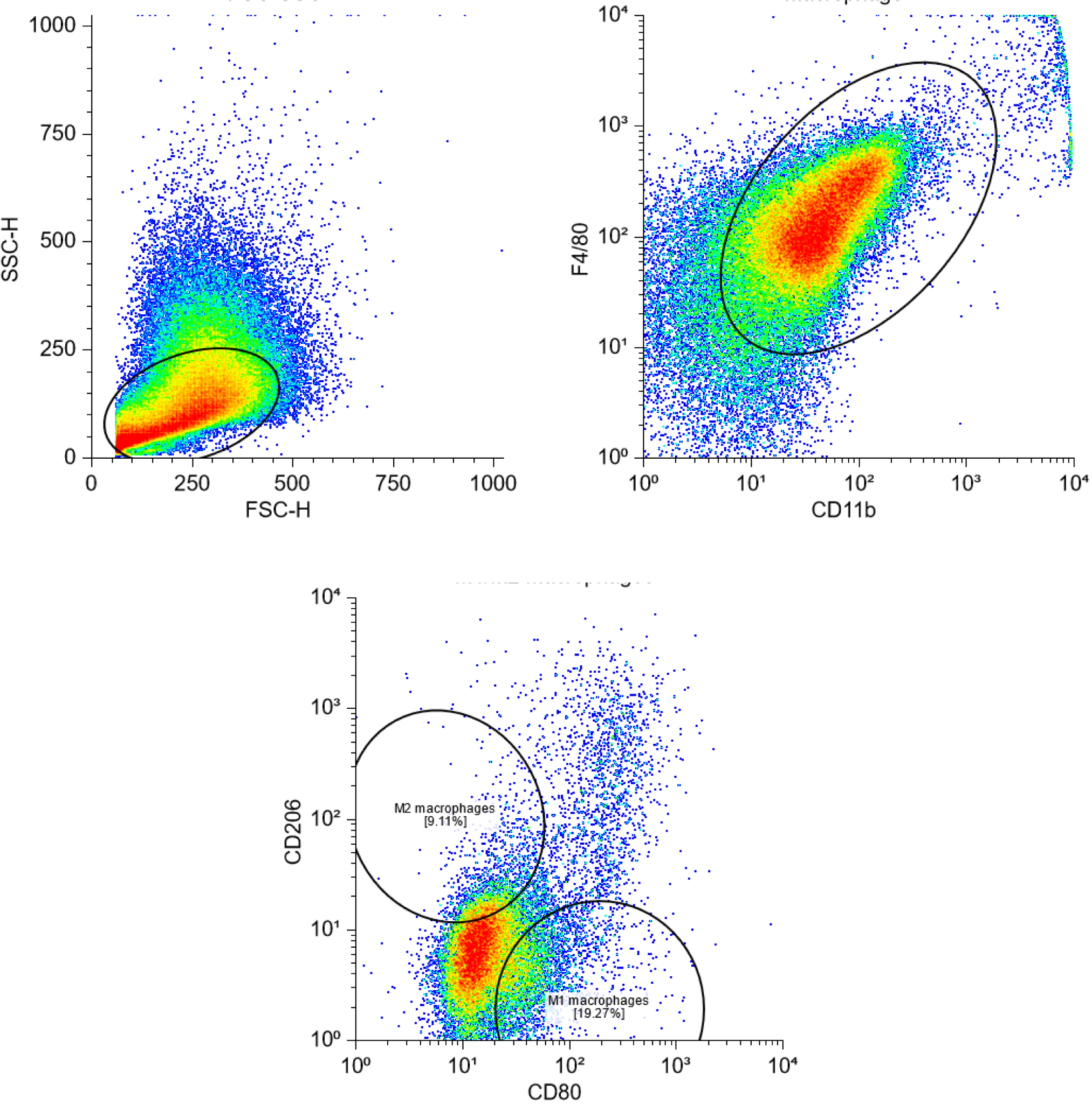

**Supplemental Figure 22. Survival study in 4T1 tumor-bearing Balb/C mice treated IP with Nico-52 at 10 mg/kg three times a week**

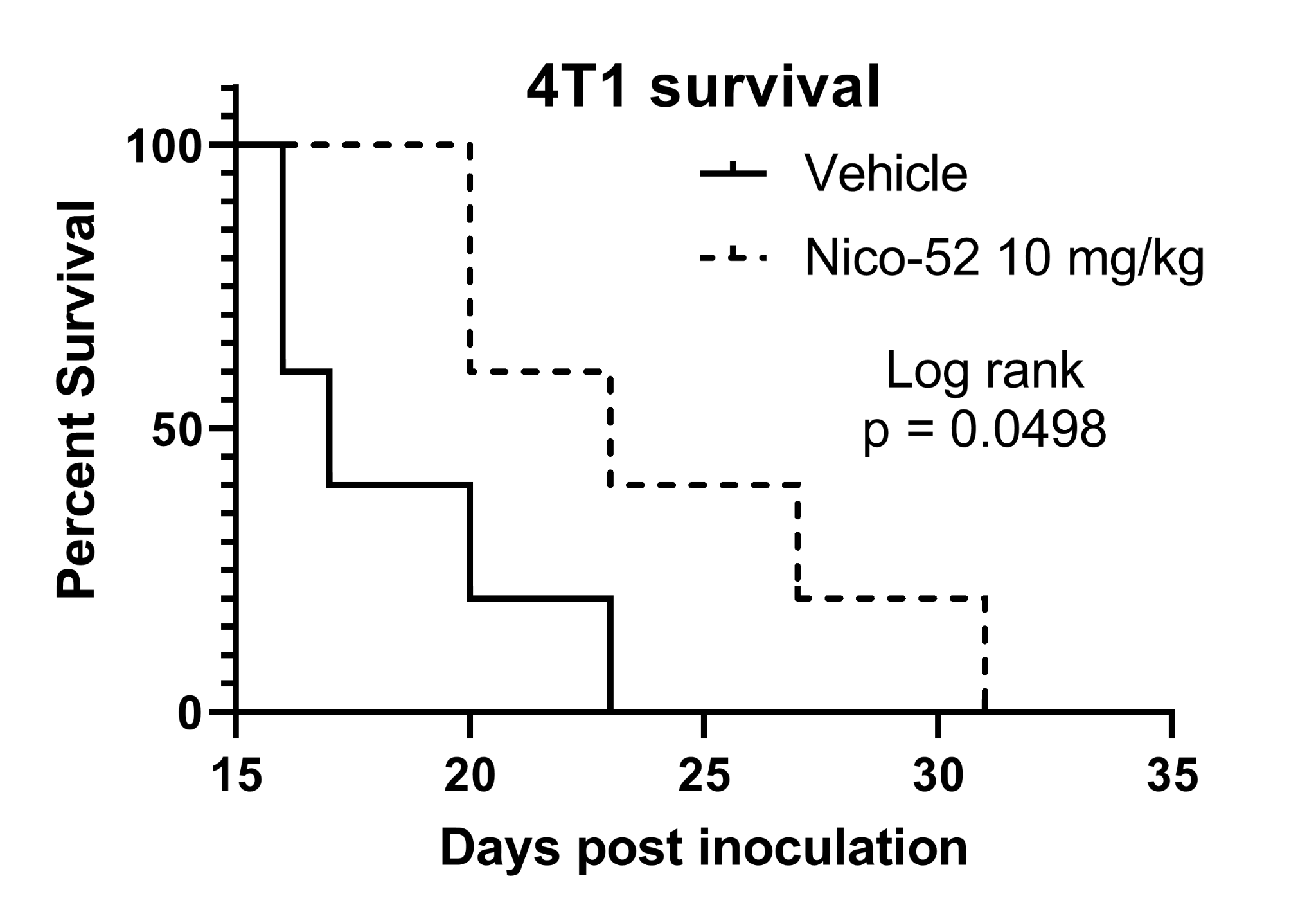

**Supplemental Figure 23.** **Nico-52 UPLC-MS absorbance AUC standard curve. R^2^ = 0.9976.**

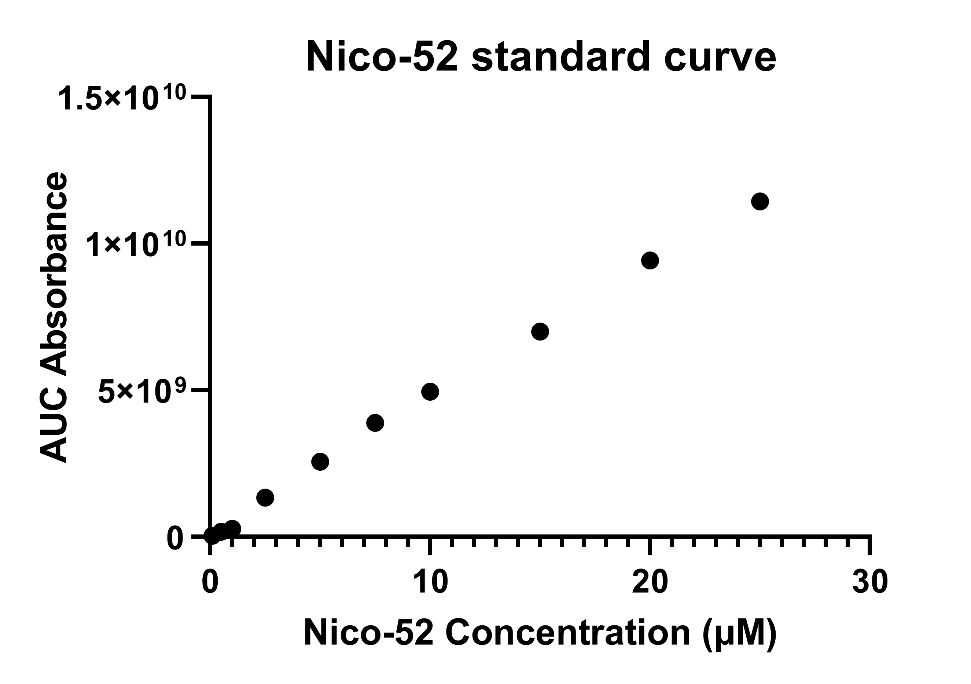

**HPLC traces of representative analogs:**

**Compound 1**

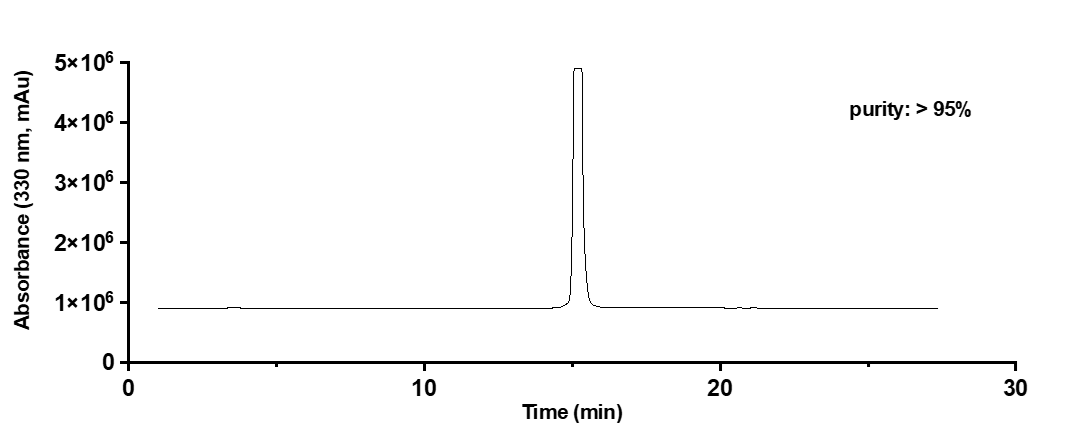

**Compound 2**

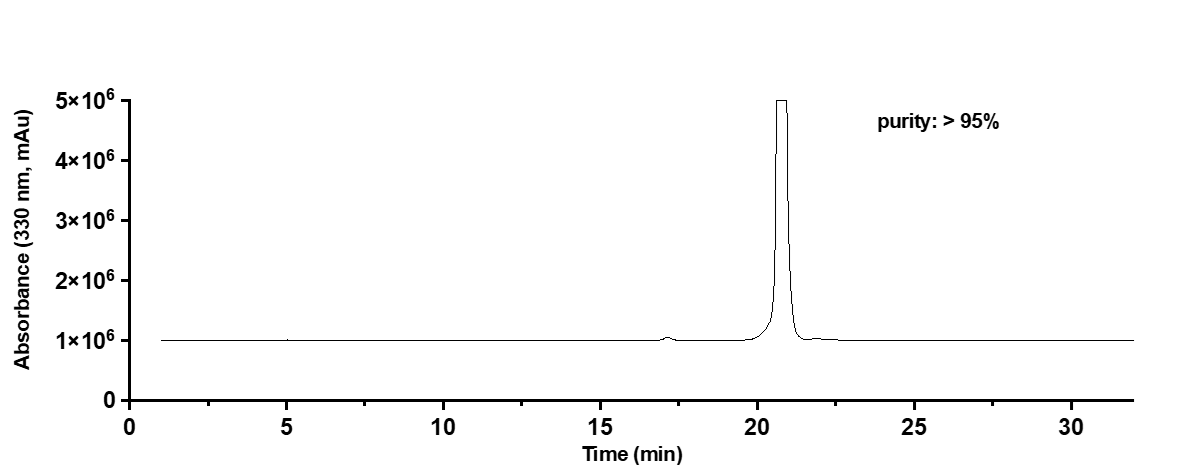

**Compound 10**

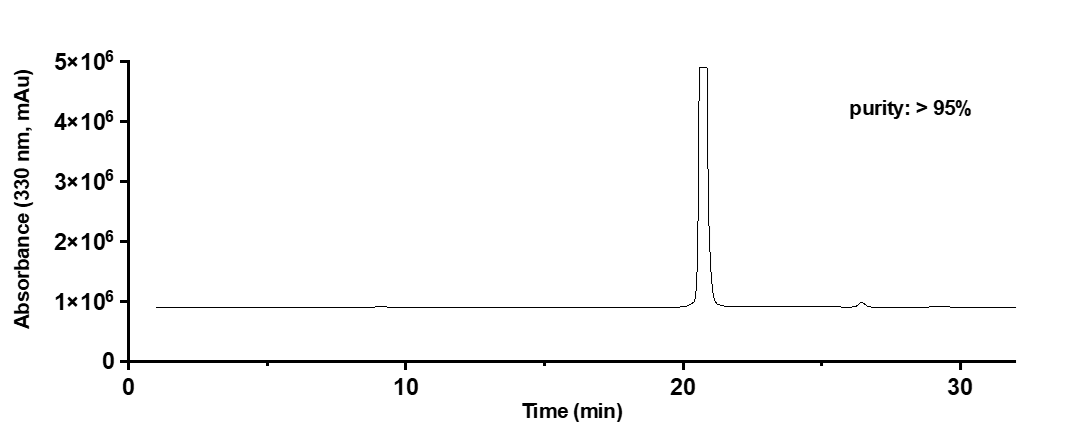

**Compound 11**

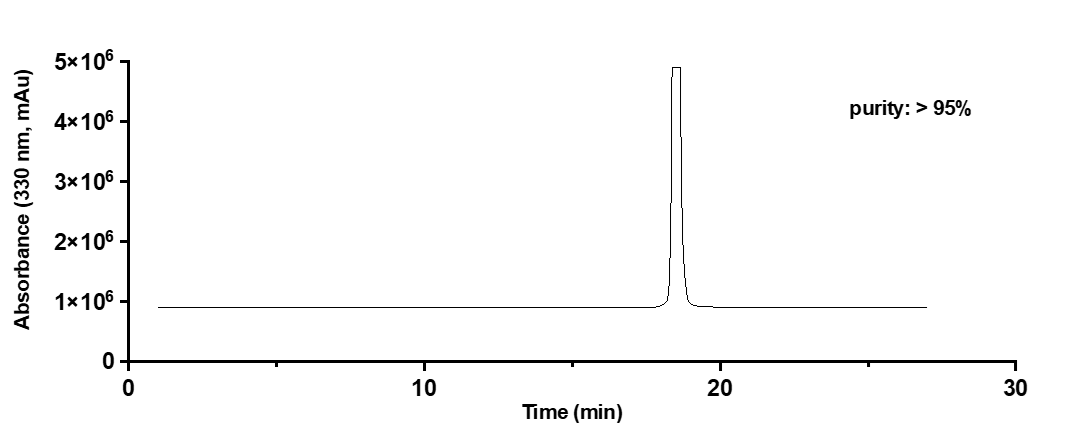

**Compound 12**

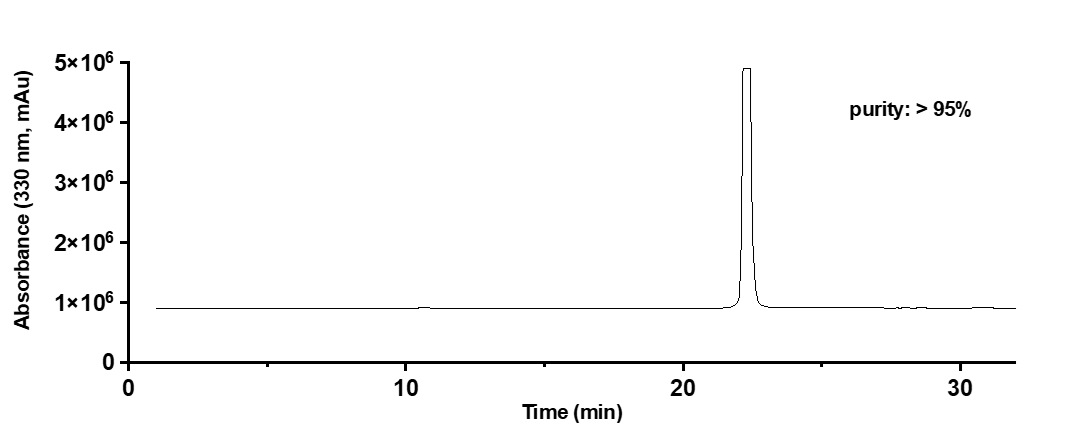

**Compound 43**

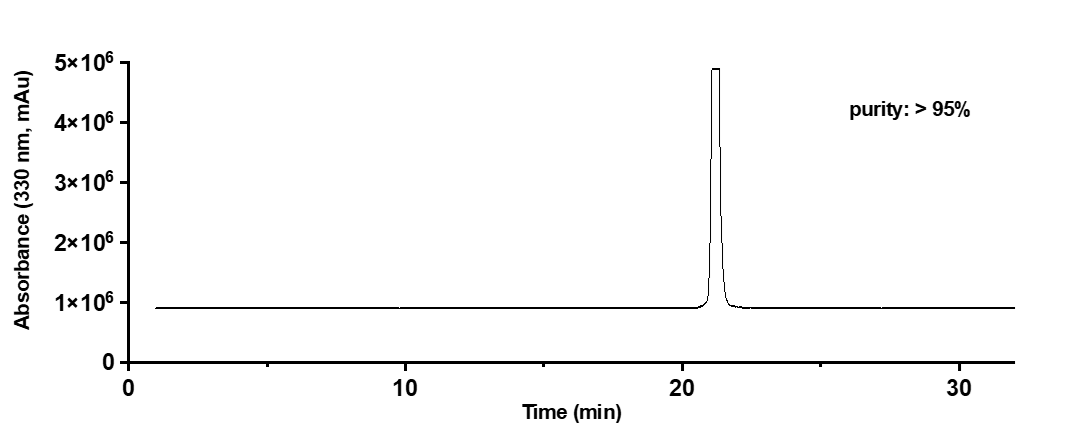

**LC-MS traces of representative analogs:**

**Compound 3: [M+H] ^+^ = 382 Da**

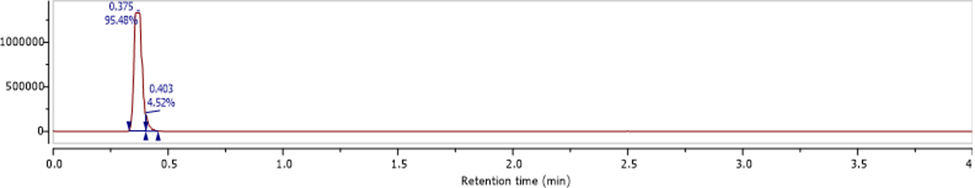

**Compound 6: [M+H] ^+^ = 334 Da**

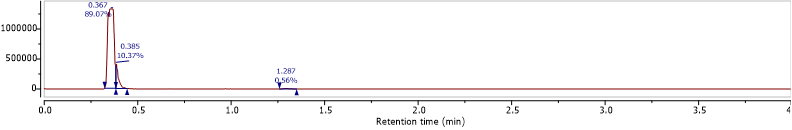

**Compound 8: [M+H] ^+^ = 294 Da**

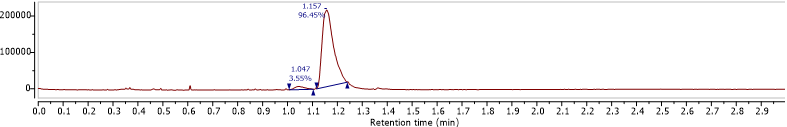

**Compound 9: [M+H] ^+^= 306 Da**

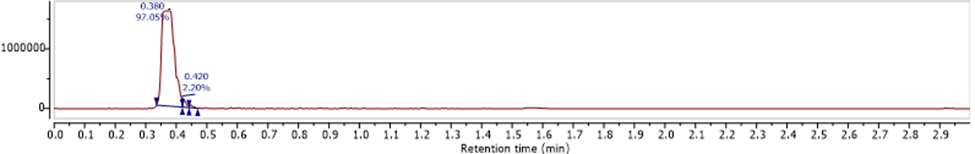

**Compound 13: [M+H] ^+^ = 242 Da**
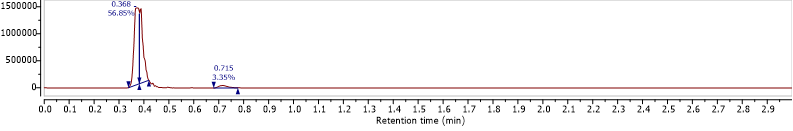
